# Kinetic design of reversible probe exchange enables continuous single-molecule tracking beyond the photobleaching limit

**DOI:** 10.64898/2026.03.13.711506

**Authors:** Kohei Iijima, Toshikuni Awazu, Shahi Imam Reja, Toshiyasu Sowa, Taketoshi Kambara, Masafumi Minoshima, Yasushi Okada, Kazuya Kikuchi

## Abstract

Single-molecule fluorescence imaging enables direct tracking of molecular dynamics. Prolonged observation, however, is limited by photobleaching, restricting access to rare events and slow transitions. Exchangeable fluorogenic labeling has been developed to overcome this limitation, yet continuous single-molecule tracking has not been achieved. We define two quantitative kinetic conditions for continuous tracking: pre-bleach dissociation and within-frame rebinding. Mapping reported systems onto a kinetic landscape delineates the regime required for continuous tracking. To reach this regime, we turned to odorant-binding proteins, whose biological role favors rapid ligand capture and release. We paired this scaffold with high-contrast fluorogenic probes to maximize rebinding capacity while preserving rapid dissociation. The resulting system, EverGreen, sustains uninterrupted single-molecule observation for over 24 hours and establishes a general kinetic design framework beyond the photobleaching limit.

## Introduction

Proteins populate multiple conformational states with distinct functional properties. Advances in single-particle cryo-electron microscopy have revealed extensive structural heterogeneity, identifying coexisting conformations within the same sample^1,2^. However, static structural ensembles do not establish the kinetic connectivity or temporal ordering between states^2,3^. Single-molecule fluorescence imaging can, in principle, track such dynamics in real time^4–7^. In practice, however, photobleaching limits observation duration. Considerable effort has focused on improving photostability through protein engineering and synthetic dye development^8–11^. These efforts extend observation time but do not eliminate photobleaching as the ultimate constraint on continuous single-molecule tracking.

Exchangeable fluorogenic labeling enables probes to reversibly bind and be replenished from solution. This strategy decouples signal persistence from photobleaching of individual fluorophores. Multiple exchangeable labeling systems have been developed based on this principle^12–19^. They have been applied to live-cell imaging and super-resolution microscopy^15–19^.

These exchangeable labeling systems perform robustly at the ensemble level. Probe replenishment sustains fluorescence even as individual probes photobleach. At the single-molecule level, the situation is different. Probe turnover can be directly resolved. Each probe is detected only for its residence time, and dissociation interrupts the signal^16,20^. This intermittency can be useful for localization-based super-resolution imaging, where transient binding events act as stochastic fluorescence switches^14–16,19^. For continuous single-molecule tracking, however, intermittency becomes a limitation. Dissociation-induced gaps fragment trajectories. A single trajectory is often confined to one binding event. Concatenating multiple tag copies can partially improve continuity^18^, but larger tags can perturb the dynamics under measurement. Thus, uninterrupted tracking beyond a single binding event remains challenging^20^.

Continuous single-molecule tracking through probe exchange requires two independent kinetic conditions. Newly arriving probes must bind rapidly enough to prevent resolvable gaps between successive binding events. Bound probes must dissociate before photobleaching destroys the fluorophore and damages the protein tag through reactive oxygen species generation^16,20–22^. Both conditions must be satisfied simultaneously. However, no existing exchangeable labeling system has, to our knowledge, been demonstrated to meet both requirements^13,15,16,20^. Achieving this regime by affinity tuning is often difficult. Association and dissociation can, in principle, be tuned independently, but affinity tuning often shifts both rates in a correlated manner within a given binding scaffold. Lowering affinity to accelerate dissociation often slows association, while tightening binding to speed association retards dissociation. This coupling makes it difficult to reach the regime in which both rates are simultaneously fast through incremental affinity tuning of a given protein–ligand pair. Kinetic design is therefore required to access this regime.

Here, we present EverGreen, an exchangeable fluorogenic labeling system designed to operate in this kinetic regime (Fig. 1). We formalize the two kinetic requirements as quantitative design criteria. We then map reported exchangeable labeling systems in this space. This analysis shows that the regime required for continuous single-molecule tracking has not been experimentally accessed. To reach this regime, we exploited odorant-binding proteins (OBPs) as a kinetic scaffold. OBPs are small soluble carriers whose biological role as olfactory transporters has imposed evolutionary pressure for both rapid ligand capture and rapid release^23,24^. This can represent a rare exception to the kinetic trade-off that constrains most protein–ligand systems. The diversity of the OBP family suggests that orthogonal ligand–protein pairs could be engineered, enabling potential multiplexing. We paired this scaffold with high-contrast fluorogenic probes to maximize rebinding capacity while preserving rapid dissociation. Probe turnover outpaces both photobleaching and the imaging frame rate. This enables continuous tracking of individual proteins for over 24 hours. EverGreen extends single-molecule observation beyond photobleaching limits, opening access to slow conformational dynamics and rare state transitions.

**Figure 1.**
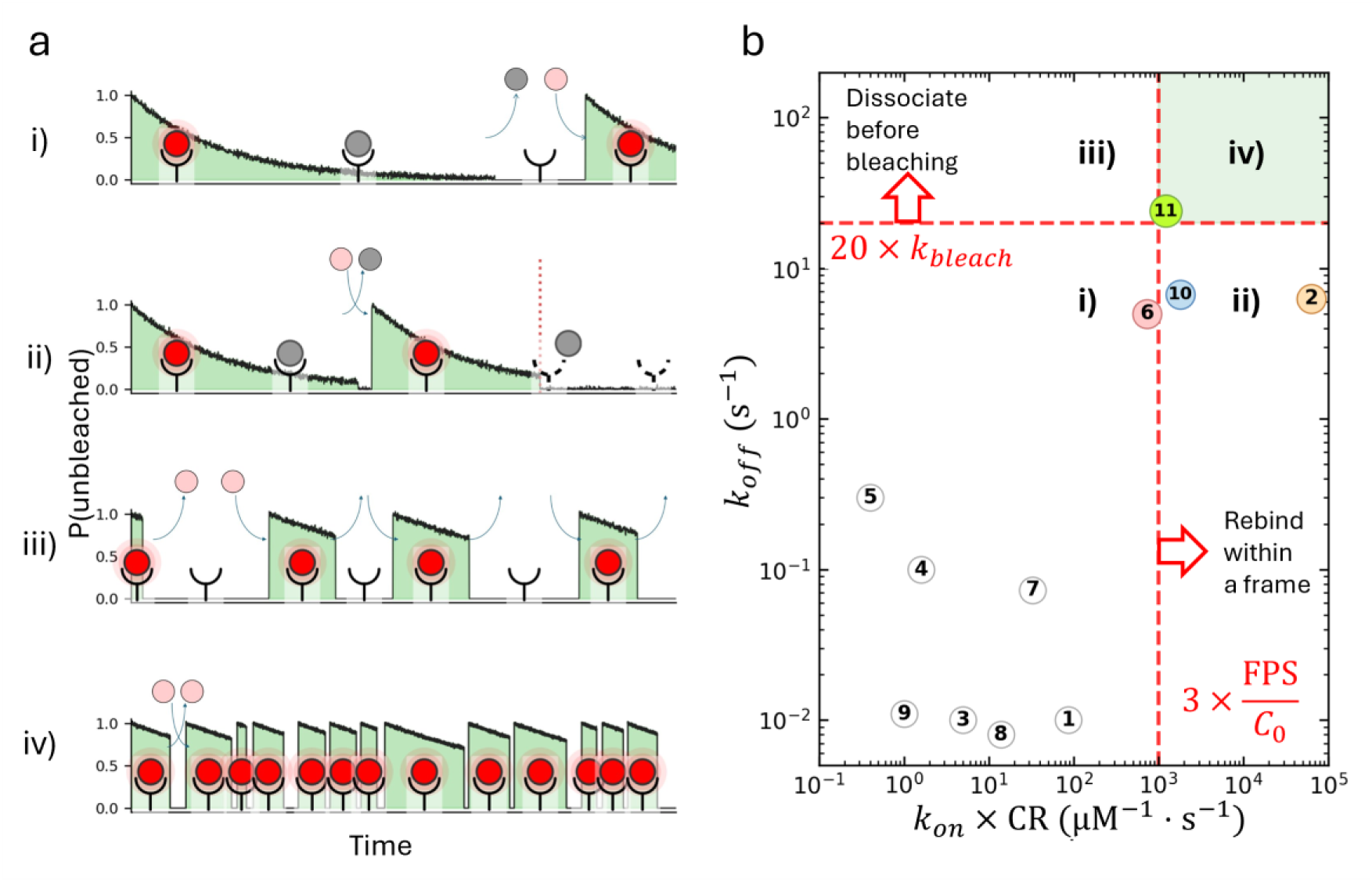
Kinetic requirements for continuous single-molecule tracking with exchangeable labeling systems. **a**, Four kinetic regimes (i–iv) for exchangeable fluorogenic labeling, showing the probability that a tag carries an unbleached probe. Only regime iv supports continuous single-molecule observation. **b**, Regime map of reported exchangeable labeling systems (Supplementary Table 1), plotted as *k*_*off*_versus *k*_*on*_ × *CR*. Quadrants i–iv correspond to the regimes in **a**. Dashed red lines mark design benchmarks (Supplementary Note 1); green shading denotes quadrant iv. Colored markers highlight systems discussed in the text: FAST (2), xHaloTag (6), SiR-tag (10), and EverGreen (11, this study).

## Results

### Quantitative criteria for continuous single-molecule tracking

In an exchangeable labeling system, probe molecules reversibly bind to a protein tag. When many tagged molecules contribute to the signal, stochastic turnover averages out and fluorescence appears continuous. In single-molecule imaging, however, each exchange event becomes directly observable. If the turnover kinetics are slow, probe binding and dissociation appear as distinct blinks; if sufficiently fast, dark intervals fall within individual imaging frames and are no longer resolved.

For this to hold, both ON and OFF kinetics must be fast (Fig. 1a). If a bound probe dissociates too slowly, it photobleaches while still on the tag. The fluorescence turns off, and the tag must wait for the bleached probe to dissociate before a new one can bind (Fig. 1a, i and ii). If, conversely, a new probe binds too slowly, the fluorescence turns off while the tag waits for the next probe to arrive (Fig. 1a, i and iii). Crucially, when photobleaching occurs while the probe is bound, the resulting reactive oxygen species can damage the tag protein itself, irreversibly destroying its ability to bind new probes^16,20,22^ (Fig. 1a, ii).

These considerations lead to two quantitative design criteria. First, each probe molecule should dissociate before photobleaching occurs:

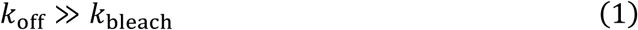

A competing first-order analysis (Supplementary Note 1) shows that pre-bleach dissociation with > 95% probability requires *k*_off_ ≳ 20 × *k*_bleach_. Given typical single-molecule photobleaching rates on the order of 1 s^−1^ or below^10^, this corresponds to *k*_off_ ≳ 20 s^−1^.

Second, a new probe must bind before the next imaging frame. The rebinding rate is the product of the association rate constant *k*_on_ and the free probe concentration, which is limited by the requirement that background fluorescence from unbound probes remain well below the single-molecule signal. For a fluorogenic system with contrast ratio CR, the maximum usable probe concentration scales with CR × *C*_0_, where *C*_0_ is the characteristic concentration scale for single-molecule detection, determined by the effective observation volume and detection sensitivity (≈ 30 nM for typical TIRF geometries; see Supplementary Note 1 for derivation). Requiring rebinding within a single frame gives:

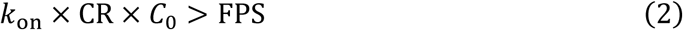

where FPS is the imaging frame rate (10 Hz in this study). The product *k*_on_ × CR serves as a figure of merit for the rebinding capacity of the labeling system, analogous to ε × Φ for fluorophore brightness.

These inequalities represent probabilistic design criteria rather than sharp physical thresholds. The dissociation condition ensures that pre-bleach dissociation dominates over bound-state photobleaching as a competing first-order process, whereas the rebinding condition ensures that the waiting time for a new probe is shorter than the camera frame interval with high probability. A detailed probabilistic analysis (Supplementary Note 1) shows that achieving within-frame rebinding with > 95% probability requires approximately a threefold kinetic margin, *k*_on_ × CR ≳ 3 × FPS/*C*_0_, corresponding to *k*_on_ × CR ≳ 1,000 μM^−1^ s^−1^ at 10 Hz. Together, these criteria define a two-dimensional kinetic design space for continuous single-molecule tracking.

Plotting *k*_off_against *k*_on_ × CR maps the four regimes onto a kinetic landscape. Because the underlying waiting times are stochastic, we define these regime boundaries using practical design margins rather than sharp thresholds (*k*_off_ ≳ 20 *k*_bleach_and *k*_on_ × CR ≳ 3 FPS/*C*_0_; Fig. 1b; see Supplementary Note 1 for derivation). We compiled kinetic data for reported exchangeable labeling systems and plotted them on this map (Supplementary Table 1). Several systems with *k*_on_approaching the diffusion limit have been reported, including FAST (2 in Fig. 1b)^13^, xHaloTag (6)^15^, and SiR-tag (10)^19^, and applied to single-molecule observation. FAST, in particular, achieves a rebinding capacity (*k*_on_ × CR) well above the threshold for within-frame rebinding. However, its *k*_off_remains below *k*_bleach_, and detailed photochemical studies have demonstrated that reactive oxygen species generated during bound-state photobleaching cause irreversible oxidative damage to the tag protein^22^, directly confirming the consequence predicted for regime ii. Continuous single-molecule tracking has not been achieved with any of these systems^20^. To our knowledge, no reported system has been experimentally demonstrated to occupy regime iv under sustained single-molecule imaging conditions. The goal of this work is to develop a system that enters this regime.

### Design of EverGreen protein labeling system

Because association and dissociation rates are often kinetically coupled, satisfying both conditions simultaneously by conventional protein engineering alone is challenging. Rather than engineering an existing tag to move toward regime iv, we sought a protein scaffold whose natural function already demands both rapid ligand capture and rapid release.

Odorant-binding proteins (OBPs) are small (19 kDa) soluble carriers that bind hydrophobic odorant molecules and transport them through the aqueous mucus layer to olfactory receptors^23–25^. Efficient olfactory sensing requires that OBPs rapidly capture dilute odorants from the mucus and subsequently release them to receptors on a timescale compatible with sensory processing. This biological role imposes evolutionary pressure on both association and dissociation kinetics, representing a rare exception to the empirical trade-off typically observed in protein–ligand systems. We therefore reasoned that an OBP-based tag may be able to access the kinetic regime required for continuous single-molecule tracking.

Even with fast association and dissociation, the rebinding condition (Eq. 2) requires that the product *k*_on_ × CR exceed the threshold. Because OBP binds its ligands with moderate affinity (micromolar *K*_d_), high probe concentrations are needed for rapid rebinding, which in turn demands a large fluorogenic contrast ratio to suppress background. We therefore designed fluorogenic probes based on conjugates of 8-dimethylamino-benzo[*g*]coumarin (DMABC) dye^26^ and linear aliphatic carboxylic acids (Fig. 2a). DMABC exhibits a non-fluorescent state in polar solvents owing to the preferential formation of a non-emissive twisted intramolecular charge-transfer (TICT) state, and emits green fluorescence in low-polarity environments^26^. To mimic fatty acid-based odorant molecules, linear aliphatic carboxylic acids with different alkyl chains were conjugated to the DMABC dye. A series of fluorogenic EverGreen probes (DMABC-FA6, –FA8, –FA10, and –FA12) were synthesized by attaching medium-chain carboxylic acids, such as hexanoic, octanoic, decanoic, and dodecanoic acids, which served as non-covalent OBP ligands with moderate binding affinities (Scheme S1).

**Figure 2.**
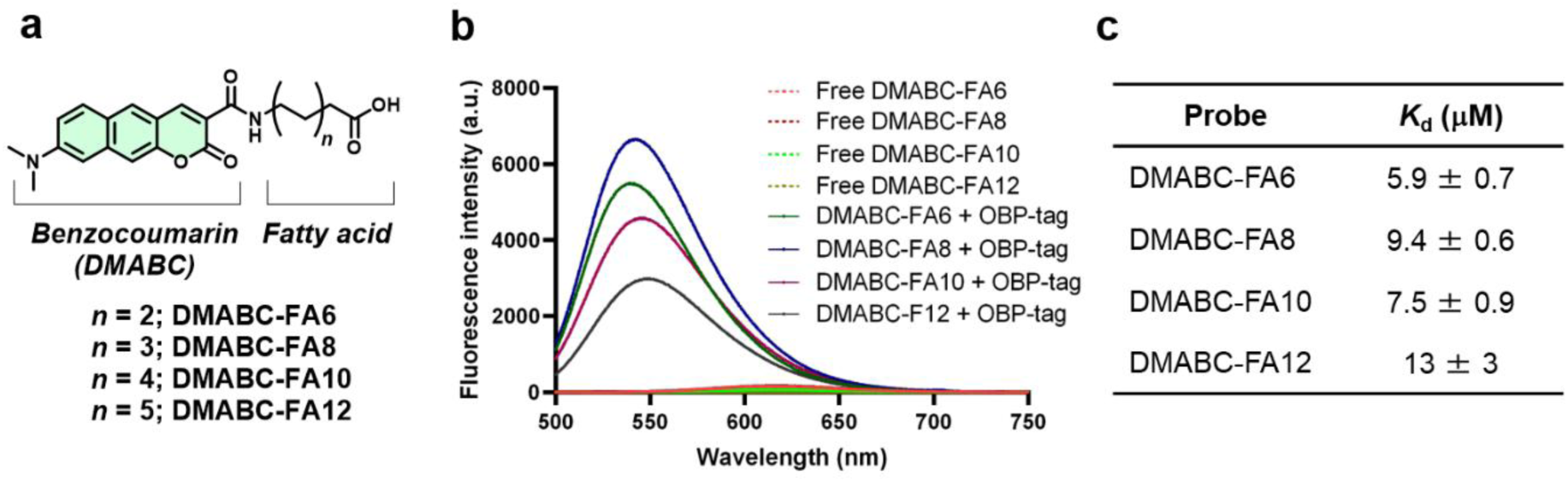
EverGreen probe design and labeling properties. **a.** Chemical structures of EverGreen probes based on DMABC fluorophore. **b.** Fluorescence spectra of DMABC-FA probes (1.0 μM, 0.2% DMSO) in the absence and presence of OBP-tag (50 μM) in 20 mM HEPES buffer (pH 7.4), 150 mM NaCl at 37 °C. The excitation wavelengths for DMABC-FAs were 467 nm (DMABC-FA6, DMABC-FA8, and DMABC-FA10) and 465 nm (DMABC-FA12). **c.** *K*_*d*_ values of DMABC probes with OBP-tag.

We term the resulting combination of an OBP-derived tag and these fluorogenic probes EverGreen. Whether this system satisfies both kinetic criteria and occupies the continuous-tracking regime was tested by the quantitative measurements described below.

### Fluorogenic response of EverGreen probes for OBP-tag

We first examined the fluorogenic response of the EverGreen probes upon binding to the OBP-tag. Upon incubation with recombinant human OBP2a proteins^25^ (C99S K112N mutant, hereafter referred to as OBP-tag), the EverGreen probes exhibited a substantial (over 350-fold) increase in fluorescence intensity accompanied by an approximately 70 nm hypsochromic shift (Fig. 2b, Supplementary Fig. 1a). These fluorogenic responses indicate that the DMABC fluorophores are positioned within a hydrophobic environment, corresponding to the polarity-sensitive characteristics in a series of organic solvents (Supplementary Fig. 1b). In contrast, the DMABC dye without fatty acids showed less fluorescence enhancement, underscoring the important role of this moiety in fluorogenic response (Supplementary Fig. 1c). The fluorescent probe-protein complexes remained photostable under continuous excitation for up to 30 min (Supplementary Fig. 1d). The binding affinities between the EverGreen probes and the OBP-tag were measured by fluorescence titration. EverGreen probes bound to OBP-tag with micromolar dissociation constants (*K*_d_ = 5.9–13 μM), slightly higher than those reported for natural fatty acids^25^ (Fig. 2c, Supplementary Fig. 1e). These results indicate that the EverGreen exhibits large fluorogenic responses and moderate binding affinity as a protein labeling system.

### Single-molecule imaging with a kinesin-OBP model

To evaluate the applicability of EverGreen at the single-molecule level, we used the motor protein kinesin^27^ as a model system. Kinesin is one of the best-characterized molecular motors in single-molecule biophysics^28^, and well-established assay conditions allow flexible control of its functional state on microtubules—including processive movement, stationary binding, and defined oligomeric states—making it an ideal test bed for validating a new labeling system at the single-molecule level^5^.

To test whether the OBP-tag functions as a fusion tag, we fused it to the C-terminus of motor-domain constructs of murine kinesin KIF5C^5,29^. Kinesin provides a convenient readout. Microtubule-bound signals are readily distinguished from nonspecific surface adsorption of the fluorogenic probes. Specific binding to microtubules and ATP-driven motility report on the functional integrity of the fusion construct.

We first imaged a monomeric construct K351–OBP (residues 1–351) bound to microtubules in a nonmotile rigor binding state using the non-hydrolyzable ATP analog AMP–PNP^29^. At 300 nM K351–OBP, microtubules were uniformly labeled, and the EverGreen signal showed strong colocalization with TMR-labeled microtubules in a separate detection channel, confirming specific labeling (Supplementary Fig. 2). At 10 pM K351–OBP, fluorescent spots were observed along the microtubules at a density consistent with the kinesin concentration (Fig. 3b). Their intensity distribution confirmed single-molecule detection (Supplementary Note 2, Supplementary Fig. 3).

**Figure 3.**
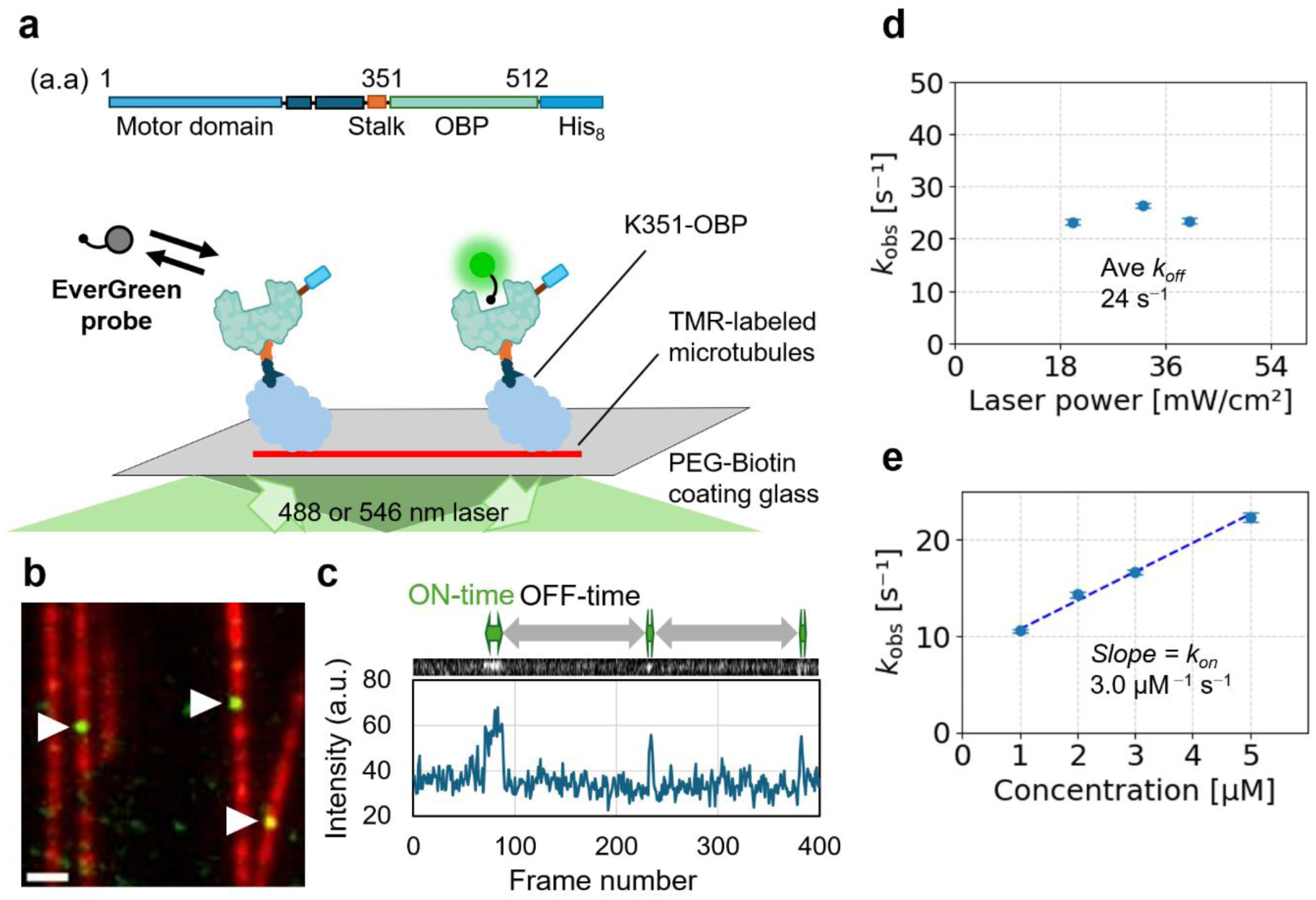
Single-molecule kinetic analysis of EverGreen probe exchange. **a.** Schematic representation of the K351–OBP construct and experimental setup for single-molecule fluorescence imaging. A monomeric kinesin construct (K351) was fused at its C-terminus to the OBP-tag and imaged on surface-immobilized microtubules using total internal reflection fluorescence microscopy (TIRFM). **b.** Representative EverGreen fluorescence signals were observed along the microtubules (white arrowheads, left; scale bar, 1 μm). **c.** Kymograph and corresponding fluorescence intensity time trace obtained by continuous streaming acquisition at the same spatial position, illustrating the stochastic transitions between fluorescent and non-fluorescent states. **d.** Comparison of dissociation rate constants (*k*_*off*_) determined at different excitation light intensities. The extracted *k*_*obs*_ values were insensitive to the excitation power, indicating that probe dissociation was not dominated by photophysical processes. **e.** Observed reappearance rate (*k*_*obs*_) as a function of the probe (DMABC-FA8) concentration. The slope of the plot indicates the association rate constant (*k*_*on*_). Error bars in d and e represent 90% confidence intervals obtained by bootstrap resampling (N_boot = 1000).

We next examined a dimeric construct K560–OBP (residues 1–560) in the presence of ATP. The characteristic unidirectional movement of kinesin along microtubules was clearly observed, with a velocity consistent with previous reports^5^ (Supplementary Fig. 4), confirming that the OBP-tag does not affect motor function. Together, these results establish that EverGreen provides specific labeling and sufficient signal quality for single-molecule detection without compromising the activity of the fusion partner.

### Single-molecule kinetics of EverGreen

To quantify the binding and dissociation kinetics of EverGreen, we analyzed fluorescence time traces of individual K351–OBP molecules immobilized on microtubules (Fig. 3c). Stochastic transitions between fluorescent (ON) and non-fluorescent (OFF) states were identified, and the dwell times of each state were extracted for kinetic analysis (Supplementary Note 3).

The ON-dwell time distribution was well described by a single exponential (Supplementary Fig. 5a and 6a). Because the observed rate is the sum of two competing first-order processes, *k*_obs_ = *k*_off_ + *k*_bleach_, and *k*_bleach_ scales with excitation power density, any substantial contribution from photobleaching would manifest as a power-dependent increase in *k*_obs_. Over a twofold range of excitation power density, *k*_obs_ showed no marked dependence on excitation power density (Fig. 3d, Supplementary Fig. 5b, 6b), indicating that *k*_off_dominates over *k*_bleach_and that the measured ON-dwell time of 41 ms largely reflects intrinsic probe dissociation.

The OFF-dwell time distribution was also well approximated by a single exponential (Supplementary Fig. 5c, 6c). The observed reappearance rate increased linearly with probe concentration (Fig. 3e, Supplementary Fig. 5d, 6d), yielding an association rate constant of *k*_on_ = 3.0 μM^−1^s^−1^. These results show that fluorescence switching in EverGreen is governed by simple binding and dissociation kinetics. The dissociation constants calculated from the single-molecule kinetic parameters (*K*_d_ = *k*_off_/*k*_on_) are consistent with those measured by bulk fluorescence titration, confirming the internal consistency of the kinetic analysis.

These kinetic parameters place EverGreen at *k*_on_ × CR ≈ 1,050 μM^−1^s^−1^ and *k*_off_ ≈ 24 s^−1^ (11 in Fig. 1b), exceeding the 95%-probability design thresholds for both within-frame rebinding (Eq. 2) and pre-bleach dissociation (Eq. 1). The dissociation rate is the fastest among reported exchangeable labeling systems, yet the rebinding capacity remains above the threshold (Supplementary Table 1). EverGreen thus occupies regime iv, the first experimental entry into the continuous-tracking regime.

### Long-term single-molecule fluorescence imaging with EverGreen

The kinetic measurements above place EverGreen within regime iv, predicting that at 10 μM DMABC-FA8 and 10 Hz frame rate, probe exchange should in principle sustain fluorescence without a photobleaching-imposed time limit. To test this prediction, we performed continuous single-molecule imaging of K351–OBP immobilized on glass substrates. K351–OBP was tethered via biotinylated anti-His-tag antibodies and avidin, because in vitro microtubule assays have limited stability over extended observation periods. An enzymatic oxygen-scavenging system (GLOX) was also included to suppress potential cumulative photodamage from reactive oxygen species generated by the high concentration of free probes under sustained illumination. Under these conditions, individual fluorescent spots were indeed continuously detected throughout 24 h of imaging, corresponding to approximately one million consecutive frames, with no detectable loss of fluorescence intensity (Fig. 4a, 4b, Supplementary Movie 1). The observation duration in this experiment was limited not by signal degradation but by practical considerations such as probe supply and stage drift correction.

**Figure 4.**
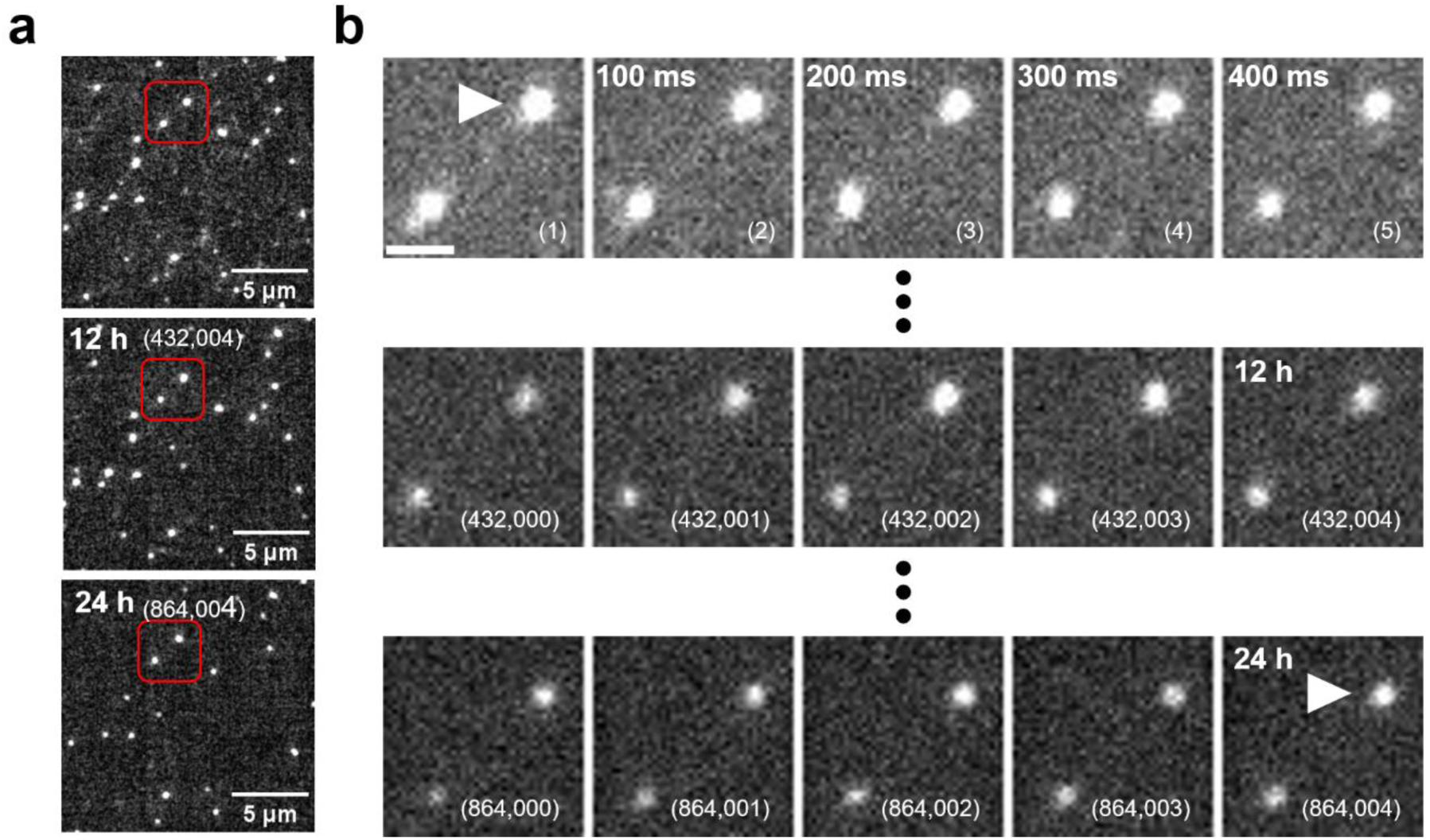
Long-term single-molecule imaging enabled by EverGreen. **a.** Wide-field views from continuous single-molecule fluorescence imaging of K351–OBP immobilized via biotinylated anti-His-tag antibodies and avidin, acquired at the start of imaging, after 12 h, and after 24 h during continuous acquisition at 10 Hz with 10 μM DMABC-FA8 under TIRFM. The numbers in parentheses indicate the corresponding frame numbers. The red boxes denote the same regions across time points. **b.** Magnified view of the region outlined in red in **a**. The white arrowhead indicates the same individual molecules observed throughout the experiment. Scale bar, 1 μm.

To confirm the probe-exchange mechanism, the same spot was imaged at 100 Hz. The fluorescence signal exhibited discontinuous behavior, alternating between ON and OFF states with kinetics consistent with the probe exchange kinetics described above (Supplementary Fig. 7), confirming that the sustained signal at 10 Hz originates from repeated cycles of probe dissociation and rebinding rather than from static, long-lived fluorescent contaminants. Conversely, this controlled blinking pattern provides an intrinsic kinetic fingerprint: because the photon budget no longer limits observation, high-speed acquisition can be performed at any point during a long experiment to verify that a fluorescent spot represents a labeled target molecule.

To assess whether EverGreen can sustain single-molecule signals without photochemical stabilization, we further performed an analogous experiment in the absence of the oxygen-scavenging system. Continuous single-molecule imaging at 10 Hz was maintained for at least one hour (36,000 consecutive frames) with minimal apparent signal degradation (Supplementary Fig. 8). This result indicates that rapid probe exchange alone is sufficient to sustain single-molecule fluorescence over timescales typical of most imaging experiments, broadening the applicability of EverGreen to conditions where oxygen-scavenging systems are impractical, including live-cell environments.

### Live-cell fluorescence imaging using EverGreen

As discussed above, long-term continuous observation of single molecules is now within reach. A natural next step would be to extend this capability to live cells. For intracellular applications, the probes must first permeate the cell membrane and reach their target proteins. Membrane permeability alone, however, is not sufficient. The probes must also retain labeling specificity within the complex intracellular environment. This is particularly challenging for environment-sensitive fluorogenic dyes. Hydrophobic fluorogens can partition into lipid membranes and other intracellular hydrophobic compartments rather than binding to their target. Even subtle structural differences can result in nonspecific membrane staining^30^. We therefore first assessed the ensemble-level labeling performance of EverGreen within live cells.

HEK293T cells expressing OBP-tagged proteins were labeled with EverGreen probes and imaged by confocal microscopy without washing. Bright fluorescence was observed in OBP-expressing cells, whereas non-transfected cells showed negligible signal (Fig. 5a). Signal-to-background ratios exceeded 10-fold for DMABC-FA6, –FA8, and –FA10 (Supplementary Fig. 9b, 9d). DMABC-FA12, bearing a longer dodecanoic chain, showed a reduced ratio owing to nonspecific accumulation from its higher lipophilicity. Fluorescence was localized specifically to the compartments where the OBP-tag was expressed, including the nucleus and mitochondria (Fig. 5b, Supplementary Fig. 9c, 9e–g). Based on these results, we selected DMABC-FA6 and –FA8 as optimal probes for subsequent experiments.

**Figure 5.**
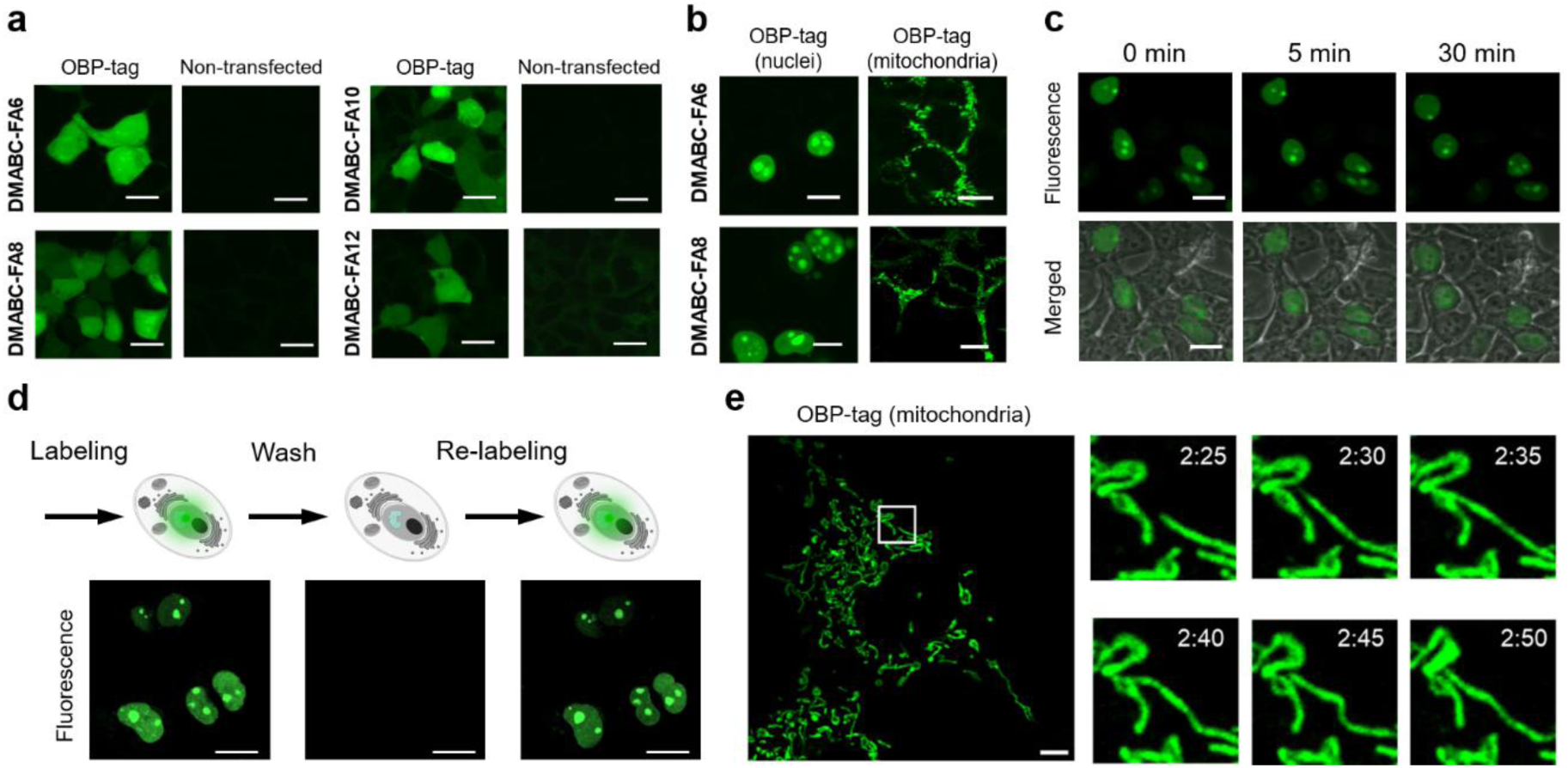
Live-cell fluorescence imaging using EverGreen. **a.** Live-cell fluorescence images of HEK293T cells expressing the OBP-tag or non-transfected cells after labeling with DMABC-FAs probes (λ_*ex*_ = 473 nm; λ_*em*_: 490–590 nm; scale bars: 20 μm). **b.** Fluorescence images of live HEK293T cells expressing OBP-tag in the nucleus and mitochondria after labeling with DMABC-FA6 and DMABC-FA8 (λ_*ex*_= 473 nm; λ_*em*_: 490–590 nm; scale bars: 20 μm). **c.** Time-lapse imaging of DMABC-FA8 in HEK293T cells expressing OBP-tag in the nucleus. The 0 min image was captured immediately after probe addition (λ_*ex*_ = 473 nm; λ_*em*_: 490–590 nm; scale bars: 20 μm). **d.** Schematic representation of repetitive labeling and fluorescence images of live COS-7 cells expressing OBP-tag in the nucleus after repetitive labeling with DMABC-FA6 (λ_*ex*_= 488 nm; scale bars: 20 μm). **e.** NSPARC imaging of DMABC-FA6 (1.0 μM) in COS-7 cells expressing OBP-tag in mitochondria, showing time-dependent changes in mitochondrial dynamics (λ_*ex*_= 488 nm), scale bar: 5 μm.

Fluorescence signals appeared immediately upon probe addition (Fig. 5c), indicating rapid labeling in cells. To test reversibility, cells were subjected to repeated cycles of labeling and washing. Signals disappeared after washing and reappeared upon re-addition of probes (Fig. 5d), confirming the non-covalent and reversible nature of EverGreen labeling in live cells. Time-lapse imaging showed that signals were maintained for up to 6 h, albeit with gradual decrease (Supplementary Fig. 9h). We further applied EverGreen to visualize mitochondrial dynamics using confocal microscopy with NSPARC super-resolution detection (Fig. 5e). Time-lapse imaging at 5 s intervals over 15 min resolved morphological changes, extension, and contacts of mitochondria in live cells.

These results demonstrate that EverGreen enables specific, rapid, and reversible labeling within live cells. As with other exchangeable labeling systems^13–19^, the renewable nature of the fluorescent signal provides sufficient contrast for wash-free ensemble imaging, including photobleaching-resistant time-lapse and super-resolution applications. Extending these capabilities to single-molecule tracking within live cells will require further optimization to control nonspecific probe partitioning, and represents an important next objective.

### Expansion to multicolor imaging using orthogonal pairs of fluorogenic ligands and OBP variants

Another natural extension of EverGreen is multicolor imaging. Because OBP accommodates a broad range of hydrophobic ligands, replacing the fluorophore moiety of the probe readily yields spectrally distinct variants. However, spectral diversity alone does not enable simultaneous multicolor imaging. Different probes sharing the same tag would compete for the same binding pocket, precluding independent labeling of distinct targets. True multicolor capability therefore requires orthogonal tag–probe pairs, in which each probe binds selectively to its cognate tag with minimal cross-reactivity. We explored this possibility by developing spectrally distinct probes and an engineered OBP variant with altered ligand selectivity.

We designed blue– and red-emitting fluorogenic probes based on 7-(dimethylamino)coumarin (DMAC)^31^ and 7-(6-(diethylamino)benzofuran-2-yl)benzo[*c*][1,2,5]thiadiazole (DMABT)^32^, respectively, selected for their polarity-sensitive fluorescence properties. As with EverGreen probes, linear aliphatic carboxylic acids were conjugated to each fluorophore (DMAC-FA and DMABT-FA; Fig. 6a, Schemes S2 and S3). These probes showed increased fluorescence with a hypsochromic shift in the presence of OBP-tag (Fig. 6b, 6c, Supplementary Fig. 10a, 11a), with DMAC-FA10 and DMABT-FA2 exhibiting the best fluorogenic responses (100-fold and 40-fold increase, respectively). These probes also showed sufficient photostability (Supplementary Fig. 10d, 11c) and micromolar binding affinities comparable to EverGreen probes (Supplementary Fig. 10e, 11d, Supplementary Tables 3 and 4). In live cells, DMAC– and DMABT-based probes specifically labeled OBP-tagged proteins (Fig. 6d, 6e; Supplementary Fig. 12a–d). Sequential exchange of blue (DMAC-FA10), green (DMABC-FA6), and red (DMABT-FA2) probes on the same OBP-tagged protein demonstrated color switching without significant cross-talk between cycles (Fig. 6f).

**Figure 6.**
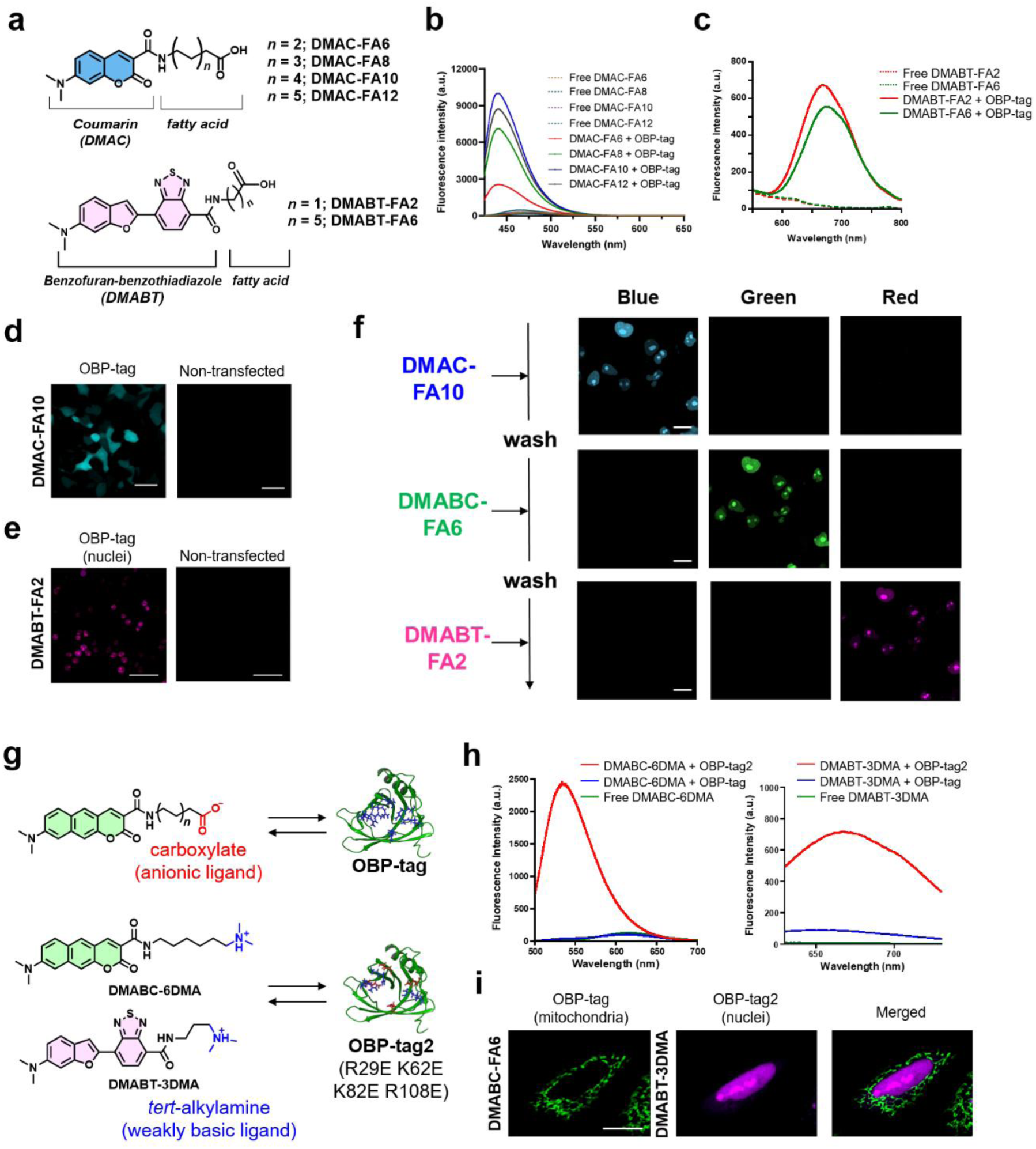
Multicolor imaging platform with EverGreen. **a.** Chemical structures of blue and red-fluorogenic probes for OBP-tag. **b,c.** Fluorescence spectra of DMAC-FA and DMABT-FA probes in the presence of OBP-tag. The excitation wavelengths for the DMAC-FA and DMABT-FA probes were 413 and 530 nm, respectively. **d,e.** Live cell fluorescence imaging of HEK293T cells expressing OBP-tag in nuclei after staining with DMAC-FA10 and DMABT-FA2. The excitation wavelengths for the DMAC-FA and DMABT-FA probes were 405 nm and 635 nm, respectively. Scale bar: 40 μm. **f.** Live cell fluorescence imaging of COS-7 cells expressing OBP-tag in nuclei after sequential exchange of blue, green, and red fluorogenic probes. The excitation wavelengths for DMAC-FA10, DMABC-FA6, and DMABT-FA2 were 405, 488, and 638 nm, respectively. Scale bar: 20 μm. **g.** Chemical structures of DMABC-6DMA and DMABT-3DMA for OBP-tag2, an orthogonal OBP variant that exhibits anionic residues at the entrance of the binding pocket. **h.** Fluorescence spectra of DMABC-6DMA and DMABT-3DMA (1.0 μM) in the presence of OBP-tag and OBP-tag2 (5.0 μM). The excitation wavelengths for the DMAC-FA and DMABT-FA probes were 450 and 540 nm, respectively. **i.** Live-cell multicolor fluorescence imaging of U2OS cells expressing OBP-tag in mitochondria and OBP-tag2 in nuclei after staining with DMABC-FA6 and DMABT-3DMA in the presence of 0.1 μM bafilomycin A1. The excitation wavelengths of DMABC-FA6 and DMABT-3DMA were 473 and 635 nm, respectively. Scale bar: 20 μm.

To achieve orthogonal labeling, we exploited electrostatic interactions between probes and the OBP binding pocket. The crystal structure of human OBP2a reveals that positively charged residues (lysine and arginine) are enriched at the entrance of the binding pocket^33,34^, suggesting electrostatic complementarity with the anionic carboxylate groups of EverGreen probes. We designed a weakly basic probe, DMABC-6DMA, by replacing the carboxyl group with a weakly basic tertiary alkylamino group (Fig. 6g). We then introduced four charge-reversal mutations (R29E, K62E, K82E, R108E) at the binding pocket entrance, yielding OBP-tag2 (p*I* = 4.6, from 6.8). DMABC-FA6 showed markedly reduced fluorescence with OBP-tag2, consistent with electrostatic repulsion, whereas DMABC-6DMA showed a substantial fluorescence increase (Fig. 6h). A red cationic probe, DMABT-3DMA, also showed pronounced fluorescence enhancement with OBP-tag2 (Fig. 6h). Quantitative comparison confirmed orthogonal selectivity: DMABC-FA6 bound 10.5-fold more strongly to OBP-tag, whereas DMABT-3DMA bound 21-fold more strongly to OBP-tag2 (Supplementary Fig. 13a, 13b, Supplementary Tables 5 and 6).

To test orthogonal labeling in cells, we expressed OBP-tag in mitochondria and OBP-tag2 in the nucleus, and stained with DMABC-FA6 and DMABT-3DMA. Green and red fluorescence signals were observed in the expected compartments (Fig. 6i). We note that the cationic probes accumulated in acidic lysosomal compartments, requiring co-treatment with bafilomycin A1^35^ to suppress nonspecific signals (Supplementary Fig. 13c, 13d). This pharmacological intervention limits the current applicability under physiological conditions. Nevertheless, these results establish that the charge-reversal strategy provides a rational design principle for generating orthogonal tag–probe pairs within the OBP scaffold. Extending this approach to multicolor single-molecule tracking will require kinetic tuning of each orthogonal pair to satisfy the design criteria for regime iv, as well as improvement of probe selectivity to eliminate the need for pharmacological intervention in live-cell applications.

## Discussion

Reversible labeling strategies have enabled ensemble live-cell imaging, including super-resolution microscopy, that is effectively resilient to photobleaching. Yet continuous single-molecule tracking—the most stringent application of exchange-based labeling—has remained experimentally unachieved. The present results clarify why. Continuous tracking requires the simultaneous satisfaction of two quantitative kinetic conditions: dissociation prior to photobleaching and rebinding within a single imaging frame. While these requirements have been discussed qualitatively in prior studies, they have not been systematically articulated as quantitative design criteria, nor has any reported system satisfied both at the single-molecule level. EverGreen occupies this previously unachieved kinetic regime at the single-molecule level.

The critical bottleneck was not association kinetics alone. Several exchangeable systems achieve association rates approaching the diffusion limit and have been applied to single-molecule measurements. However, their dissociation rates remain comparable to or slower than the photobleaching rate, placing them in a regime where photobleaching-induced damage compromises tag function before exchange can renew the signal^16,20,22^. Our single-molecule measurements show that EverGreen probes dissociate on a timescale (∼40 ms) well before excitation-dependent photophysical processes become limiting, as evidenced by the excitation-power independence of *k*_off_ (Fig. 3d). This pre-bleach dissociation preserves the tag for sustained turnover cycles. Thus, the decisive advance is not merely fast binding, but rapid and repeated renewal without cumulative tag damage.

Reaching this fast-exchange regime has implications that extend beyond prolonged observation time. In conventional single-molecule fluorescence imaging, temporal resolution, excitation intensity, and observation duration compete within a finite photon budget; investigators typically adopt the minimum frame rate required for tracking in order to conserve fluorophore lifetime. When probe turnover outpaces both photobleaching and the imaging frame rate, these constraints are substantially relaxed. This decoupling transforms experimental design rather than merely extending observation time. Imaging can be performed at higher temporal resolution than strictly required for spatial tracking, allowing the stochastic ON/OFF dynamics of probe exchange to be resolved at each molecular locus. Continuous single-molecule imaging was also achieved in the absence of an oxygen-scavenging system, confirming that rapid probe exchange can sustain fluorescence without requiring photochemical stabilization.

The rapid exchange dynamics accessible in this regime introduce an additional analytical dimension into single-molecule imaging. Because ON and OFF dwell times are governed by well-defined binding kinetics under experimentally tunable conditions, rather than by photophysical processes, they encode information intrinsic to the labeled molecule. In principle, this kinetic fingerprint may assist in discriminating genuine single-molecule signals from static background, validating molecular identity, and estimating molecular occupancy at a given locus from discrete intensity levels. Fluorescence intermittency in this regime ceases to be an experimental nuisance and may instead become a source of mechanistic information. Systematic exploration of these capabilities will be an important direction for future work.

Extending continuous single-molecule tracking to the intracellular environment introduces two additional requirements: membrane permeability and labeling specificity within the cell. These conditions present an inherent trade-off. Membrane permeation demands hydrophobic character. Yet hydrophobic probes can partition into intracellular hydrophobic compartments, where they may generate background fluorescence. At the ensemble level, EverGreen satisfies both requirements. The probes are cell-permeable and provide wash-free labeling with signal-to-background ratios exceeding 10-fold. Single-molecule detection, however, would require substantially higher signal-to-background ratios.

Tracking multiple molecular species simultaneously is essential for resolving their interactions and coordination. Unlike genetically encoded fluorescent proteins, fluorogenic labeling systems require fluorogens to find their cognate tags from solution. Spectral diversity alone does not enable multicolor imaging; orthogonal tag–fluorogen pairs with minimal cross-reactivity are essential. The OBP family, with its diverse ligand specificities^23^, provides a natural foundation for generating orthogonal pairs. In this study, we further demonstrated that protein engineering can generate orthogonal selectivity within the same scaffold through charge reversal at the binding pocket entrance. Practical multicolor single-molecule tracking will require kinetic tuning and improvement of fluorogenic contrast for each orthogonal pair.

This work establishes a quantitative kinetic framework for continuous single-molecule tracking and demonstrates its experimental realization. The OBP tag is only 19 kDa, smaller than GFP (27 kDa) and HaloTag (33 kDa). Yet it sustains continuous single-molecule tracking for over 24 hours, no longer limited by the photon budget of individual fluorophores. This opens direct access to slow conformational dynamics, rare state transitions, and long-range temporal correlations within individual molecular trajectories. This capability would contribute to resolving the kinetic connectivities between coexisting structural states that cryo-electron microscopy has revealed.

## Methods

Detailed synthetic procedures and characterization data for all compounds, supplementary tables, and NMR spectra, are provided in the Supplementary Information.

### Materials and instruments

All chemicals were of the highest commercially available grade and were purchased from Tokyo Chemical Industry, FUJIFILM Wako Pure Chemical Corporation, Sigma-Aldrich, and Kishida Chemical Co. Reagents were used as received without additional purification. Enzymes used for molecular biological manipulations were purchased from Takara Bio and New England Biolabs. NMR spectra were recorded with a Bruker Ascend 500 NMR spectrometer with 500 MHz for ^1^H and 125 MHz for ^13^C NMR using tetramethylsilane as an internal standard. High-resolution mass spectra (HRMS) were recorded using JEOL JMS-700. Normal-phase column chromatography was conducted using silica gel BW-300 (Fuji Silysia Chemical Ltd.). Reversed-phase high-performance liquid chromatography (RP-HPLC) purification was conducted using an Inertsil ODS-3 column (4.6 or 10.0 mm × 250 mm, GL-Science, Inc.). Gel permeation chromatography was performed with a JAIGEL 1H–2H column (Japan Analytical Industry Co., Ltd.) using a system consisting of an LC-6AD pump and an SPD-20A detector (Shimadzu). Gel-filtration chromatography was performed using a Superdex 75 10/300 GL column (GE Healthcare Life Sciences) connected to an ÄKTA Explorer system (GE Healthcare Life Sciences) or an NGC Chromatography System (Bio-Rad). Spectroscopic experiments were carried out using a fluorometer (Hitachi F7000 spectrometer) with a photomultiplier voltage of 700 V and an absorption spectrophotometer (JASCO, V-650). Live-cell fluorescence images were obtained using a confocal laser scanning microscope (Andor BC43 (Oxford Instruments) or FV10i (Olympus) with a 60× lens with an oil-immersion objective.

### Preparation of recombinant proteins (OBP-tag, OBP-tag2)

*E. coli* (BL21 (DE3) (Novagen)) cells transformed with pET21b(+)-OBP-tag-His or pET21b(+)-OBP-tag2-His were cultivated in Luria Bertani medium containing 100 /of ampicillin at 37 °C. When the OD_600_ of the culture medium reached 0.6–0.8, the culture flask was incubated for 24 hours with 400 μM IPTG at 16 °C. The cells were harvested by centrifugation at 5000 rpm for 15 min, resuspended in bind buffer (50 mM sodium phosphate, 300 mM NaCl, pH 8.0), and lysed by sonication in five cycles of 2 min on and 2 min off. Following cell lysis, the supernatant was obtained by centrifugation at 5,000 rpm for 20 min and passed through a column packed with cOmplete His-Tag Purification Resin (Roche). The resin was then washed with 50 mM sodium phosphate buffer (pH 8.0) containing 300 mM NaCl and 5 mM imidazole. Proteins adsorbed onto the resin were eluted in 50 mM sodium phosphate buffer (pH 8.0) containing 300 mM NaCl and 250 mM imidazole. The eluted fraction was further purified by size exclusion chromatography (Superdex^TM^ 75 10/300 GL, GE healthcare) using running buffer (20 mM HEPES, 150 mM NaCl, pH 7.4). SDS-PAGE analysis further confirmed the purity and size of the protein. The protein concentration was measured using absorbance at 280 nm. The purified protein was divided into small volumes, flash-frozen with liquid nitrogen, and stored in a −80 °C freezer.

### Preparation of recombinant kinesin–OBP fusion proteins

Monomeric K351–OBP-tag and dimeric K560–OBP fusion proteins were generated using distinct expression systems. The K351 construct (amino acids 1–351 of mouse KIF5C) was fused to OBP, subcloned into pET21a(+) (Novagen), and expressed in *E. coli* with a C-terminal octa-histidine tag. Protein expression was induced with isopropyl *β*-D-1-thiogalactopyranoside (IPTG), and the cells were harvested 15 h after induction. The K560–OBP construct was generated using the Bac-to-Bac Baculovirus Expression System. The K560 sequence (amino acids 1–560 of mouse KIF5C) fused to OBP was subcloned into a pFastBac donor vector and transposed into a bacmid in *E. coli* DH10Bac competent cells, following the manufacturer’s protocol. Recombinant bacmid DNA was then prepared for baculovirus production.

Cell pellets were washed with ice-cold PBS and resuspended in lysis buffer (50 mM HEPES, 10 mM imidazole, 300 mM KCl, 5 mM magnesium acetate, 20 μg mL^−1^ DNase I, 0.5 mg mL^−1^ lysozyme, pH 7.4 adjusted with KOH) supplemented with 1 mM ATP and protease inhibitors. Cells were disrupted by sonication, and soluble proteins were collected by centrifugation at 40,000 rpm for 10 min at 4 °C. The supernatant was applied to a TALON-immobilized metal affinity chromatography column (Takara). After washing with high-salt buffer (40 mM HEPES, 30 mM imidazole, 300 mM KCl, 1 mM magnesium acetate, pH 7.4 adjusted with KOH) supplemented with 10 μM ATP, the bound proteins were eluted with elution buffer (30 mM HEPES, 250 mM imidazole, 30 mM KCl, 2 mM magnesium acetate, pH 7.2 adjusted with KOH) containing 10 μM ATP and 10 μg mL^−1^ leupeptin. Peak fractions were pooled, rapidly frozen in liquid nitrogen, and stored at −80 °C until use.

### Plasmid constructions

**pET21b(+)-OBP(wt)-His**.

To construct pET21b(+)-OBP(wt)-His, the fusion DNA fragment encoding wild-type human OBP2a was prepared from the synthetic gene construct pUCFa-OBPIIa (FASMAC) by PCR using the following primers:

Fwd_primer: 5-GATGACGAATTCCATGCTGAGCTTTACCCTCGAAGAAG-3 s

Fwd_primer: 5-GATGACGAATTCCATGCTGA-3

Rev_Primer: 5-AAGATCAAGCTTATGCTCCAGAACACACGAG-3 s

Rev_Primer: 5-AAGATCAAGCTTATGCTCCAGAA-3

The OBP(wt) fragment and pET21b(+) empty vector were digested with *Eco*RI and *Hind*III and ligated to form pET21b(+)-OBP(wt)-His.

**pET21b(+)-OBP-tag-His.**

To construct pET21b(+)-OBP-tag-His, a point mutation (C99S/K112N) was generated in the pET21b(+)-OBP(wt)-His plasmid using Quikchange II Site-Directed Mutagenesis kit (Agilent Technologies) and the following primers:

5-GATGACTACGTGTTCTACAGCAAAGATCAACGTCGTGGT-3 (C99S, forward)

5-ACCACGACGTTGATCTTTGCTGTAGAACACGTAGTCATC-3 (C99S, reverse)

5-CTTCGCTATATGGGCAACCTGGTTGGGCGCAAT-3 (K112N, forward)

5-ATTGCGCCCAACCAGGTTGCCCATATAGCGAAG-3 (K112N, reverse)

**pET21b(+)-OBP-tag2-His.**

For the construction of pET21b(+)-OBP-tag2-His, point mutations (R29E/K62E/K82E/R108E) were generated in pET21b(+)-OBP-tag-His using Quikchange II Site-Directed Mutagenesis kit and the following primers:

5-GACTTTCCGGAAGATGAACGTCCACGCAAAGTAAG-3 (R29E, forward)

5-CTTACTTTGCGTGGACGTTCATCTTCCGGAAAGTC-3 (R29E, reverse)

5-GAAGATCGGTGCATTCAGGAAAAGATTCTGATGCGCAAAAC-3 (K62E, forward)

5-GTTTTGCGCATCAGAATCTTTTCCTGAATGCACCGATCTTC-3 (K62E, reverse)

5-GCGTATGGCGGTCGTGAACTGATCTATCTCCAG-3 (K82E, forward)

5-CTGGAGATAGATCAGTTCACGACCGCCATACGC-3 (K82E, reverse)

5-CGTCGTGGTGGACTTGAATATATGGGCAACCTG-3 (R108E, forward)

5-CAGGTTGCCCATATATTCAAGTCCACCACGACG-3 (R108E, reverse)

**pCMV6-OBP-tag.**

A wild-type human OBP2a-encoding plasmid was purchased from ORIGENE. For the construction of pCMV6-OBP-tag, a point mutation (C99S/K112N) was generated in the plasmid using Quikchange II Site-Directed Mutagenesis kit and the following primer pairs:

5-CGACTACGTCTTTTACAGCAAAGACCAGCGCCGTG-3 (C99S, forward)

5-CACGGCGCTGGTCTTTGCTGTAAAAGACGTAGTCG-3 (C99S, reverse)

5-CCTGCGCTACATGGGAAACCTTGTGGGTAGGAATC-3 (K112N, forward)

5-GATTCCTACCCACAAGGTTTCCCATGTAGCGCAGG-3 (K112N, reverse)

**pCMV6-OBP-tag-NLS.**

To construct pCMV6-OBP-tag-NLS, the fusion DNA fragment of NLS was prepared from pcDNA3.1(+)-BL-NLS^36^ by PCR using the following primers:

Fwd_primer: 5-GATGACACGCGTGGAGGATCTGGAGGATCTGA-3

sFwd_primer: 5-GATGACACGCGTGGAGG-3

Rev_Primer: 5-AAGATCGCGGCCGCGTTACCTTTCTCTTTTTCTTAGGATCAAC-3 s

Rev_Primer: 5-AAGATCGCGGCCGCGT-3

The NLS fragment and pCMV6-OBP-tag were digested with *Mlu*I and *Not*I and were ligated to form pCMV6-OBP-tag-NLS.

**pKmcl-2xCox8-OBP-tag.**

To construct pKmcl-2xCox8-OBP-tag, a fusion DNA fragment of OBP-tag was prepared from pCMV6-OBP-tag by PCR using the following primers:

Fwd_primer: 5-GATGACGGATCCATGCTGTCCTTCACC-3 s

Fwd_primer: 5-GATGACGGATCCATGCTG-3

Rev_Primer: 5-AAGATCGAATTCTTAGTGTTCGAGAACGCAGCTT-3 s

Rev_Primer: 5-AAGATCGAATTCTTAGTGTTCGAG-3

The OBP-tag fragment and pKmcl-2xCox8-BL^37^ were digested with *Bam*HI and *Eco*RI and ligated to form pKmcl-2xCox8-OBP-tag.

**pCMV6-OBP-tag2-NLS.**

For the construction of the pCMV6-OBP-tag2-NLS, point mutations (R29E/K62E/K82E/R108E) were generated in pCMV6-OBP-tag-NLS using Quikchange II Site-Directed Mutagenesis kit and the following primer pairs:

5-GACTTTCCGGAGGACGAGAGGCCCAGGAAGGTG-3 (R29E, forward)

5-CACCTTCCTGGGCCTCTCGTCCTCCGGAAAGTC-3 (R29E, reverse)

5-GGATCGGTGCATCCAGGAGAAAATCCTGATGCGG-3 (K62E, forward)

5-CCGCATCAGGATTTTCTCCTGGATGCACCGATCC-3 (K62E, reverse)

5-GCCTATGGGGGCAGGGAGCTCATATACCTGCAG-3 (K82E, forward)

5-CTGCAGGTATATGAGCTCCCTGCCCCCATAGGC-3 (K82E, reverse)

5-CGCCGTGGGGGCCTGGAGTACATGGGAAACCTTG-3 (R108E, forward)

5-CAAGGTTTCCCATGTACTCCAGGCCCCCACGGCG-3 (R108E, reverse)

**pET21a(+)-K351–OBP-His.**

To construct pET21a(+)-K351–OBP-His, a fusion DNA fragment encoding amino acids 1–351 of mouse KIF5C fused to OBP was prepared by PCR from a synthetic gene construct using the following primers:

Ins_Fwd_primer: 5-GGCGGAGGAGGATCTATGCTGTCCTTCACCCTGG-3

Vec_Fwd_primer: 5-GGCGGAGGCGGAAGTCATCACCATCACCATCACCATC-3

Ins_Rev_Primer: 5-GAACACGGCGGAGGCGGAAGT-3

Vec_Rev_Primer:

5-GAGTGGAAAAAGAAATATGAAAAAGAGAAAGAGGGCGGAGGAGGATCT-3

The K351–OBP fragment and the linearized pET21a(+) empty vector were assembled using the In-Fusion HD Cloning Kit (Takara Bio) according to the manufacturer’s protocol to form pET21a(+)-K351–OBP-His.

**pFastBac-K560–OBP-His.**

To construct pFastBac-K560–OBP-His, a fusion DNA fragment encoding amino acids 1–560 of mouse KIF5C fused to OBP was prepared by PCR from a synthetic gene construct using the following primers:

Ins_Fwd_primer: 5-GGAGATAGGCGGACCATGCTGTCCTTCACCCTGGAGG-3 Vec_Fwd_primer: 5-CATCACCATCACCATCACCATC-3

Ins_Rev_Primer: 5-GGAAGCTGCGTTCTCGAACACCATCACCATCACCAT-3

Vec_Rev_Primer: 5-GGACCTGGGGGAGATAGGCGGACC-3

The K560–OBP fragment and linearized pFastBac donor vector were assembled using the In-Fusion HD Cloning Kit (Takara Bio) according to the manufacturer’s protocol to form pFastBac-K560–OBP-His. The donor plasmid was transformed into *E. coli* DH10Bac competent cells to generate recombinant bacmid DNA, according to the Bac-to-Bac Baculovirus Expression System protocol.

### Fluorescence and absorption spectroscopy

For fluorescence measurements, OBP-tag or OBP-tag2 was incubated with probes at 37 °C in 20 mM HEPES buffer, pH 7.4, containing 150 mM NaCl (0.2% DMSO used). The spectra were recorded immediately after the addition of the probes at an excitation wavelength of 413 nm for DMAC-type probes, with slit widths of 5.0 nm and 2.5 nm for both excitation and emission. The fluorescence spectra were recorded at excitation wavelengths of 467, 467, 467, 465, and 450 nm for DMABC-FA6, DMABC-FA8, DMABC-FA10, DMABC-FA12, and DMABC-6DMA, respectively, with slit widths of 5.0 nm for both excitation and emission. Fluorescence spectra were recorded at an excitation wavelength of 540 nm for DMABT-type probes with slit widths of 10.0 nm for both excitation and emission.

### Quantum yield determination

The fluorescence spectra of DMAC-type, DMABC-type, and DMABT-type probes were measured in PBS buffer (pH 7.4) containing 1% DMSO. The fluorescence quantum yields of the probes were determined using coumarin 153 in cyclohexane (Φ = 0.9), fluorescein in 0.1 N NaOH (Φ = 0.88), and rhodamine 6G in ethanol (Φ = 0.95) as references^38^. The relative quantum yields were calculated using the following equation:

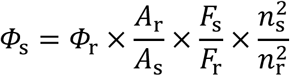

where Φ_s_ = quantum yield of the sample, Φ_r_ = quantum yield of reference, *A*_r_ = absorbance of the reference, *A*_s_ = absorbance of the sample, *F*_s_ = FL area of the sample, *F*_r_ = FL area of the reference, *n*_s_ = refractive index of the solvent used to measure sample, and *n*_r_ = refractive index of the solvent used to measure reference.

### Calculation of dissociation constant (*R*_d_)

The dissociation constant was calculated using fluorescence titration. We used a fixed concentration of probes with variable concentrations of OBP-tag, and the graphs were fitted using the following equation:

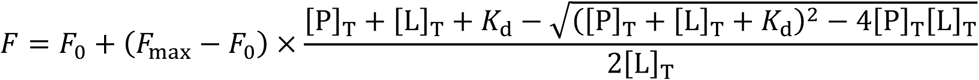

### Photostability test

The sample solutions of probes (1.0 μM, 200 μL) (0.2% DMSO) in the presence of OBP-tag (*K*_d_ value) in HEPES buffer (pH 7.4) containing 150 mM NaCl were added to a quartz cell and irradiated with an excitation light beam (4 mW/cm^2^) using a xenon light source (MAX-303, Asahi Spectra, Torrance, CA, USA) equipped with a bandpass filter (405 ± 10 nm: DMAC-type, 490 ± 10 nm: DMABC-type, and 550 ± 10 nm: DMABT-type). The fluorescence intensity was measured every 5 min for 30 min.

### Fluorescence lifetime measurement

Fluorescence lifetimes were measured by time-correlated single-photon counting (TCSPC) using a Horiba FluoroCube Fluorescence Lifetime System. The fluorescence intensity at 620 nm (free DMABC probes) and 544 nm (DMABC probes + OBP-tag) was measured using a 454 nm NanoLED pulse laser source. The fluorescence decay data were analyzed using IBH DAS6 software and fitted with the following bi-exponential function employing nonlinear least-squares deconvolution analysis:

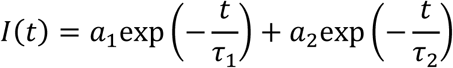

### Cell culture and transfection

Mammalian culture cells (HEK293T, COS-7, and U2OS) were seeded onto a cleaned glass coverslip and maintained in complete media (Dulbecco’s modified Eagle’s medium supplemented with 10% (v/v) FBS) under growth conditions (37 °C, 5% CO_2_). To express OBP-tags in the cytosol, nuclei, or mitochondria, the cells were transfected with pCMV6-OBP-tag, pCMV6-OBP-tag-NLS, or pKmcl-2xCox8-OBP-tag using Lipofectamine 3000 (Thermo Fisher Scientific) according to the manufacturer’s protocol. To express OBP-tag2 in the nuclei, the cells were transfected with pCMV6-OBP-tag2-NLS using Lipofectamine 3000. Hoechst33342 and MitoTracker^TM^ DeepRed were used for co-staining the nuclei and mitochondria, respectively.

### Live cell fluorescence imaging

After incubating the transfected cells at 37 °C for 24 h, the cells were washed twice with Hank’s Balanced Salt Solution (HBSS). The transfected cells were incubated with 1 μM EverGreen (DMABC-based), 2 μM DMAC, or 1 μM DMABT probes in DMEM containing 0.1–0.2% DMSO. Microscopic images of the cells were acquired using a confocal laser scanning microscope immediately after labeling. For DMABT-3DMA, cells were incubated with the probe in the presence of 0.1 μM Bafilomycin A_1_ for 5 h prior to fluorescence imaging.

### Time-lapse live-cell NSPARC super-resolution imaging

After transfecting COS-7 cells with pKmcl-2xCox8-OBP-tag for 24 h, the cells were washed twice with Hank’s Balanced Salt Solution (HBSS). The cells were then incubated with 1 μM DMABC-FA6 in FluoroBrite^TM^ DMEM containing 0.1% DMSO. The cells were then imaged using a Nikon AX R with NSPARC Inverted-type Confocal-based Super Resolution Microscope without washing. Time-lapse images were acquired with an excitation wavelength of 488 nm and a ×60 lens (Numerical Aperture: 1.42 and Refractive Index: 1.515) for 15 min at 5-sec intervals.

### Live cell fluorescence imaging with probe exchange

After transfecting COS-7 cells with pCMV6-OBP-tag-NLS for 24 h, the cells were washed twice with HBSS. The cells were then incubated with 2 μM DMAC-FA10 in FluoroBrite^TM^ DMEM containing 0.2% DMSO. After imaging, the cells were washed three times with HBSS and stained with 1 μM DMABC-FA6 in FluoroBrite^TM^ DMEM containing 0.1% DMSO. After imaging, the cells were washed three times with HBSS and stained with 1 μM DMABT-FA2 in FluoroBrite^TM^ DMEM containing 0.1% DMSO. The cells were imaged using a confocal laser scanning microscope (BC43, Andor) with excitation wavelengths of 405, 488, and 635 nm.

### Multicolor fluorescence imaging using green and red probes

U2OS cells were co-transfected with pCMV6-2xCox8-OBP-tag and pCMV6-OBP-tag2-NLS using Lipofectamine 3000 (Invitrogen) according to the manufacturer’s protocol. After incubating the transfected cells in 5% CO_2_ at 37 °C for 24 h, the cells were washed twice with HBSS. The cells were then incubated with 1 μM DMABC-FA6, 1 μM DMABT-3DMA, and 0.1 μM Bafilomycin A_1_ in DMEM for 5 h. Microscopic images of the probe-treated cells were recorded using a confocal laser-scanning microscope (FV10i) with excitation at 473 and 635 nm.

### Single-molecule imaging

For single-molecule imaging and kinetic analysis of the OBP-based labeling system, glass coverslips were passivated with polyethylene glycol (PEG) to suppress the nonspecific adsorption of K351–OBP to the surface.

Glass coverslips were cleaned by incubation in 1 M potassium hydroxide for 15 min, followed by plasma treatment for 15 min (FEMTO; Diener Electronic). The coverslips were then heated at 150 °C for 10 min and allowed to cool to room temperature. Surface amination was performed by incubating the coverslips in freshly prepared 3-(2-aminoethylamino)propyltriethoxysilane (LS-4265, Shin-Etsu Chemical) for 20 min at room temperature. After extensive rinsing with ultrapure water and drying, PEGylation was performed by sandwiching 10 μL of PEG solution between a pair of coverslips and incubating for 1.5 h. The PEG solution consisted of methoxy-PEG–NHS and biotin-PEG–NHS (NOF Corporation) mixed at a molar ratio of 50:1 in freshly prepared 1 M MOPS buffer (pH 7.5). After PEGylation, the coverslips were rinsed with water, dried, and stored in a vacuum desiccator until use. Flow chambers were assembled using PEG-coated coverslips and double-sided adhesive tape (No. 5603, Nitto Denko). Neutravidin (0.25 mg mL^−1^, Invitrogen) was introduced into the chamber and incubated for 1 min, followed by washing with assay buffer (AB: 80 mM PIPES, 1 mM EGTA, 0.2 mM AMP-PNP, 2 mM MgSO_4_, 2 mM dithiothreitol, 0.2 mg mL^−1^ glucose oxidase, 40 μg mL^−1^ catalase, 1 mM glucose, 0.2 mg mL^−1^ casein, 10 μM taxol, pH 6.9 adjusted with KOH). Tetramethylrhodamine-labeled microtubules (3% labeling, 5% biotinylation) were introduced into the chamber and incubated for 1 min to allow surface attachment, followed by washing with AB. The surface was then blocked by incubation with blocking buffer (2 mg mL^−1^ casein) for 2 min and washed again with AB. The DMABC-FA8 probe diluted in motility buffer was subsequently introduced, and the chamber was sealed with a silicone sealant (Dent Silicone-V, SHOFU). Fluorescence imaging was performed using a total internal reflection fluorescence microscope (IX83, Olympus) equipped with a UPlanApo 100×/NA 1.50 oil-immersion objective and an ORCA-Fusion CMOS camera (Hamamatsu Photonics). Image acquisition was controlled using MetaMorph software.

### Long-term single-molecule imaging

For long-term single-molecule imaging, glass coverslips were passivated with PEG–biotin using the same procedure described above. K351–OBP was incubated with biotinylated anti-His-tag monoclonal antibody (MBL) at an equimolar ratio for 30 min at room temperature prior to surface immobilization. NeutrAvidin (0.25 mg mL^−1^; Invitrogen) was introduced into the flow chamber and incubated for 1 min, followed by washing. The antibody–K351–OBP complex (100 nM) was then flowed into the chamber and incubated for 5 min to allow immobilization via biotin–avidin interactions. Unbound molecules were removed by washing with blocking buffer (80 mM PIPES, 1 mM EGTA, 2 mM MgSO_4_, 2 mg mL^−1^ casein, pH 6.9 adjusted with KOH), which also served to block residual nonspecific binding sites. Subsequently, the DMABC-FA8 probe diluted in assay buffer (80 mM PIPES, 1 mM EGTA, 2 mM MgSO_4_, 0.2 mg mL^−1^ glucose oxidase, 40 μg mL^−1^ catalase, 1 mM glucose, 0.2 mg mL^−1^ casein, 0.1% Pluronic F-127, pH 6.9 adjusted with KOH) was introduced into the chamber. The chamber was sealed with a silicone sealant (Dent Silicone-V, SHOFU) and immediately imaged using TIRF microscopy.

### Image analysis of reversible fluorescence probe

Time-lapse TIRF image sequences were analyzed using Fiji/ImageJ and custom-written Python scripts. Fluorescence intensity time traces were extracted for individual diffraction-limited spots defined by fixed regions of interest (ROIs). For each ROI, the mean intensity per frame was exported from ImageJ as CSV files and used for downstream analysis. To quantify the probe exchange dynamics, intensity trajectories were converted into binary ON/OFF state sequences by thresholding. Frames with intensities within a predefined signal range were assigned to the ON state, whereas those outside this range were assigned to the OFF state. To minimize spurious state transitions arising from single-frame intensity drops, brief gaps of one frame were bridged and treated as part of the same ON event. The ON– and OFF-dwell times were defined as the durations of consecutive ON and OFF segments, respectively, and were converted from frame counts to time using the acquisition frame interval. Abnormally short or long events were excluded to reduce the contribution of noise and rare outliers. Dwell times obtained from multiple ROIs and independent recordings acquired under the same conditions were pooled for statistical analysis. The empirical cumulative distribution functions (CDFs) of the ON– and OFF-dwell times were computed and fitted with a single-exponential model to estimate the rate constants. The dissociation rate constant (*k*_off_) was obtained from the ON-dwell time distributions. The observed reappearance rate (*k*_obs_) was obtained from the OFF-dwell time distributions measured at each probe concentration. The association rate constant (*k*_on_) was determined from the concentration dependence of *k*_obs_ (linear fit of *k*_obs_ versus probe concentration). All fitting procedures were performed using non-linear least squares in Python.

## Acknowledgments

This work was supported by a Grant-in-Aid for Transformative Research Area (A) “Latent Chemical Space” [23H04880 and 23H04881 (K.K.)] from the Ministry of Education, Culture, Sports, Science and Technology, Japan. This research was also supported by Japan Society for the Promotion of Science (JSPS) KAKENHI grants (JP21H04706, JP23KK0106, JP24H00494 to K.K.; JP20K05747, JP21H00401, JP22H05425, JP25K01915 to M.M.; JP19H03394, JP19H05794, JP19H05795, JP22H04926 and JP21H05254 to Y.O.; JP22H02798 and JP25K02438 to Y.O. and T.K.); Japan Science and Technology Agency (JST) PRESTO (JPMJPR22EC to M.M.); JST CREST grants (JPMJCR1852, JPMJCR20E2 and JPMJCR24T2 to Y.O.); JST Moonshot Research and Development Program grant (JPMJMS2025-14 to Y.O.); Japan Agency for Medical Research and Development (AMED) CREST grant (JP23gm1710001s0502 to Y.O. and T.K.); the JSPS CORE-to-CORE Program “Asian Chemical Biology Initiative”. This work was also supported by the IFReC advanced postdoc program (S.I.R. and K.K.). The parts of the schematic illustrations in the figures were created with BioRender.com. The authors thank Namiko Yamada for technical assistance in the experiments using recombinant OBP-tag. We also thank lab members at RIKEN and the University of Tokyo for their discussion and support, especially Ms. Xu Shang Dan, Yukiko Onishi, Mikako Hayashi, Sayuri Yamamoto, and Junko Asada for technical support, and Ms. Manaho Kakiuchi, Ryoko Araki, and Tomoko Furuya for secretarial assistance. We thank Yugo Inutsuka for assistance with single-molecule data analysis. The authors thank the analytical instrument facility, Graduate School of Engineering, the University of Osaka, for HRMS analysis; Nikon Imaging Center (NIC), the University of Osaka, for AX R with NSPARC Inverted-type Confocal-based Super Resolution Microscope.

## Supplementary Information

### Supplementary Note 1 | Quantitative requirements and probabilistic design framework for continuous single-molecule tracking

This Note provides quantitative derivations of the two kinetic requirements for continuous single-molecule tracking presented in the main text, establishes their probabilistic interpretation, and estimates the threshold values used to define the regime boundaries in Fig. 1b.

#### Requirement 1: pre-bleach dissociation (*k*_*off*_)

If a bound probe photobleaches before dissociating, two consequences follow: the tag remains dark while waiting for the bleached probe to leave, and the reactive oxygen species generated during photobleaching can irreversibly damage the tag itself (Fig. 1a, ii)^22^. To avoid both, each probe molecule should dissociate before photobleaching occurs (Eq. 1 in the main text). When a probe is bound, dissociation and photobleaching proceed as competing first-order processes with rate constants *k*_off_and *k*_bleach_, respectively. The probability that dissociation occurs before photobleaching is

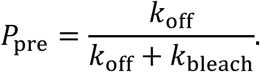

When *k*_off_ exceeds *k*_bleach_ by approximately a factor of 20, *P*_pre_ > 0.95 and photobleaching during the bound state becomes statistically negligible. Under representative TIRF single-molecule imaging conditions, organic fluorophores typically exhibit photobleaching time constants of a few seconds to several tens of seconds (e.g., τ ≈ 3 s for Alexa Fluor 647 under TIRF illumination at 60 W cm^−2^ ^10^), corresponding to effective bleaching rates on the order of 0.1–1 s ^−1^. Accordingly, we adopt *k*_off_ ≳ 20 × *k*_bleach_ as a practical design criterion for pre-bleach dissociation with > 95% probability. Given that *k*_bleach_ is typically on the order of 1 s^−1^ or below, this corresponds to *k*_off_ ≳ 20 s^−1^, which defines the horizontal boundary in Fig. 1b. For EverGreen, *k*_off_ ≈ 24 s^−1^ substantially exceeds this criterion, yielding *P*_pre_ > 0.95. This is directly confirmed by the observation that the measured *k*_off_ is independent of excitation laser power (Fig. 3d).

#### Requirement 2: rebinding capacity (*k*_*o*n_ × *CR*)

If a new probe fails to bind before the next imaging frame, the tag is recorded as dark, introducing a gap in the fluorescence trajectory (Fig. 1a, i and iii). To avoid such gaps, probe rebinding must occur within a single camera frame. The rebinding rate is the product of the association rate constant *k*_on_ and the free probe concentration, so a straightforward way to accelerate rebinding is to raise the probe concentration. However, in single-molecule detection, the probe concentration is bounded by the requirement that background fluorescence from unbound probes remain well below the signal from a single bound molecule. This constraint depends on the detection volume, which we now estimate.

Single-molecule detection requires that individual molecules be spatially resolved within a diffraction-limited spot. By Poisson statistics, the probability of two or more molecules occupying the same detection volume is *P*(≥ 2) = 1 − *e*^−μ^(1 + μ). For μ = 0.3, this probability is ≈ 3.7%, below the ∼ 5% empirical tolerance commonly adopted in single-molecule experiments. Accordingly, μ ≲ 0.3 is used as a practical design criterion. For a typical TIRF geometry with a lateral PSF radius of ∼ 220 nm (0.61λ/NA; λ = 500 nm, NA = 1.4) and an evanescent field penetration depth of ∼ 100 nm, *V*_PSF_ ≈ π × (220 nm)^2^ × 100 nm ≈ 0.015 fL. The single-molecule condition μ = *C* × N_A_ × *V*_PSF_ ≲ 0.3 then sets the characteristic concentration scale

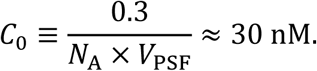

This value represents a geometric upper bound derived from the diffraction-limited detection volume. In practice, single-molecule imaging is typically performed at somewhat lower concentrations (1–10 nM) to maintain additional safety margins against background, autofluorescence, and detector noise.

For a fluorogenic system with contrast ratio CR, each free probe contributes 1/CR times the signal of a single bound molecule. At a probe concentration [probe] = CR × *C*_0_, the background-equivalent occupancy per PSF equals μ ≈ 0.3, so that the background from free probes does not exceed the expected single-molecule signal. This sets [probe]_max_ ∼ CR × *C*_0_ as the practical upper bound on probe concentration.

Requiring that the rebinding rate at this maximum concentration exceed the imaging frame rate yields the condition presented as Eq. 2 in the main text:

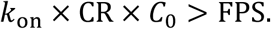

For a frame rate of 10 Hz and *C*_0_ ≈ 30 nM, the central boundary becomes *k*_on_ × CR > 300 μM^−1^s^−1^.

This boundary represents a probabilistic benchmark rather than a sharp threshold. Following probe dissociation, the waiting time for the next binding event is exponentially distributed. The probability that a new probe binds within one camera frame of duration Δ*t* = 1/FPS is

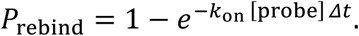

At the central boundary, the exponent equals unity, yielding *P*_rebind_ ≈ 0.63. Achieving within-frame rebinding with high probability (e.g., *P*_rebind_ > 0.95) requires *k*_on_ [probe] Δ*t* = −ln(0.05) ≈ 3, corresponding to approximately a threefold kinetic margin. For FPS = 10 Hz and *C*_0_ ≈ 30 nM, this translates to *k*_on_ × CR ≳ 1,000 μM^−1^s^−1^. For EverGreen, *k*_on_ × CR ≈ 1,050 μM^−1^s^−1^, which exceeds this benchmark.

#### Evaluation of reported systems

Supplementary Table 1 compiles the kinetic and fluorogenic parameters of EverGreen and reported exchangeable labeling systems. For EverGreen (DMABC-FA8):

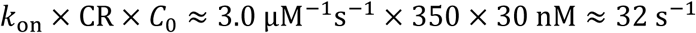

which exceeds the 10 Hz threshold, confirming that Requirement 2 is satisfied. The mean residence time of ∼40 ms satisfies Requirement 1. The evaluation of other reported systems and the interpretation of the regime map are presented in the main text.

### Supplementary Note 2 | Validation of single-molecule regime by fluorescence intensity analysis

To assess whether individual diffraction-limited fluorescent spots predominantly correspond to single K351–OBP molecules under our imaging conditions, we analyzed fluorescence intensity distributions over a range of K351–OBP concentrations. Supplementary Fig. 3 shows histograms of fluorescence intensities measured for individual spots at different protein concentrations.

At low K351–OBP concentrations (10 pM), the intensity distribution was dominated by a single population, consistent with the majority of detected signals arising from single molecules. Upon increasing the K351–OBP concentration, the overall shape of the intensity distribution broadened, and a minor higher-intensity component became detectable, reflecting an increased contribution from spots with higher fluorescence intensity.

The intensity histograms were fitted using a two-component model (see Methods) to quantitatively assess the relative contributions of the lower– and higher-intensity populations. At 10 pM K351–OBP, approximately 90% of detected signals were assigned to the lower-intensity component, which we interpret as corresponding to single-molecule events. Although the relative contribution of the higher-intensity component increased at elevated K351–OBP concentrations, the lower-intensity population remained predominant across the concentration range examined.

Based on this analysis, a K351–OBP concentration of 10 pM was used for subsequent single-molecule imaging and kinetic analyses, ensuring that the majority of detected fluorescence signals originated from individual molecules.

### Supplementary Note 3 | Single-molecule kinetic analysis of EverGreen probe exchange

To quantify the binding and dissociation kinetics underlying fluorescence signal renewal in the EverGreen system, we analyzed the dwell-time distributions of fluorescence ON and OFF states obtained from single-molecule fluorescence trajectories of OBP-tagged proteins.

#### Dissociation kinetics (***k***_*off*_)

Fluorescence ON dwell times, defined as the duration over which a fluorescent signal was continuously detected at a fixed molecular position, were extracted and analyzed. The cumulative distributions of ON dwell times were well fitted by a single-exponential function under all experimental conditions (Supplementary Fig. 5a), indicating that probe dissociation from the OBP tag follows a single-rate kinetic process.

To assess potential contributions from photophysical effects, ON dwell-time analyses were performed at multiple excitation laser powers. The observed dissociation rate constants showed no dependence on laser power (Supplementary Fig. 5b), indicating that the measured dissociation kinetics were not limited by photobleaching or laser-induced blinking. The dissociation rate constant *k*_off_was therefore determined as the average value across all laser powers, yielding *k*_off_= 24 s^−1^, corresponding to a mean binding lifetime of approximately 41 ms.

#### Association kinetics (***k***_*o***n**_)

Fluorescence OFF dwell times, defined as the intervals between successive fluorescence ON events, were analyzed at multiple probe concentrations. The cumulative distributions of OFF dwell times were also well described by single-exponential functions for all concentrations tested (Supplementary Fig. 5c), consistent with a single dominant association process. The observed reappearance rate *k*_on_increased linearly with probe concentration (Supplementary Fig. 5d). From the slope of this relationship, the association rate constant was determined to be *k*_on_ = 3.0 μM^−1^ s^−1^.

#### Equilibrium affinity

From the independently determined rate constants, the equilibrium dissociation constant was calculated as:

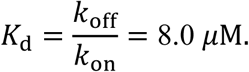

This value is in good agreement with ensemble solution measurements performed independently (*K*_d_ = 9.4 ± 0.6 μM), supporting the validity of the single-molecule kinetic analysis.

These results demonstrate that fluorescence signal renewal in the EverGreen system is governed by simple first-order dissociation and concentration-dependent association kinetics. The ability to describe both ON and OFF dwell times with single-exponential distributions indicates that probe exchange proceeds through a well-defined kinetic process, enabling quantitative single-molecule analysis without distortion by photobleaching.

**Supplementary Fig. 1.**
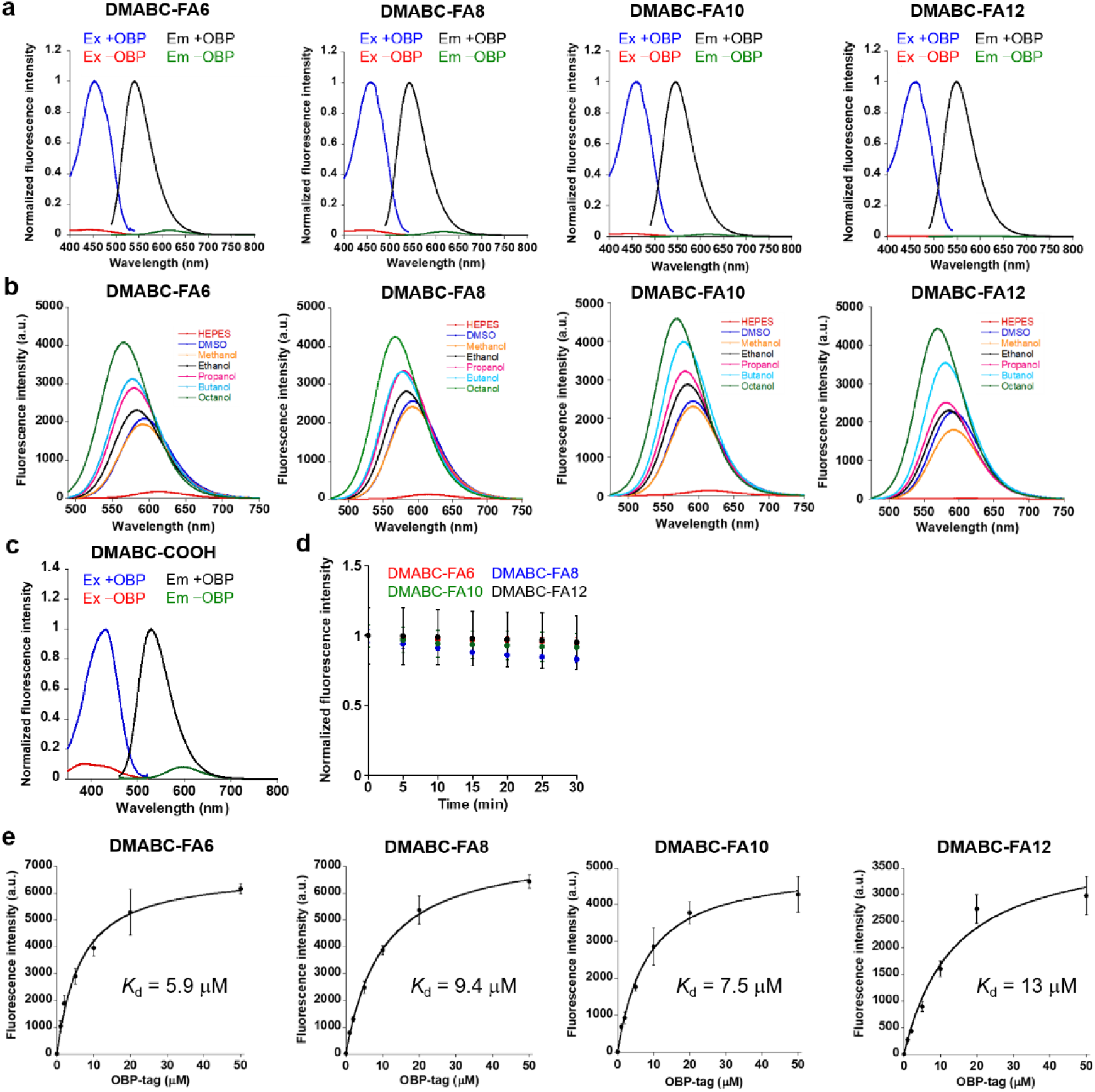
| Fluorescence and OBP-binding properties of DMABC probes. **a.** Fluorescence excitation and emission spectra of 1 μM DMABC-FA6, –FA8, –FA10, and –FA12 (0.1% DMSO) in the absence and presence of 50 μM OBP-tag in 20 mM HEPES buffer (pH = 7.4) containing 150 mM NaCl at 37 °C. λ_*ex*_= 467 nm (FA6, FA8, and FA10) and 465 nm (FA12) for emission spectra, λ_*em*_ = 550 nm for excitation spectra. **b.** Fluorescence spectra of 1 μM DMABC-probes (0.1% DMSO) in different organic solvent and 20 mM HEPES buffer (pH = 7.4) containing 150 mM NaCl at 37 °C. λ_*ex*_= 467 nm (FA6, FA8, and FA10) and 465 nm (FA12). **c.** Fluorescence excitation and emission spectra of 1 μM DMABC-COOH (0.1% DMSO) in the absence and presence of 50 μM OBP-tag in 20 mM HEPES buffer (pH = 7.4) containing 150 mM NaCl at 37 °C. λ_*ex*_ = 435 nm for emission spectra, λ_*em*_ = 550 nm for excitation spectra. **d.** Photostability of DMABC probes under continuous light illumination (490 nm, 4 mW/cm^2^). The sample solution contains 1 μM probe (0.1% DMSO) in the presence OBP-tag (50% labeling condition) in 20 mM HEPES buffer (pH = 7.4) and 150 mM NaCl at 37 °C. λ_*ex*_= 467 nm (FA6, FA8, and FA10) and 465 nm (FA12). Error bars denote standard deviation (*N* = 3). **e.** Fluorescence titration of DMABC-FA6, –FA8, –FA10, and –FA12 for different concentrations of OBP-tag at 37 °C. λ_*ex*_= 467 nm (FA6, FA8, and FA10) and 465 nm (FA12). Error bars denote standard deviation (*N* = 3).

**Supplementary Fig. 2.**
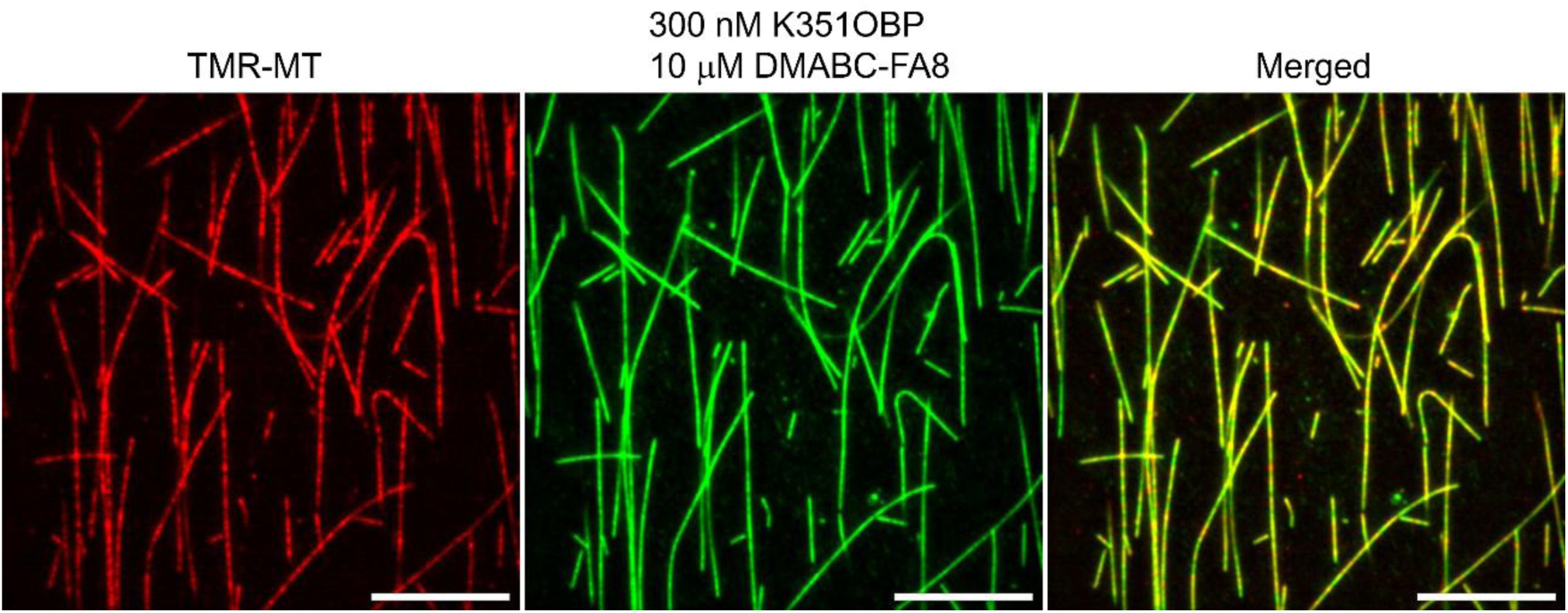
| Colocalization of EverGreen-labeled kinesin with TMR-labeled microtubules. Microtubules sparsely labeled with tetramethyl rhodamine (TMR) at 30% labeling density are shown in red (left). EverGreen fluorescence signals observed in the presence of 300 nM K351–OBP and 10 μM probe are shown in green (middle). The merged image demonstrates strong colocalization of EverGreen-labeled kinesin with TMR-labeled microtubules (right). Scale bar, 10 μm.

**Supplementary Fig. 3.**
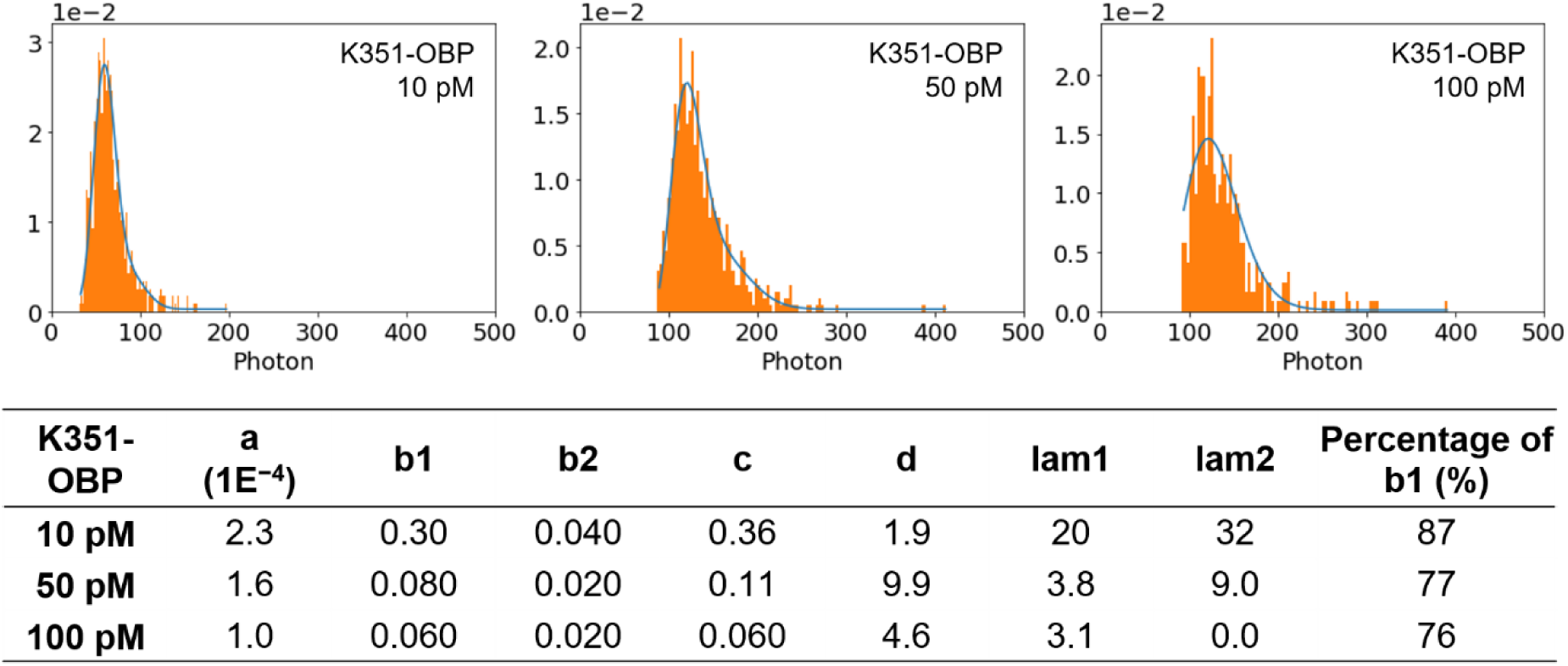
| Fluorescence intensity distributions of K351–OBP spots under different labeling densities. Histograms of fluorescence intensity measured for individual diffraction-limited spots of K351–OBP observed on microtubules at different protein concentrations. At low K351–OBP concentrations, the intensity distribution was well described by a single dominant population corresponding to single-molecule signals. At higher K351–OBP concentrations, an additional higher-intensity population emerged, consistent with multiple K351–OBP molecules occupying the same diffraction-limited area. The distributions were fitted with a sum of exponential components (see Supplementary Methods), and the relative amplitudes of the fitted components reflect the fraction of single-molecule versus multi-molecule contributions.

**Supplementary Fig. 4.**
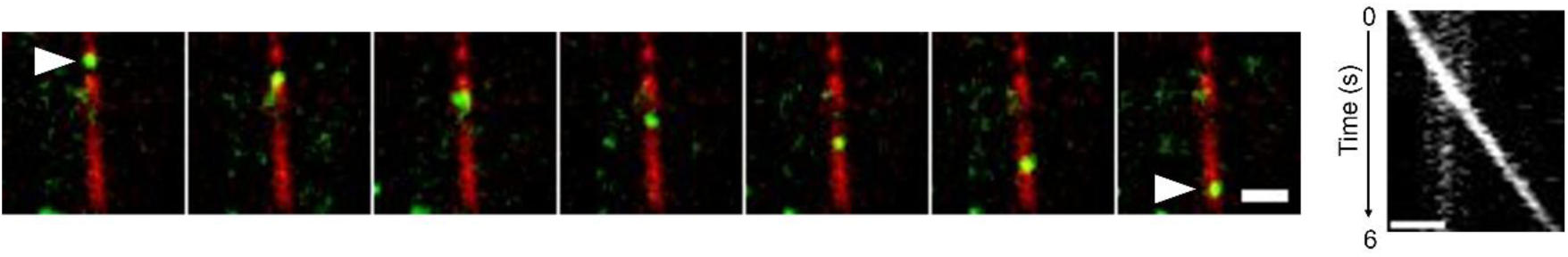
| Single-molecule motility of dimeric kinesin labeled with EverGreen. (Left) Total internal reflection fluorescence (TIRF) microscopy image showing unidirectional movement of individual dimeric kinesin molecules (K560–OBP, green) along surface-immobilized microtubules (red) in the presence of ATP. White arrowheads indicate the starting and ending positions of a representative motile kinesin molecule. This experiment was performed using DMABC-FA6 (10 μM), chosen for its low background fluorescence under the imaging conditions used. **(Right)** Kymograph generated from the trajectory shown in the left panel illustrating continuous single-molecule movement of K560 along a microtubule over time. Scale bar, 1 μm.

**Supplementary Fig. 5.**
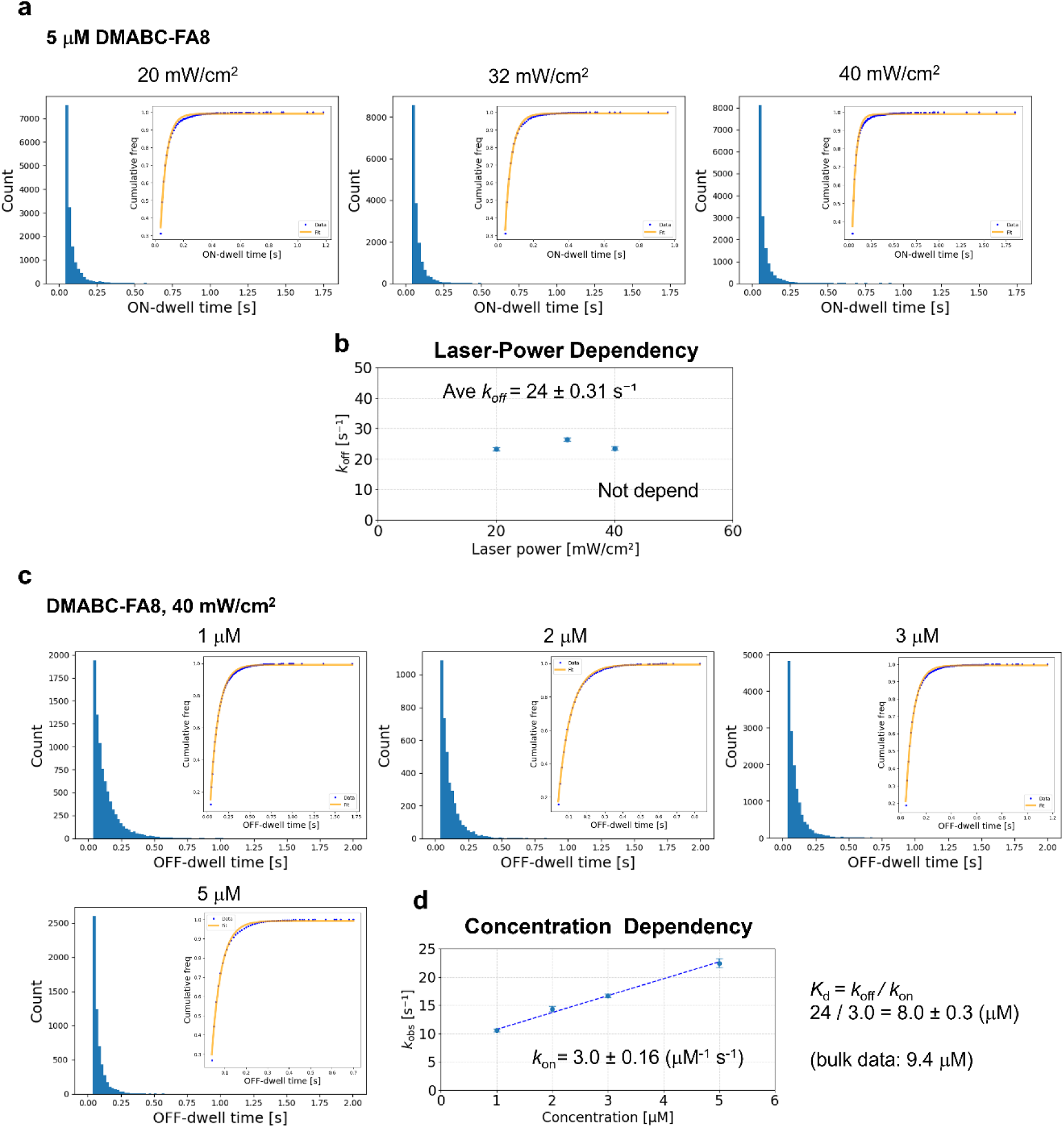
| Dissociation kinetics of the EverGreen probe DMABC-FA8. **(a)** Cumulative distributions of fluorescence ON dwell times measured under different excitation laser powers. ON dwell times were defined as the durations of continuous fluorescence signals at fixed molecular positions. For all excitation conditions, the cumulative distributions were well described by single-exponential functions, indicating a single dominant dissociation process. **(b)** Observed dissociation rate constants (*k*_*obs*_) extracted from the fits in (a), plotted as a function of excitation laser power. *k*_*obs*_showed no systematic dependence on excitation intensity, indicating that probe dissociation is not limited by photophysical effects. The dissociation rate constant was therefore determined as the average value, yielding *k*_*off*_ = 24 s^−1^. **(c)** Cumulative distributions of fluorescence OFF dwell times measured at different probe concentrations, fitted with single-exponential functions. **(d)** Concentration dependence of the observed reappearance rate constant *k*_*obs*_. Linear fitting yields an association rate constant of *k*_*on*_ = 3.0 μ*M*^−1^ *s*^−1^. Error bars in b and d represent 90% confidence intervals obtained by bootstrap resampling (N_boot = 1000).

**Supplementary Fig. 6.**
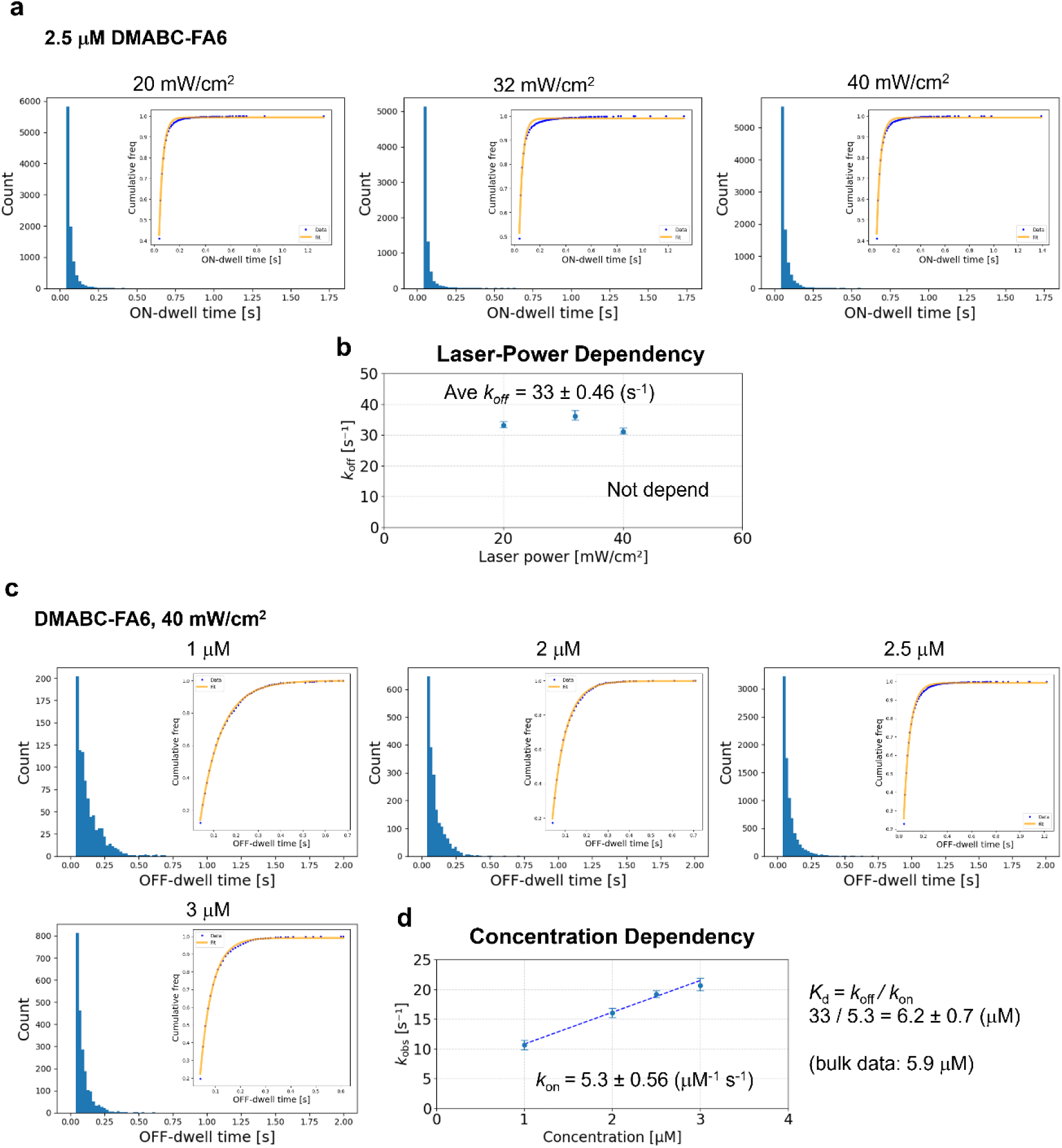
| Dissociation kinetics of the EverGreen probe DMABC-FA6. **(a)** Cumulative distributions of fluorescence ON dwell times measured under different excitation laser powers. ON dwell times were defined as the durations of continuous fluorescence signals at fixed molecular positions. For all excitation conditions, the cumulative distributions were well described by single-exponential functions, indicating a single dominant dissociation process. **(b)** Observed dissociation rate constants (*k*_*obs*_) extracted from the fits in (a), plotted as a function of excitation laser power. *k*_*obs*_showed no systematic dependence on excitation intensity, indicating that probe dissociation is not limited by photophysical effects. The dissociation rate constant was therefore determined as the average value, yielding *k*_*off*_ = 33 s^−1^. **(c)** Cumulative distributions of fluorescence OFF dwell times measured at different probe concentrations, fitted with single-exponential functions. **(d)** Concentration dependence of the observed reappearance rate constant *k*_*obs*_. Linear fitting yields an association rate constant of *k*_*on*_ = 5.3 μ*M*^−1^ *s*^−1^. Error bars in b and d represent 90% confidence intervals obtained by bootstrap resampling (N_boot = 1000).

**Supplementary Fig. 7.**
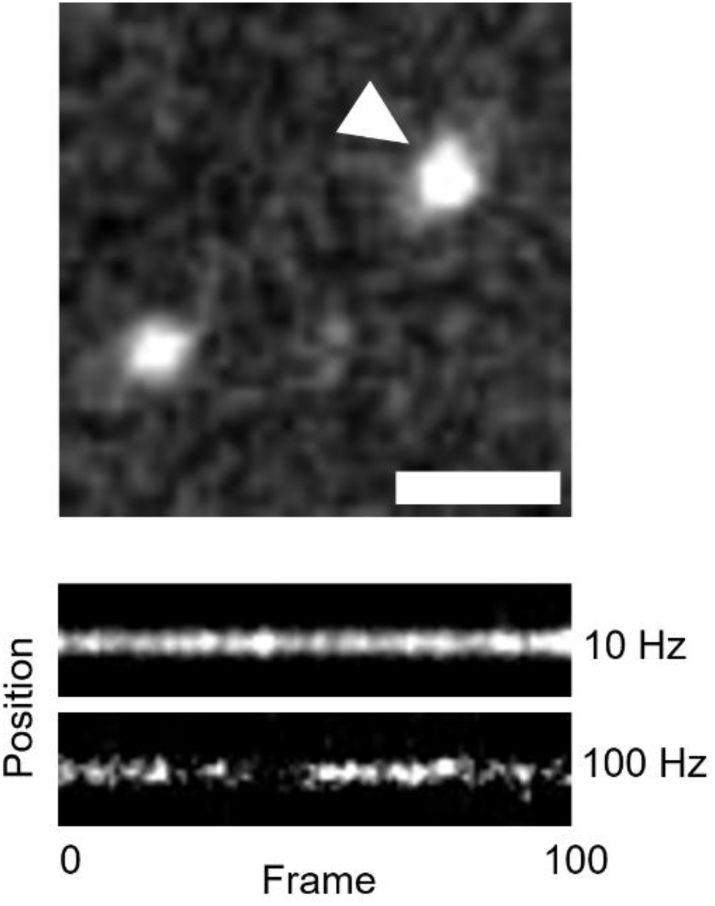
| Frame-rate dependence of EverGreen fluorescence dynamics. (Top) TIRF image indicating the single fluorescent spot selected for analysis (arrowhead). Scale bar, 1 μm. **(Bottom)** Kymographs of the same molecule recorded at 10 Hz and 100 Hz. While the signal appears continuous at 10 Hz, intermittent ON/OFF behavior becomes evident at higher temporal resolution, consistent with reversible probe binding dynamics.

**Supplementary Fig. 8.**
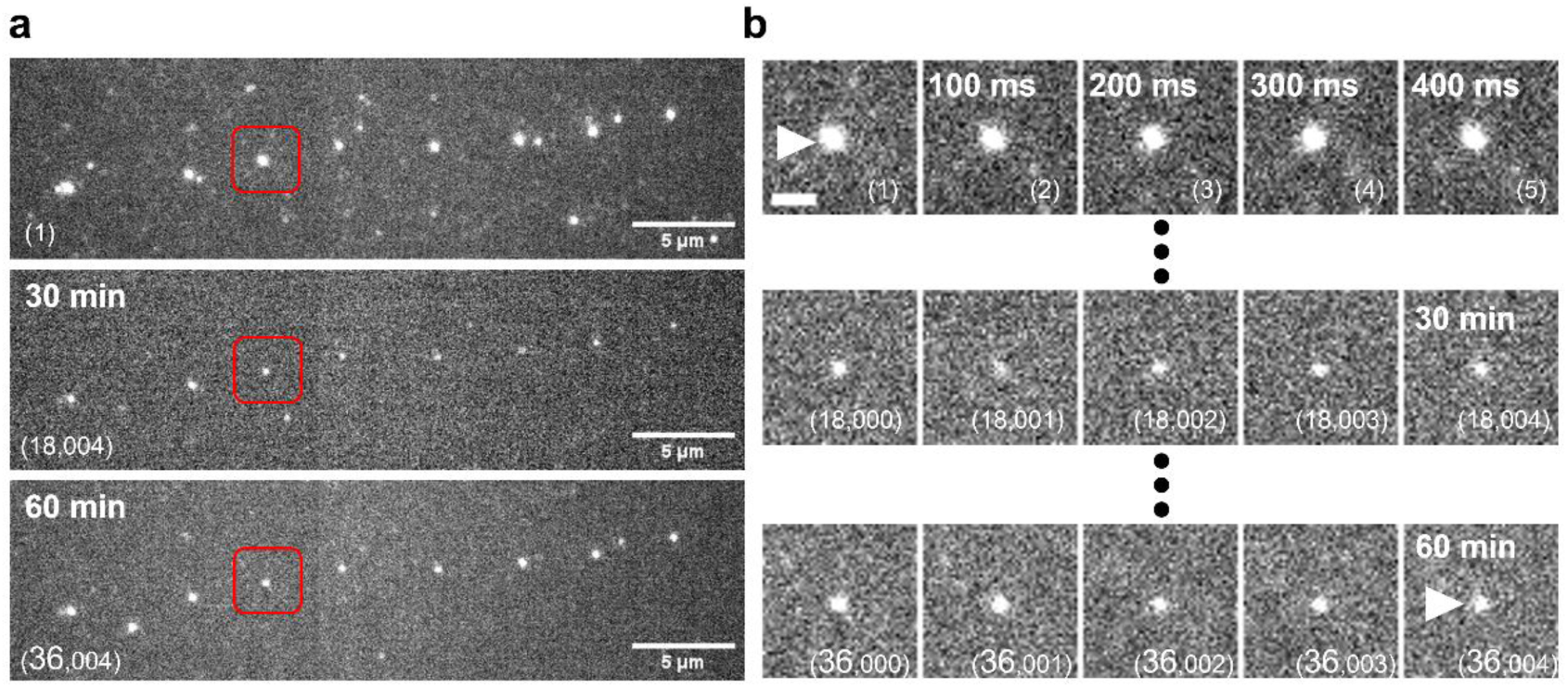
| Long-term single-molecule tracking by EverGreen without an oxygen scavenging system. **(a)** Wide-field views from continuous single-molecule fluorescence imaging acquired at the start of imaging, after 30 minutes, and after 60 minutes during continuous acquisition at 10 Hz under EverGreen labeling conditions in the absence of GLOX. Numbers in parentheses indicate the corresponding frame numbers. Red boxes denote the same region across time points. **(b)** Magnified view of the region outlined in red in (a). The white arrowhead indicates the same individual molecule observed throughout the experiment. Scale bars, 5 μm (a) and 1 μm (b).

**Supplementary Fig. 9.**
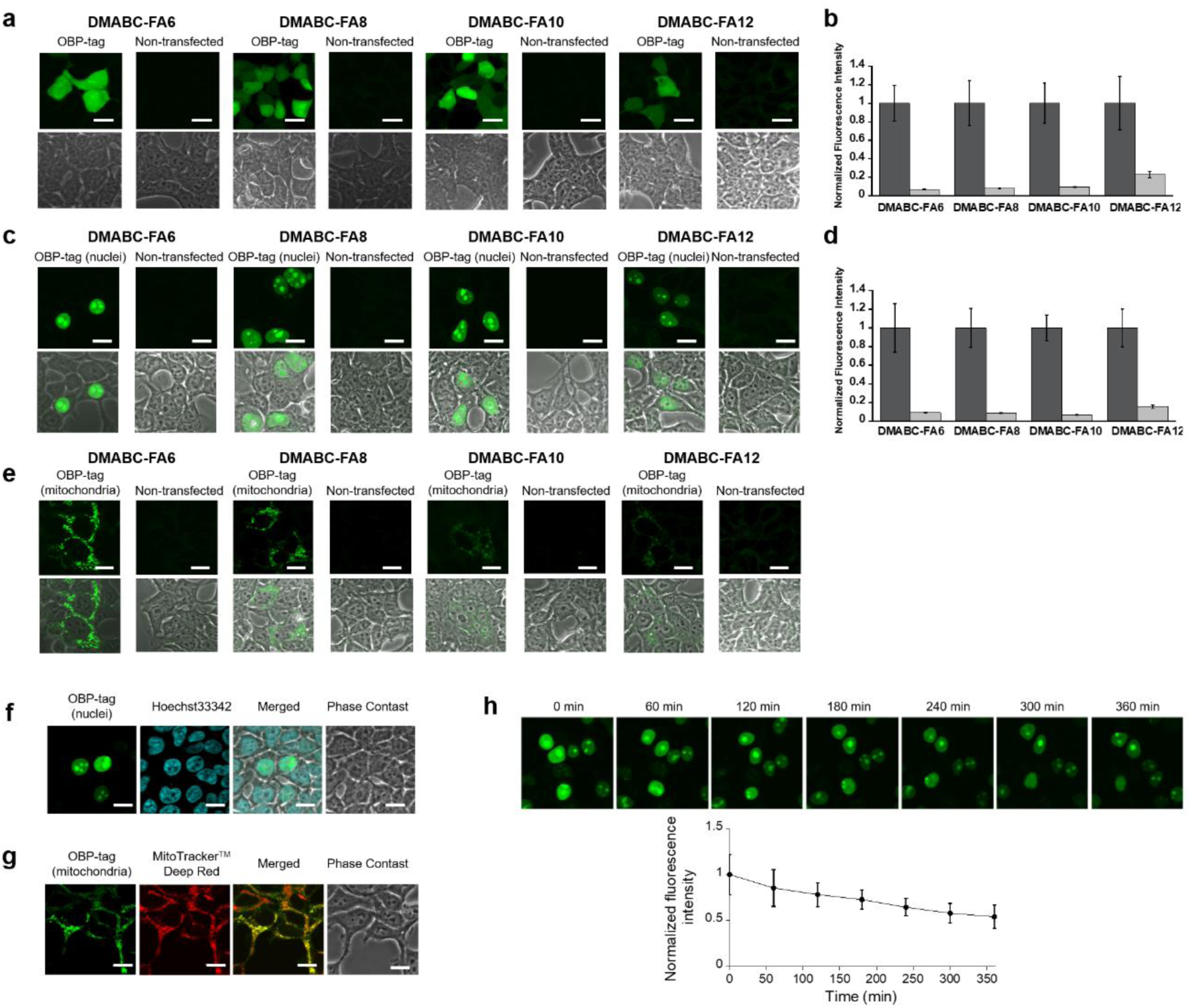
| Fluorescence imaging with EverGreen probes. **a.** Fluorescence images of live HEK293T cells expressing OBP-tag in cytosol or non-transfected cells after labeling with 1 μM DMABC-FA6, –FA8, –FA10, and –FA12. λ_*ex*_ = 473 nm, Scale bar: 20 μm. **b.** Analysis of fluorescence intensity in the OBP-tag-expressing and non-transfected cells. Error bars denote standard deviation (*N* = 9 cells). **c.** Fluorescence images of live HEK293T cells expressing OBP-tag in nuclei or non-transfected cells after labeling with 1 μM DMABC-FA6, –FA8, –FA10, and –FA12. λ_*ex*_ = 473 nm, Scale bar: 20 μm. **d.** Analysis of fluorescence intensity in the OBP-expressing and non-transfected cells. Error bars denote standard deviation (*N* = 9 cells). **e.** Fluorescence images of live HEK293T cells expressing OBP-tag in mitochondria or non-transfected cells after labeling with 1 μM DMABC-FA6, –FA8, –FA10, and –FA12. λ_*ex*_= 473 nm, Scale bar: 20 μm. **f.** Fluorescence images of live HEK293T cells expressing OBP-tag in nuclei after co-staining with DMABC-FA8 (1 μM) and Hoechst 33342 (0.4 μg/mL). λ_*ex*_= 405 and 473 nm. Scale bar: 20 μm. **g.** Fluorescence images of live HEK293T cells expressing OBP-tag in mitochondria after co-staining with DMABC-FA8 (1 μM) and MitoTracker^TM^ DeepRed (0.4 μg/mL). λ_*ex*_ = 473 nm and 635 nm. Scale bar: 20 μm. **h.** (top) Time-lapse fluorescence images of live HEK293T cells expressing OBP-tag in nuclei after labeling with DMABC-FA8. λ_*ex*_ = 473 nm. Scale bar: 20 μm. (bottom) Time course of mean fluorescence intensity. Error bars represent standard deviation (*N* = 18 cells).

**Supplementary Fig. 10.**
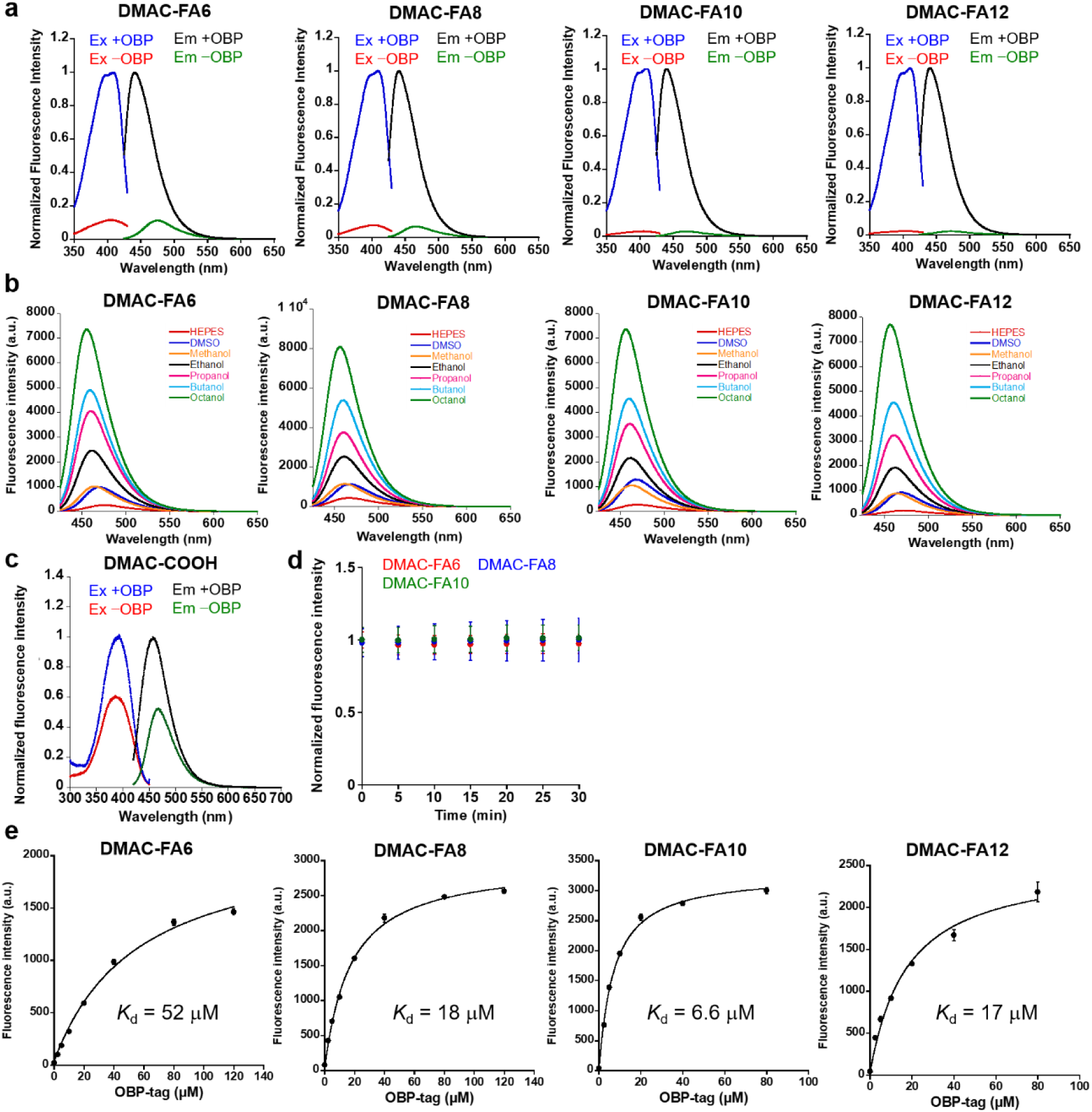
| Fluorescence and OBP-binding properties of DMAC probes. **a.** Fluorescence excitation and emission spectra of 1 μM DMAC-FA6, –FA8, –FA10, and – FA12 (0.2% DMSO) in the absence and presence of 50 μM OBP-tag in 20 mM HEPES buffer (pH = 7.4) containing 150 mM NaCl at 37 °C. λ_*ex*_ = 413 nm for emission spectra, λ_*em*_ = 465 nm for excitation spectra. **b.** Fluorescence spectra of 1 μM DMAC probes (0.2% DMSO) in different organic solvent and 20 mM HEPES buffer (pH = 7.4) containing 150 mM NaCl at 37 °C. λ_*ex*_ = 413 nm. **c.** Fluorescence excitation and emission spectra of 1 μM DMAC-COOH (0.2% DMSO) in the absence and presence of 50 μM OBP-tag in 20 mM HEPES buffer (pH = 7.4) containing 150 mM NaCl at 37 °C. λ_*ex*_= 403 nm for emission spectra, λ_*em*_ = 465 nm for excitation spectra. **d.** Photostability of DMAC probes under continuous light illumination (405 nm, 4 mW/cm^2^). The sample solution contains 1 μM probe (0.2% DMSO) in the presence OBP-tag (50% labeling condition) in 20 mM HEPES buffer (pH = 7.4) and 150 mM NaCl at 37 °C. λ_*ex*_ = 413 nm. Error bars denote standard deviation (*N* = 3). **e.** Fluorescence titration of DMAC-FA6, –FA8, –FA10, and – FA12 for different concentrations of OBP-tag at 37 °C. λ_*ex*_= 413 nm. Error bars denote standard deviation (*N* = 3).

**Supplementary Fig. 11.**
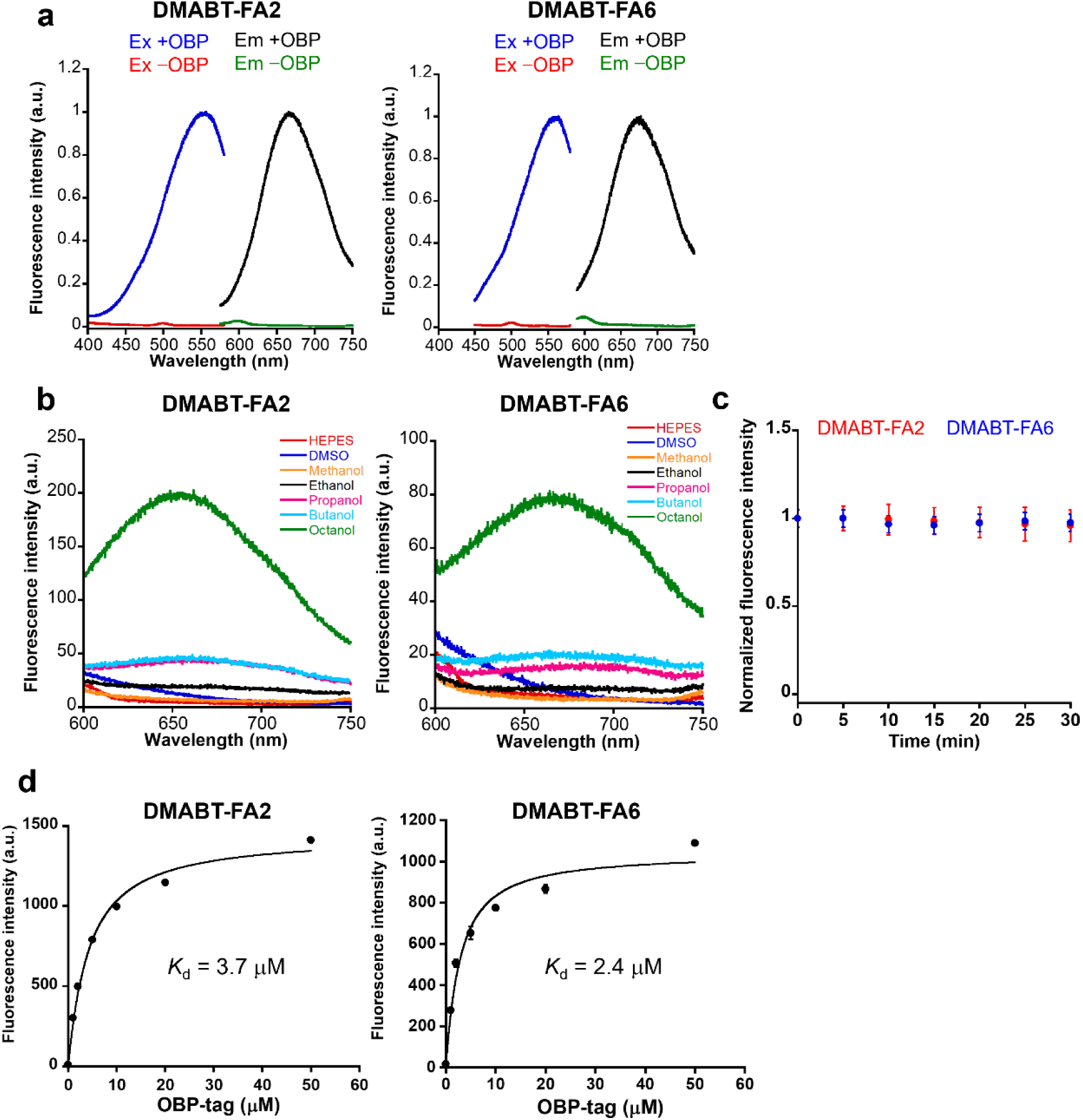
| Fluorescence and OBP-binding properties of DMABT probes. **a.** Fluorescence excitation and emission spectra of 1 μM DMABT-FA2 and –FA6 (0.1% DMSO) in the absence and presence of 5 μM OBP-tag in 20 mM HEPES buffer (pH = 7.4) containing 150 mM NaCl at 37 °C. λ_*ex*_= 540 nm (emission spectra), λ_*em*_= 670 nm (excitation spectra). **b.** Fluorescence spectra of 1 μM DMABT-FA2 and –FA6 (0.1% DMSO) in different organic solvent at 37 °C. λ_*ex*_ = 540 nm. **c.** Photostability of DMABT probes under continuous light illumination (550 nm, 4 mW/cm^2^). The sample solution contains 1 μM probe (0.1% DMSO) in the presence OBP-tag (50% labeling condition) in 20 mM HEPES buffer (pH = 7.4) and 150 mM NaCl at 37 °C. λ_*ex*_ = 540 nm. Error bars denote standard deviation (*N* = 3). **d.** Fluorescence titration of DMABT-FA2 and –FA6 for different concentrations of OBP-tag at 37 °C. λ_*ex*_= 540 nm. Error bars denote standard deviation (*N* = 3).

**Supplementary Fig. 12.**
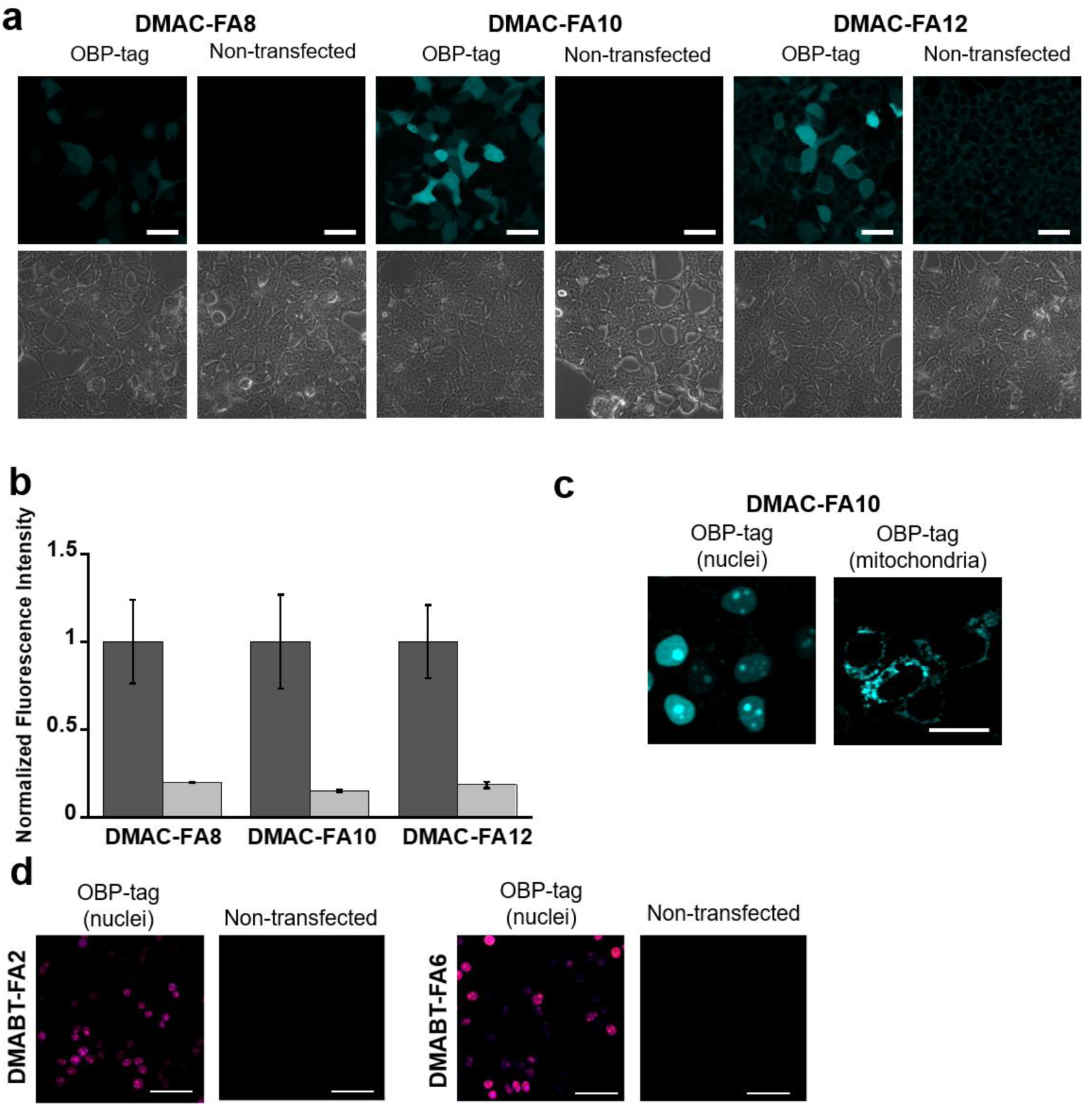
| Fluorescence imaging with DMAC and DMABT probes. **a.** Fluorescence images of live HEK293T cells expressing OBP-tag in cytosol after labeling with 2 μM DMAC-FA8, FA10, and FA12. λ_*ex*_= 405 nm, Scale bar: 50 μm. **b.** Quantitative analysis of fluorescence intensity between OBP-expressing and non-expressing cells. Error bars denote standard deviation (*N* = 9 cells). **c.** Fluorescence images of live HEK293T cells expressing OBP-tag in nuclei and mitochondria after labeling with 1 μM DMAC-FA10. λ_*ex*_= 405 nm, Scale bar: 50 μm. **d.** Fluorescence images of live HEK293T cells expressing OBP-tag in nuclei after labeling with 1 μM DMABT-FA2 and –FA6. λ_*ex*_= 635 nm, Scale bar: 50 μm.

**Supplementary Fig. 13.**
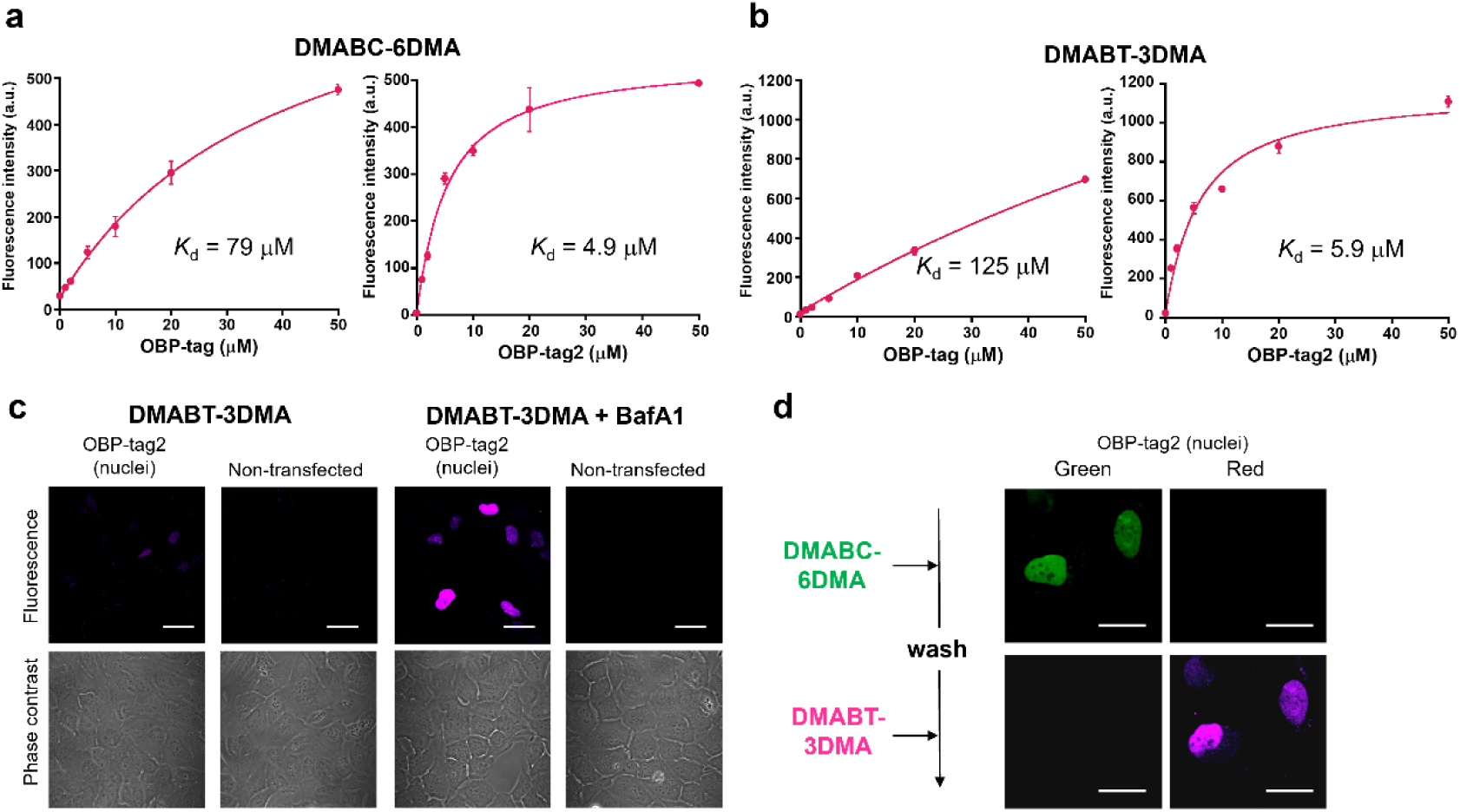
| Labeling properties of probes for OBP-tag2. **a.** Fluorescence titration of DMABC-6DMA for different concentrations of OBP-tag (left) and OBP-tag2 (right). The probes were excited at 455 nm and the fluorescence intensity at 540 nm was measured at 37 °C. Error bars denote standard deviation (*N* = 3). **b.** Fluorescence titration of DMABT-3DMA for different concentrations of OBP-tag (left) and OBP-tag2 (right) at 37 °C. λ_*ex*_ = 540 nm. Error bars denote standard deviation (*N* = 3). **c.** Fluorescence images of live U2OS cells expressing OBP-tag2 in nuclei after labeling with 1 μM DMABT-3DMA in the absence and presence of 0.1 μM Bafilomycin A1. λ_*ex*_ = 635 nm, Scale bar: 50 μm. **d.** Probe exchange by sequential labeling with DMABC-6DMA and DMABT-3DMA probes. Live U2OS cells expressing OBP-tag2 in nuclei were imaged after labeling with 1 μM DMABC-6DMA (top) and 1 μM DMABT-3DMA after washing (bottom). 0.1 μM Bafilomycin A1 was added prior to labeling. λ_*ex*_= 473 nm and 635 nm, Scale bar: 50 μm.

**Supplementary Table 1.** | Kinetic and fluorogenic parameters of exchangeable protein labeling systems.

| No. | System | Tag–fluorogen | $K_d$<br>($\mu\text{M}$) | $k_{\text{on}}$<br>($\mu\text{M}^{-1} \text{s}^{-1}$) | $k_{\text{off}}$<br>($\text{s}^{-1}$) | $\text{CR}^a$ | $k_{\text{on}} \times \text{CR}$<br>($\mu\text{M}^{-1} \text{s}^{-1}$) | Ref. |
| --- | --- | --- | --- | --- | --- | --- | --- | --- |
| 1 | FAP | HL1–TO1 | 0.3 | $0.033^b$ | 0.01 | $2,600^d$ | 86 | [12,39] |
| 2 | FAST | Y-FAST–HMBR | 0.13 | 63 | 6.3 | $\sim 1,000^e$ | $\sim 63,000$ | [13,22] |
| 3 | Blc | DiB1–M739 | 0.11 | $0.1^b$ | 0.01 | 52 | 5 | [14] |
| 4 | Blc | DiB2–M739 | 4.0 | $0.025^b$ | 0.1 | 64 | 1.6 | [14] |
| 5 | Blc | DiB3–M739 | 9.1 | $0.033^b$ | 0.3 | 11 | 0.4 | [14] |
| 6 | xHalo | dHaloTag7–MaP618-Hy4 | 0.27 | 20 | 5 | 37 | 740 | [15] |
| 7 | reHaloTag<sup>c</sup> | reHaloTagF–MaP618 | 0.066 | 1.1 | 0.073 | $\sim 30^f$ | $\sim 33$ | [16] |
| 8 | srTag | FKBP<sub>F36L</sub>–ZCD-1 | 0.049 | 0.16 | 0.008 | 86 | 14 | [17] |
| 9 | ReTag1 | ReTag1–MaP655-TMP | 2.8 | 0.006 | 0.011 | 161 | 1.0 | [18] |
| 10 | SiR-tag | SiR-tag–SiR | 0.048 | 140 | 6.7 | 13 | 1,820 | [19] |
| 11 | **EverGreen** | **OBP–DMABC- FA8** | **8.0** | **3.0** | **24** | **350** | **1,050** | This work |

<sup>a</sup>Fluorogenic contrast ratio ( $F_{\text{bound}}/F_{\text{free}}$ ), determined by *in vitro* spectroscopy unless otherwise noted.
<sup>b</sup>Not directly measured; estimated as $k_{\text{off}}/K_d$ .
<sup>c</sup>Covalent ester formation/hydrolysis mechanism.
<sup>d</sup>Fluorogenicity of HL1.0.1–TO1 complex <sup>12</sup>; kinetics measured for HL1 variant <sup>39</sup>.
<sup>e</sup>Based on corrected quantum yield of HMBR <sup>22</sup>.
<sup>f</sup>Estimated from HaloTag7–MaP618 fluorogenicity <sup>15</sup>, as reHaloTag shares the same binding pocket.

**Supplementary Table 2.**
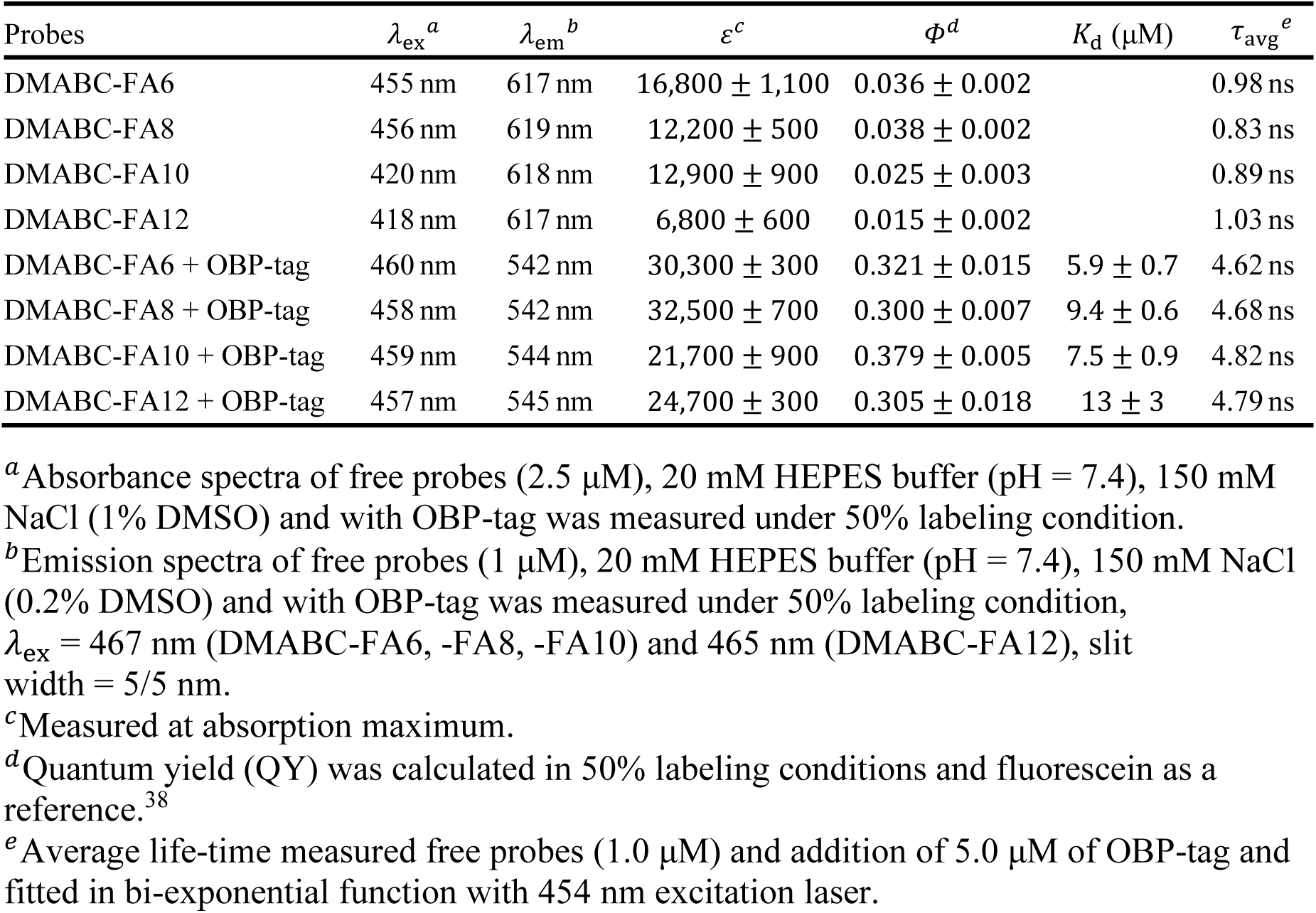
| Photophysical properties of DMABC-FA probes.

| Probes | $\lambda_{\text{ex}}^a$ | $\lambda_{\text{em}}^b$ | $\varepsilon^c$ | $\Phi^d$ | $K_d$ ($\mu\text{M}$) | $\tau_{\text{avg}}^e$ |
| --- | --- | --- | --- | --- | --- | --- |
| DMABC-FA6 | 455 nm | 617 nm | $16,800 \pm 1,100$ | $0.036 \pm 0.002$ | | 0.98 ns |
| DMABC-FA8 | 456 nm | 619 nm | $12,200 \pm 500$ | $0.038 \pm 0.002$ | | 0.83 ns |
| DMABC-FA10 | 420 nm | 618 nm | $12,900 \pm 900$ | $0.025 \pm 0.003$ | | 0.89 ns |
| DMABC-FA12 | 418 nm | 617 nm | $6,800 \pm 600$ | $0.015 \pm 0.002$ | | 1.03 ns |
| DMABC-FA6 + OBP-tag | 460 nm | 542 nm | $30,300 \pm 300$ | $0.321 \pm 0.015$ | $5.9 \pm 0.7$ | 4.62 ns |
| DMABC-FA8 + OBP-tag | 458 nm | 542 nm | $32,500 \pm 700$ | $0.300 \pm 0.007$ | $9.4 \pm 0.6$ | 4.68 ns |
| DMABC-FA10 + OBP-tag | 459 nm | 544 nm | $21,700 \pm 900$ | $0.379 \pm 0.005$ | $7.5 \pm 0.9$ | 4.82 ns |
| DMABC-FA12 + OBP-tag | 457 nm | 545 nm | $24,700 \pm 300$ | $0.305 \pm 0.018$ | $13 \pm 3$ | 4.79 ns |

<sup>a</sup>Absorbance spectra of free probes (2.5 $\mu\text{M}$ ), 20 mM HEPES buffer (pH = 7.4), 150 mM NaCl (1% DMSO) and with OBP-tag was measured under 50% labeling condition.
<sup>b</sup>Emission spectra of free probes (1 $\mu\text{M}$ ), 20 mM HEPES buffer (pH = 7.4), 150 mM NaCl (0.2% DMSO) and with OBP-tag was measured under 50% labeling condition, $\lambda_{\text{ex}} = 467$ nm (DMABC-FA6, -FA8, -FA10) and 465 nm (DMABC-FA12), slit width = 5/5 nm.
<sup>c</sup>Measured at absorption maximum.
<sup>d</sup>Quantum yield (QY) was calculated in 50% labeling conditions and fluorescein as a reference.<sup>38</sup>
<sup>e</sup>Average life-time measured free probes (1.0 $\mu\text{M}$ ) and addition of 5.0 $\mu\text{M}$ of OBP-tag and fitted in bi-exponential function with 454 nm excitation laser.

**Supplementary Table 3.** | Photophysical properties of DMAC-FA probes.

| Probes | $\lambda_{\text{ex}}^a$ | $\lambda_{\text{em}}^b$ | $\varepsilon^c$ | $\Phi^d$ | $K_d$ ($\mu\text{M}$) |
| --- | --- | --- | --- | --- | --- |
| DMAC-FA6 | 423 nm | 470 nm | $38,500 \pm 900$ | $0.030 \pm 0.001$ | |
| DMAC-FA8 | 421 nm | 465 nm | $38,000 \pm 2,100$ | $0.048 \pm 0.004$ | |
| DMAC-FA10 | 422 nm | 469 nm | $33,200 \pm 2,100$ | $0.038 \pm 0.006$ | |
| DMAC-FA12 | 422 nm | 469 nm | $17,700 \pm 800$ | $0.053 \pm 0.006$ | |
| DMAC-FA6 + OBP-tag | 416 nm | 444 nm | N.D. | N.D. | $52 \pm 6$ |
| DMAC-FA8 + OBP-tag | 415 nm | 444 nm | $46,000 \pm 500$ | $0.188 \pm 0.026$ | $18 \pm 1$ |
| DMAC-FA10 + OBP-tag | 416 nm | 439 nm | $34,900 \pm 200$ | $0.237 \pm 0.014$ | $6.6 \pm 0.5$ |
| DMAC-FA12 + OBP-tag | 416 nm | 439 nm | $39,500 \pm 2,600$ | $0.210 \pm 0.032$ | $17 \pm 3$ |

<sup>a</sup>Absorbance spectra of free probes (2.5 $\mu\text{M}$ ), 20 mM HEPES buffer (pH = 7.4), 150 mM NaCl (0.5% DMSO) and with OBP-tag was measured under 50% labeling condition.
<sup>b</sup>Emission spectra of free probes (1 $\mu\text{M}$ ), 20 mM HEPES buffer (pH = 7.4), 150 mM NaCl (0.2% DMSO) and with OBP-tag was measured under 50% labeling condition, $\lambda_{\text{ex}} = 413$ nm, slit width = 5/2.5 nm.
<sup>c</sup>Measured at absorption maximum.
<sup>d</sup>Quantum yield (QY) was calculated in 50% labeling conditions and Coumarin 153 as a reference.<sup>38</sup>

**Supplementary Table 4.**
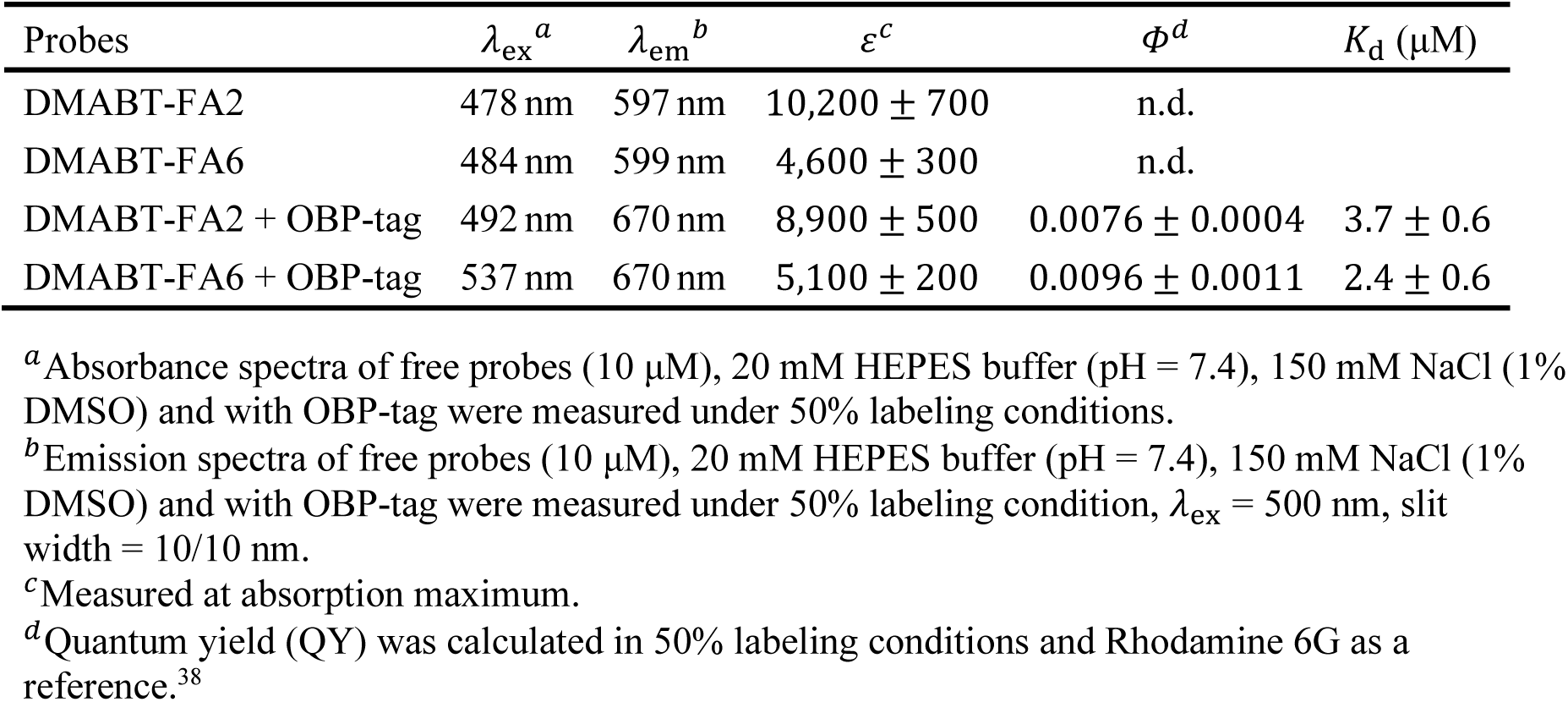
| Photophysical properties of DMABT-FA probes.

**Supplementary Table 5.** | Photophysical properties of DMABC-6DMA and DMABT-3DMA.

| Probes | $\lambda_{\text{ex}}^a$ | $\lambda_{\text{em}}^b$ | $\varepsilon^c$ | $\Phi^d$ | $K_d$ ($\mu\text{M}$) |
| --- | --- | --- | --- | --- | --- |
| DMABC-6DMA | 450 nm | 613 nm | $3,900 \pm 200$ | $0.019 \pm 0.002$ | |
| DMABT-3DMA | 478 nm | 597 nm | $4,300 \pm 300$ | n.d. | |
| DMABC-6DMA + OBP-tag2 | 454 nm | 535 nm | $6,600 \pm 100$ | $0.70 \pm 0.03$ | $4.9 \pm 0.6$ |
| DMABT-3DMA + OBP-tag2 | 540 nm | 659 nm | $5,700 \pm 100$ | $0.041 \pm 0.001$ | $5.9 \pm 1.3$ |

<sup>a</sup>Absorbance spectra of free probes (10 $\mu\text{M}$ ), 20 mM HEPES buffer (pH = 7.4), 150 mM NaCl (1% DMSO) and with OBP-tag was measured under 50% labeling condition.
<sup>b</sup>Emission spectra of free probes (10 $\mu\text{M}$ ), 20 mM HEPES buffer (pH = 7.4), 150 mM NaCl (1% DMSO) and with OBP-tag was measured under 50% labeling condition, $\lambda_{\text{ex}}$ = 450 nm (DMABC-6DMA), 500 nm (DMABT-3DMA), slit width = 5/5 (DMABC-6DMA), 10/10 (DMABT-3DMA) nm.
<sup>c</sup>Measured at absorption maximum.
<sup>d</sup>Quantum yield (QY) was calculated in 50% labeling conditions and fluorescein (DMABC-6DMA) or Rhodamine 6G (DMABT-3DMA) as a reference.<sup>38</sup>

**Supplementary Table 6.** | Binding constants of DMABC-FA6 and DMABT-3DMA for OBP-tag and OBP-tag2.

| Probes | $K_d$ (μM), OBP-tag | $K_d$ (μM), OBP-tag2 |
| --- | --- | --- |
| DMABC-FA6 | $5.9 \pm 0.7$ | $62 \pm 19$ |
| DMABT-3DMA | $124 \pm 24$ | $5.9 \pm 1.3$ |

