## Supplementary Information for "Kinetic design of reversible probe exchange enables continuous single-molecule tracking beyond the photobleaching limit"

### Table of Contents

**1. Synthetic Procedures** ........ **3–15**

**2. NMR Spectra** ........ **16–42**

**3. References** ........ **43**

#### 1. Synthetic Procedures

##### 1.1. Synthesis of benzocoumarin probes

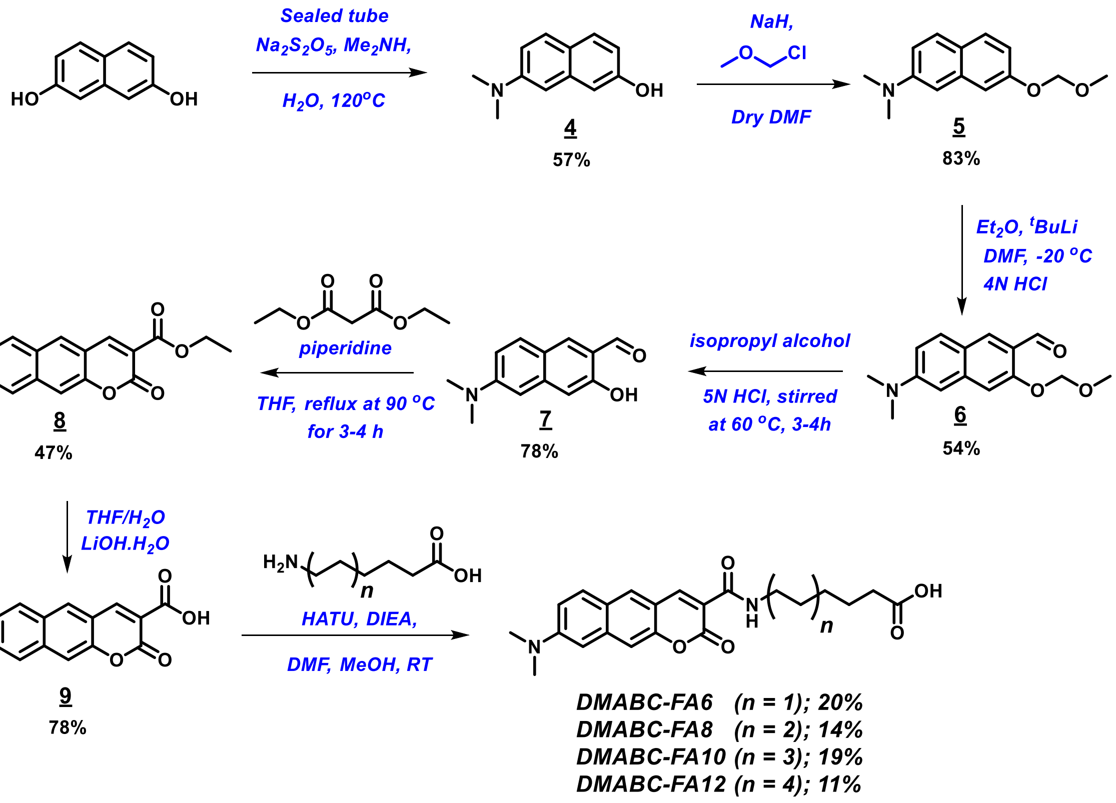


The synthesis of benzocoumarin dye **9** was reported earlier.[S1] We synthesized with literature reports and modified reaction conditions according to our set-up condition.

**7-(dimethylamino)naphthalen-2-ol (4)**

Dimethylamine (6.3 mL, 94 mmol) was added to a mixture of 2,7-dihydroxynaphthalene (3.00 g, 18.7 mmol), sodium metabisulfite (7.12 g, 37.5 mmol), and $H_2O$ (10 mL) in a seal-tube. The reaction mixture was stirred at 120 °C for 8-9 h (be careful and mix the reaction before heated). The reaction mixture was cooled to room temperature, and dichloromethane (100 mL) was added to the reaction mixture. The organic layer was washed with brine, dried over anhydrous $Na_2SO_4$, and concentrated. The residue was purified by silica gel column chromatography (eluent: 20% EtOAc in hexane) to afford compound **4** as a white solid (2.0 g, 57%).

**$^1$H NMR ($CDCl_3$, 500 MHz)**: δ 7.62–7.57 (m, 2H), 7.00 (dd, $J$ = 9.0 Hz, $J$ = 2.5 Hz, 1H), 6.94 (d, $J$ = 2.5 Hz 1H), 6.80 (dd, $J$ = 9.0 Hz, $J$ = 2.5 Hz, 1H), 6.75 (d, $J$ = 2.5 Hz, 1H), 4.94 (s, 1H), 3.02 (s, 6H) ppm.

**$^{13}$C NMR ($CDCl_3$, 125 MHz)**: δ 153.8, 149.2, 136.2, 129.4, 128.6, 122.4, 114.2, 113.7, 108.0, 105.2, 40.9 ppm.

**HRMS (FAB+)** ($m/z$): calcd $[M]^+$ for $C_{12}H_{13}NO$ 187.0997; found, 187.0995.

**7-(methoxymethoxy)-*N*,*N*-dimethylnaphthalen-2-amine (5)**

To a solution of compound **4** (1.0 g, 5.3 mmol) was added in the mixture of dry DMF (10 mL) and NaH (235 mg, 5.9 mmol, 60% NaH) at 0 °C. The resulting mixture was stirred for 1h at 0 °C. Then chloromethyl methyl ether (0.40 mL, 5.3 mmol) was added dropwise to the reaction mixture at the same temperature. The mixture was stirred at room temperature for overnight and then treated with water (50 mL). The two layers were separated, and the aqueous layer was extracted with EtOAc (100 mL). The combined organic extracts were washed with brine, dried over anhydrous $Na_2SO_4$, and concentrated. The residue was purified by silica gel column chromatography (eluent: 10% EtOAc in hexane) to afford compound **5** as a white solid (1.0 g, 83%).

**$^1$H NMR ($CDCl_3$, 500 MHz)**: δ 7.62–7.59 (m, 2H), 7.22 (d, *J* = 2.0 Hz, 1H), 7.02 (dd, *J* = 9.0 Hz, *J* = 2.5 Hz 1H), 6.93 (dd, *J* = 9.0 Hz, *J* = 2.5 Hz, 1H), 6.84 (d, *J* = 2.5 Hz, 1H), 5.27 (s, 2H), 3.52 (s, 3H) 3.03 (s, 6H) ppm.

**$^{13}$C NMR ($CDCl_3$, 125 MHz)**: δ 155.6, 149.1, 136.0, 129.0, 128.4, 122.9, 114.9, 114.5, 108.5, 105.9, 94.5, 56.0, 40.9 ppm.

**HRMS (FAB+)** (*m/z*): calcd $[M]^+$ for $C_{14}H_{17}NO_2$ 231.1259; found, 231.1256.

**6-(dimethylamino)-3-(methoxymethoxy)-2-naphthaldehyde (6)**

To a solution of compound **5** (1.0 g, 4.3 mmol) in $Et_2O$ (20 mL) cooled to -20 °C was added *t*-BuLi (1.7 M in pentane, 4.1 mL, 6.9 mmol) dropwise over 20 min. The resulting mixture was stirred at -20 °C for 2 h, which was treated with DMF (15 mL) dropwise. The mixture was stirred at -20 °C for 60 min and then treated with 4 N HCl (10 mL) slowly under vigorous stirring. The resulting two-phase system was stirred for 30 min. EtOAc (100 mL) was added and the organic layer was separated, washed with a saturated $NaHCO_3$ solution (100 mL), and brine (50 mL); it was dried ($Na_2SO_4$), and concentrated under reduced pressure to give a yellow solid. The residue was purified by silica gel column chromatography (eluent: 10% EtOAc in hexane) to afford compound **6** as a yellow solid (600 mg, 54%).

**$^1$H NMR ($CDCl_3$, 500 MHz)**: δ 10.46 (s, 1H), 8.23 (s, 1H), 7.72 (d, *J* = 9.0 Hz, 1H), 7.22 (s, 1H), 7.02 (dd, *J* = 9.0 Hz, *J* = 2.5 Hz, 1H), 6.74 (d, *J* = 2.5 Hz, 1H), 5.38 (s, 2H), 3.56 (s, 3H) 3.10 (s, 6H) ppm.

**$^{13}$C NMR ($CDCl_3$, 125 MHz)**: δ 189.6, 156.0, 150.7, 139.7, 131.2, 130.9, 122.2, 121.3, 114.8, 107.6, 104.3, 94.7, 56.4, 40.3 ppm.

**HRMS (FAB+)** (*m/z*): calcd $[M]^+$ for $C_{15}H_{17}NO_3$ 259.1208; found, 259.1211.

**6-(dimethylamino)-3-hydroxy-2-naphthaldehyde (7)**

To a solution of compound **6** (600 mg, 2.3 mmol) in isopropyl alcohol (25 mL) was added 5 M HCl (10 mL). The reaction mixture was stirred at 60 °C for 4 h. After being cooled to room temperature, isopropyl alcohol was removed under reduced pressure, and then EtOAc (100 mL) was added to the residue. The organic layer was washed with brine, dried over anhydrous $Na_2SO_4$, and concentrated. The residue was purified by silica gel column chromatography (eluent: 20% EtOAc in hexane) to afford compound **7** as a yellow solid (390 mg, 78%).

**$^1$H NMR ($CDCl_3$, 500 MHz)**: δ 10.52 (s, 1H), 9.89 (d, *J* = 0.5 Hz, 1H), 7.90 (s, 1H), 7.68 (d, *J* = 9.0 Hz, 1H), 7.00-6.97 (m, 2H), 6.65 (d, *J* = 2.5 Hz, 1H), 3.12 (s, 6H) ppm.

**$^{13}$C NMR ($CDCl_3$, 125 MHz)**: δ 195.3, 156.8, 151.4, 140.6, 137.8, 130.9, 120.7, 119.0, 114.2, 108.8, 103.2, 40.3 ppm.
**ESI-MS (*m*/*z*)**: calcd $[M+H]^+$ for $C_{13}H_{13}NO_2$ 216.10; found, 216.05.

**Ethyl 8-(dimethylamino)-2-oxo-2H-benzo[*g*]chromene-3-carboxylate (8)**

To a solution of compound **7** (200 mg, 0.93 mmol) and diethylmalonate (151 μL, 1.0 mmol) at room temp under $N_2$ atmosphere, piperidine (40 μL) was added, and the resulting solution was stirred 90 ℃ for 3 h. Then, cool the reaction mixture solid crystal will appear upon cooling. Collect the solid crystal and wash with cold THF and dried in a high vacuum to afford compound **8** as a red solid (140 mg, 47%).

**$^{1}$H NMR ($CDCl_3$, 500 MHz)**: δ 8.58 (s, 1H), 7.91 (s, 1H), 7.76 (d, $J$ = 9.5 Hz, 1H), 7.39 (s, 1H), 7.13 (dd, $J$ = 9.5 Hz, $J$ = 2.5 Hz, 1H), 6.79 (d, $J$ = 2.5 Hz, 1H), 4.41 (q, $J$ = 7.0 Hz, 2H), 3.12 (s, 6H), 1.42 (t, $J$ = 7.0 Hz, 3H) ppm.

**$^{13}$C NMR ($CDCl_3$, 125 MHz)**: δ 163.7, 157.7, 151.8, 150.7, 149.4, 138.7, 130.8, 130.4, 123.5, 116.1, 114.9,114.3, 109.6, 103.9, 61.6, 40.3, 14.3 ppm.

**HRMS (FAB+)** (*m*/*z*): calcd $[M]^+$ for $C_{18}H_{17}NO_4$ 311.1158; found, 311.1163.

**8-(dimethylamino)-2-oxo-2H-benzo[*g*]chromene-3-carboxylic acid (9)**

To a solution of compound **8** (90 mg, 0.30 mmol) in THF/$H_2O$ (10 mL/10 mL) and LiOH.$H_2O$ (20 mg) was added to the solution. The resulting solution was stirred 90 ℃ for 2 h. Then, cool the reaction mixture and add 2 M HCl solution dropwise, and red color precipitates appeared. The precipitates were washed with water and cold THF and dried in a high vacuum to afford compound **9** as a red solid (66 mg, 78%).

**$^{1}$H NMR (DMSO-*d*$_6$, 500 MHz)**: δ 8.75 (s, 1H), 8.30 (s, 1H), 7.87 (d, $J$ = 9.5 Hz, 1H), 7.50 (s, 1H), 7.28 (dd, $J$ = 9.5 Hz, $J$ = 2.5 Hz, 1H), 6.96 (d, $J$ = 2.5 Hz, 1H), 3.11 (s, 6H) ppm.

**$^{13}$C NMR (DMSO-*d*$_6$, 125 MHz)**: δ 164.7, 158.2, 151.7, 151.1, 149.7, 138.6, 132.0, 130.9, 123.4, 116.8, 114.9, 114.4, 109.0, 104.0 ppm.

**HRMS (EI+)** (*m*/*z*): calcd $[M]^+$ for $C_{16}H_{13}NO_4$ 283.0845; found, 283.0849.

**6-(8-(dimethylamino)-2-oxo-2H-benzo[*g*]chromene-3-carboxamido)hexanoic acid (DMABC-FA6)**

The compound **9** (10 mg, 0.035 mmol) was dissolved in dry DMF (2 mL) and then HATU (16 mg, 0.042 mmol) and dry DIEA (13 μL, 0.071 mmol) were added to it. The reaction mixture was stirred for 1-1.5 h at 0 ºC under $N_2$ atmosphere. The compound 6-aminohexanoic acid (7 mg, 0.05 mmol) was dissolved in DMF/MeOH (2:1;1 mL) and sonicated for 30 min and then slowly added to the reaction mixture and stirred overnight at room temperature under $N_2$ atmosphere. After removing the solvent, the residue was purified by reversed-phase HPLC using ODS-3 column and eluted with $H_2O$/acetonitrile containing 0.1% formic acid to yield **DMABC-FA6** as an orange powder (2.8 mg; yield, 20%).

**$^{1}$H NMR (DMSO-*d*$_6$; 500 MHz)**: δ 8.84 (s,1H), 8.68 (t, $J$ = 5.5 Hz, 1 H), 8.35 (s, 1 H), 7.88 (d, $J$ = 9.5 Hz, 1H), 7.57 (s, 1 H), 7.29 (dd, $J$ = 9.0 Hz, $J$ = 2.5 Hz, 1H), 6.96 (d, $J$ = 2.5 Hz, 1H), 3.11 (s, 6H), 2.21 (t, $J$ = 7.0 Hz, 2H), 1.56-1.49 (m, 4H), 1.34-1.31 (m, 2H) ppm.

**$^{13}$C NMR (DMSO-*d*$_6$, 125 MHz)**: δ 174.9, 161.9, 161.6, 151.3, 150.9, 148.4, 138.4, 131.8, 130.8, 123.6, 116.9, 116.0, 114.7, 109.1, 104.0, 34.1, 29.3, 26.5, 24.7 ppm.

**HRMS (FAB+)** (*m*/*z*): calcd $[M+H]^+$ for $C_{22}H_{25}N_2O_5$ 397.1758; found 397.1764.

**8-(8-(dimethylamino)-2-oxo-2H-benzo[*g*]chromene-3-carboxamido)octanoic acid (DMABC-FA8)**

The compound **9** (10 mg, 0.035 mmol) was dissolved in dry DMF (2 mL) and then HATU (16 mg, 0.042 mmol) and dry DIEA (13 µL, 0.071 mmol) were added to it. The reaction mixture was stirred for 1-1.5 h at 0 °C under $N_2$ atmosphere. The compound 8-aminooctanoic acid (8 mg, 0.05 mmol) was dissolved in DMF/MeOH (2:1;1 mL) and sonicated for 30 min and then slowly added to the reaction mixture and stirred overnight at room temperature under $N_2$ atmosphere. After removing the solvent, the residue was purified by reversed-phase HPLC using ODS-3 column and eluted with $H_2O$/acetonitrile containing 0.1% formic acid to yield **DMABC-FA8** as an orange powder (2.8 mg; yield, 14%).

**$^1$H NMR (DMSO-*d₆*; 500 MHz)**: δ 8.83 (s,1H), 8.67 (t, *J* = 5.5 Hz, 1H), 8.35 (s, 1H), 7.88 (d, *J* = 9.5 Hz, 1H), 7.57 (s, 1H), 7.29 (dd, *J* = 9.5 Hz, *J* = 2.5 Hz, 1H), 6.96 (d, *J* = 2.0 Hz, 1H), 3.11 (s, 6H), 2.20 (t, *J* = 7.0 Hz, 2 H), 1.56-1.49 (m, 4 H), 1.30 (br, 6 H) ppm.

**$^{13}$C NMR (DMSO-$d_6$, 125 MHz)**: δ 175.0, 161.9, 161.6, 151.3, 150.9, 148.4, 138.4, 131.8, 130.8, 123.6, 116.9, 116.0, 114.8, 109.1, 104.0, 34.1, 29.5, 29.0, 28.9, 26.8, 24.9 ppm.

**HRMS (FAB+)** (*m*/*z*): calcd $[M+H]^+$ for $C_{24}H_{29}N_2O_5$ 425.2071; found 425.2069.

**10-(8-(dimethylamino)-2-oxo-2H-benzo[*g*]chromene-3-carboxamido)decanoic acid (DMABC-FA10)**

The compound **9** (10 mg, 0.035 mmol) was dissolved in dry DMF (2 mL) and then HATU (16 mg, 0.042 mmol) and dry DIEA (13 µL, 0.071 mmol) were added to it. The reaction mixture was stirred for 1-1.5 h at 0 °C under $N_2$ atmosphere. The compound 10-aminodecanoic acid (12 mg, 0.064 mmol) was dissolved in DMF/MeOH (2:1;1 mL) and sonicated for 30 min and then slowly added to the reaction mixture and stirred overnight at room temperature under $N_2$ atmosphere. After removing the solvent, the residue was purified by reversed-phase HPLC using ODS-3 column and eluted with $H_2O$/acetonitrile containing 0.1% formic acid to yield **DMABC-FA10** as an orange powder (3.1 mg; yield, 19%).

**$^1$H NMR (DMSO-$d_6$; 500 MHz)**: δ 8.84 (s,1H), 8.68 (t, *J* = 5.5 Hz, 1H), 8.35 (s, 1H), 7.88 (d, *J* = 9.5 Hz, 1H), 7.57 (s, 1 H), 7.29 (dd, *J* = 9.5 Hz, *J* = 2.5 Hz, 1H), 6.96 (d, *J* = 2.5 Hz, 1H), 3.11 (s, 6H), 2.13 (br, 2H), 1.54-1.48 (m, 4H), 1.30-1.26 (m, 10H) ppm.

**$^{13}$C NMR (DMSO-$d_6$, 125 MHz)**: δ 175.0, 161.9, 161.6, 151.3, 150.9, 148.4, 138.4, 131.8, 130.8, 123.6, 116.9, 116.0, 114.7, 109.1, 104.0, 34.2, 29.5, 29.3, 29.2, 29.1, 29.0, 26.9, 25.0 ppm.

**HRMS (FAB+)** (*m*/*z*): calcd $[M]^+$ for $C_{26}H_{32}N_2O_5$ 452.2311; found 452.2312.

**12-(8-(dimethylamino)-2-oxo-2H-benzo[*g*]chromene-3-carboxamido)dodecanoic acid (DMABC-FA12)**

The compound **9** (10 mg, 0.035 mmol) was dissolved in dry DMF (2 mL) and then HATU (16 mg, 0.042 mmol) and dry DIEA (13 µL, 0.071 mmol) were added to it. The reaction mixture was stirred for 1-1.5 h at 0 °C under $N_2$ atmosphere. The compound 12-aminododecanoic acid (11 mg, 0.053 mmol) was dissolved in DMF/MeOH (2:1;1 mL) and sonicated for 30 min and then slowly added into the reaction mixture and stirred overnight at room temperature under $N_2$ atmosphere. After removing the solvent, the residue was purified by reversed-phase HPLC using ODS-3 column and eluted with $H_2O$/acetonitrile containing 0.1% formic acid to yield **DMABC-FA12** as an orange powder (1.9 mg; yield, 11%).

**$^1$H NMR (DMSO-$d_6$; 500 MHz)**: δ 8.83 (s,1H), 8.67 (t, $J$ = 5.5 Hz, 1H), 8.35 (s, 1H), 7.88 (d, $J$ = 9.5 Hz, 1H), 7.57 (s, 1H), 7.29 (dd, $J$ = 9.5 Hz, $J$ = 2.5 Hz, 1H), 6.96 (d, $J$ = 2.5 Hz, 1H), 3.11 (s, 6H), 2.16 (t, $J$ = 7.0 Hz, 2H), 1.54-1.45 (m, 4H), 1.34-1.31 (m, 14H) ppm.

**$^{13}$C NMR (DMSO-$d_6$, 125 MHz)**: δ 175.0, 161.9, 161.6, 151.3, 150.9, 148.4, 138.4, 131.8, 130.8, 123.6, 116.9, 116.0, 114.7, 109.1, 104.0, 34.3, 29.5, 29.4, 29.4, 29.2, 29.2, 29.1, 26.9, 25.1 ppm.

**HRMS (FAB+)** (*m*/*z*): calcd $[M]^+$ for $C_{28}H_{36}N_2O_5$ 480.2624; found 480.2622.

#### 1.2. Synthesis of coumarin probes

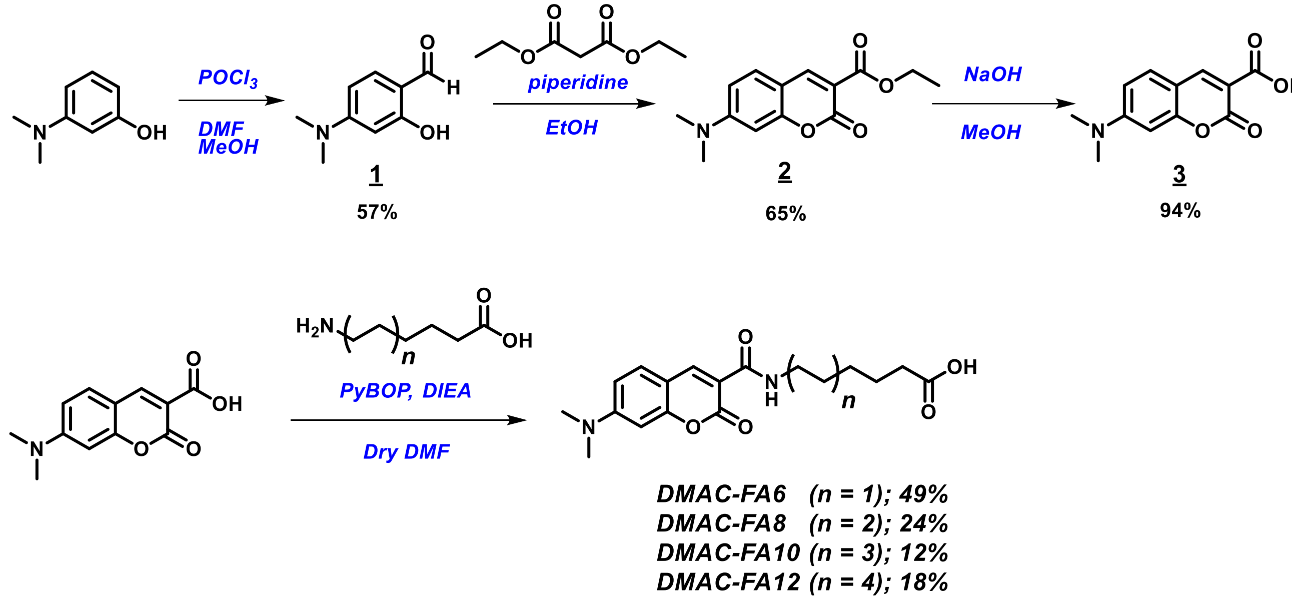


**6-(7-(dimethylamino)-2-oxo-2H-chromene-3-carboxamido)hexanoic acid (DMAC-FA6)**

The compound **3**[S2] (50 mg, 0.21 mmol) was dissolved in dry DMF/dry MeOH (10 mL/5.0 mL), and then PyBOP (73 mg, 0.14 mmol) and dry DIEA (49 µL, 0.28 mmol) were added to it. The reaction mixture was stirred for 1 h at 0 ºC under $N_2$ atmosphere. The compound 6-aminohexanoic acid (18 mg, 0.14 mmol) was slowly added to the reaction mixture and stirred overnight at room temperature under $N_2$ atmosphere. After removing the solvent, the residue was purified by reversed-phase HPLC using ODS-3 column and eluted with $H_2O$/acetonitrile containing 0.1% formic acid to yield **DMAC-FA6** as a yellow powder (24 mg, 49%).

**$^1$H NMR (500 MHz, CDCl$_3$)**: δ 8.84 (s, 1H), 8.74 (s, 1H), 7.47 (d, $J$ = 9.0 Hz, 1H), 6.71 (dd, $J$ = 2.0 Hz, $J$ = 8.5 Hz, 1H), 6.54 (d, $J$ = 2.0 Hz 1H), 3.43 (dd, $J$ = 7.0 Hz, $J$ = 13.0 Hz, 2H), 3.12 (s, 6H), 2.37 (t, $J$ = 7.5 Hz, 2H), 1.72-1.41 (br, 6H) ppm.

**$^{13}$C NMR (125 MHz, CDCl$_3$)**: δ 175.9, 161.6, 156.1, 153.3, 147.2, 129.9, 109.3, 108.0, 96.3, 39.4, 38.4, 32.5, 28.1, 25.4, 23.3 ppm.

**HRMS (ESI+)** (*m/z*): calcd $[M+H]^+$ for $C_{18}H_{23}N_2O_5$ 347.1601, found 347.1612.

**8-(7-(dimethylamino)-2-oxo-2H-chromene-3-carboxamido)octanoic acid (DMAC-FA8)**

The compound **3**[S1] (40 mg, 0.17 mmol) was dissolved in dry MeOH (15 mL) and then PyBOP (88 mg, 0.17 mmol) and dry DIEA (59 µL, 0.34 mmol) were added to it. The reaction mixture stirred for 1 h at 0 ºC under $N_2$ atmosphere. The compound 8-aminooctanoic acid (32 mg, 0.20 mmol) was slowly added to the reaction mixture and stirred overnight at room temperature under $N_2$ atmosphere. After removing the solvent, the residue was purified by reversed-phase HPLC using ODS-3 column and eluted with $H_2O$/acetonitrile containing 0.1% formic acid to yield **DMAC-FA8** as a yellow powder (15 mg, 24%).

**$^1$H NMR (500 MHz, $CDCl_3$)**: δ 8.83 (t, $J$ = 7.0 Hz, 1H), 8.73 (s, 1H), 7.46 (d, $J$ = 9.0 Hz, 1H), 6.68 (dd, $J$ = 2.5 Hz, $J$ = 9.0 Hz, 1H), 6.51 (d, $J$ = 2.5 Hz 1H), 3.43 (dd, $J$ = 7.0 Hz, $J$ = 13.0 Hz, 2H), 3.11 (s, 6H), 2.35 (t, $J$ = 7.5 Hz, 2H), 1.66-1.32 (br, 10H) ppm.
**$^{13}$C NMR (125 MHz, $CDCl_3$)**: δ 177.4, 163.0, 162.7, 157.1, 154.4, 148.2, 130.8, 110.9, 110.1, 108.7, 97.0, 40.2, 39.6, 33.7, 29.3, 28.8, 28.7, 26.7, 24.5 ppm.
**HRMS (ESI+)** (*m/z*): calcd $[M+H]^+$ for $C_{20}H_{27}N_2O_5$ 375.1914, found 375.1922.

**10-(7-(dimethylamino)-2-oxo-2H-chromene-3-carboxamido)decanoic acid (DMAC-FA10)**

The compound **3**[S1] (45 mg, 0.19 mmol) was dissolved in dry MeOH (18 mL) and then PyBOP (98 mg, 0.19 mmol) and dry DIEA (66 µL, 0.38 mmol) were added to it. The reaction mixture was stirred for 1 h at 0 °C under $N_2$ atmosphere. The compound 10-aminodecanoic acid (43 mg, 0.23 mmol) was slowly added to the reaction mixture and stirred overnight at room temperature under $N_2$ atmosphere. After removing the solvent, the residue was purified by reversed-phase HPLC using ODS-3 column and eluted with $H_2O$/acetonitrile containing 0.1% formic acid to yield **DMAC-FA10** as a yellow powder (9.1 mg, 12%).
**$^1$H NMR (500 MHz, $CDCl_3$)**: δ 8.84 (t, $J$ = 7.0 Hz, 1H), 8.74 (s, 1H), 7.46 (d, $J$ = 9.0 Hz, 1H), 6.68 (dd, $J$ = 2.5 Hz, $J$ = 9.0 Hz, 1H), 6.51 (d, $J$ = 2.5 Hz 1H), 3.43 (dd, $J$ = 7.0 Hz, $J$ = 13.0 Hz, 2H), 3.12 (s, 6H), 2.34 (t, $J$ = 7.5 Hz, 2H), 1.67-1.31 (br, 14H) ppm.
**$^{13}$C NMR (125 MHz, $CDCl_3$)**: δ 177.3, 163.1, 162.7, 157.2, 154.4, 148.2, 130.9, 110.8, 110.1, 108.7, 97.0, 40.2, 39.7, 33.7, 29.4, 29.0, 28.9, 28.8, 28.8, 26.7, 24.6 ppm.
**HRMS (ESI+)** (*m/z*): calcd $[M+H]^+$ for $C_{22}H_{31}N_2O_5$ 403.2227, found 403.2237.

**12-(7-(dimethylamino)-2-oxo-2H-chromene-3-carboxamido)dodecanoic acid (DMAC-FA12)**

The compound **3**[S1] (50 mg, 0.21 mmol) was dissolved in dry DMF/dry MeOH (5 mL/5 mL) and then PyBOP (109 mg, 0.21 mmol) and dry DIEA (73 µL, 0.42 mmol) were added to it. The reaction mixture stirred for 1 h at 0 °C under $N_2$ atmosphere. The compound 12-aminododecanoic acid (69 mg, 0.69 mmol) was slowly added to the reaction mixture and stirred overnight at room temperature under $N_2$ atmosphere. After removing the solvent, the residue was purified by reversed-phase HPLC using ODS-3 column and eluted with $H_2O$/acetonitrile containing 0.1% formic acid to yield **DMAC-FA12** as a yellow powder (16 mg, 18%).
**$^1$H NMR (500 MHz, $CDCl_3$)**: δ 8.83 (t, $J$ = 7.0 Hz, 1H), 8.75 (s, 1H), 7.46 (d, $J$ = 9.0 Hz, 1H), 6.67 (dd, $J$ = 2.5 Hz, $J$ = 9.0 Hz, 1H), 6.50 (d, $J$ = 2.5 Hz 1H), 3.11 (s, 6H), 3.43 (dd, $J$ = 7.0 Hz, $J$ = 13.0 Hz, 2H), 2.34 (t, $J$ = 7.5 Hz, 2H), 1.66-1.25 (br, 18H) ppm.
**$^{13}$C NMR (125 MHz, $CDCl_3$)**: δ 177.3, 163.0, 162.7, 157.2, 154.4, 148.3, 130.9, 110.9, 110.1, 108.7, 97.0, 40.2, 39.7, 33.7, 29.7, 29.5, 29.1, 29.1, 29.0, 28.8, 28.7, 26.8, 24.6 ppm.
**HRMS (ESI+)** (*m/z*): calcd $[M+H]^+$ for $C_{24}H_{35}N_2O_5$ 431.2540, found 431.2545.

#### 1.3. Synthesis of benzofuran-benzothiadiazole probes

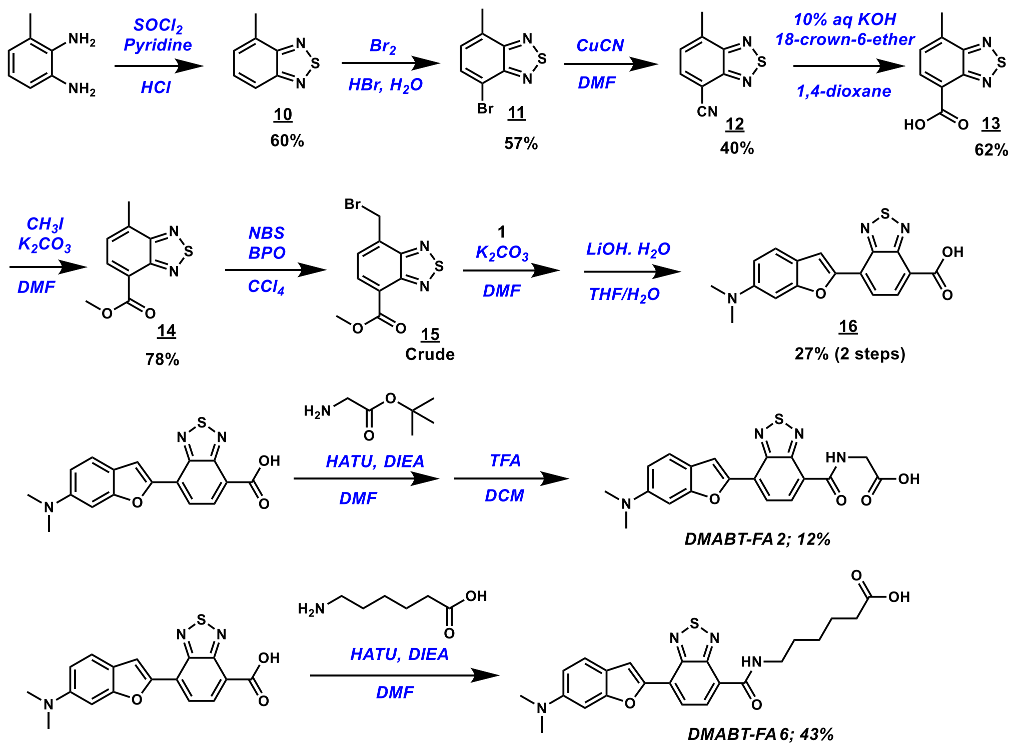


The synthesis of compound **11** was reported earlier.[S3] We synthesized compound **11** according to the reported synthetic procedure.

**7-methylbenzo[*c*][1,2,5]thiadiazole-4-carbonitrile (12)**

The compound **11** (1.0 g, 4.4 mmol) was dissolved in DMF (15 mL) and the reaction mixture was stirred at room temperature for 20 min. CuCN was added slowly (and carefully) to the reaction mixture and heated 150 °C for 16 h. Then the reaction was cooled down and stopped by addition of 15% $NH_3$ and stirred for 1 h. The resulting ppt was filtered and washed with DCM. The DCM layered was washed with brine and dried in $Na_2SO_4$. The DCM layer was evaporated and purified by silica gel column chromatography (eluent: 5% EtOAc in hexane) to afford compound **12** as a white solid (310 mg, 40%).

**$^1$H NMR ($CDCl_3$; 500 MHz)** δ 7.95 (d, $J$ = 7.0 Hz, 1H), 7.44-7.46 (m, 1H), 2.83 (s, 3H) ppm.

**$^{13}$C NMR ($CDCl_3$; 125 MHz)** δ 154.8, 152.8, 138.5, 136.1, 127.4, 115.6, 103.4, 18.5 ppm.

**HRMS (FAB+)** (*m*/*z*): calcd $[M+H]^+$ for $C_8H_6N_3S$ 176.0277; found: 176.0285

**7-methylbenzo[*c*][1,2,5]thiadiazole-4-carboxylic acid (13)**

To a solution of compound **12** (50 mg, 0.29 mmol) and 18-crown-6-ether (90 mg, 0.34 mmol) in 1,4-dioxane (2.5 mL), 10% NaOH aq. (2.5 mL) was added. After heating the mixture to 95°C for 16 h, cooling to room temperature and 2 M HCl was added to become pH = 1. The resulting mixture was extracted three times with DCM. The organic phases were washed with brine, dried over $Na_2SO_4$ and evaporated to dryness. The residue was purified by silica gel column chromatography (eluent: 5% MeOH in DCM) to afford compound **13** as a white solid (34 mg, 62%).

**$^1$H NMR (500 MHz, $CDCl_3$)** δ 13.20 (bs, 1H), 8.24 (d, $J$ = 7.0 Hz, 1H), 7.63 (d, $J$ = 7.0 Hz, 1H), 2.75 (s, 3H) ppm.

**$^{13}$C NMR (125 MHz, $CDCl_3$)** δ 164.9, 154.9, 151.0, 135.9, 133.0, 127.0, 120.9, 17.5 ppm.

**HRMS (FAB+)** (*m*/*z*): calcd $[M+Na]^+$ for $C_8H_6N_2NaO_2S$ 217.0042; found 217.0047

**methyl 7-methylbenzo[*c*][1,2,5]thiadiazole-4-carboxylate (14)**

To a solution of compound **13** (300 mg, 1.54 mmol) in DMF (10 mL) was added $K_2CO_3$ (200 mg, 1.85 mmol) and iodomethane (5 mL). The resulting mixture was stirred at 80 °C for overnight. After being cooled to room temperature, DMF was removed under reduced pressure. The residue was added to DCM (30 mL) and 2 M NaOH aq. (30 mL). The organic layer was separated, washed with brine (50 mL), dried over anhydrous $Na_2SO_4$, and concentrated under reduced pressure to afford compound **14** as a white solid (250 mg, 78%).

**$^1$H NMR (500 MHz, $CDCl_3$)** δ 8.24 (d, $J$ = 7.0 Hz, 1H), 7.38 (dd, $J$ = 7.0 Hz, $J$ = 1.0 Hz, 1H), 4.00 (s, 3H), 2.75 (d, $J$ = 1.0 Hz 3H) ppm.

**$^{13}$C NMR (125 MHz, $CDCl_3$)** δ 165.3, 156.0, 152.0, 137.8, 134.1, 127.2, 120.7, 52.6, 18.5 ppm.

**HRMS (FAB+)** (*m*/*z*): calcd $[M+Na]^+$ for $C_9H_8N_2NaO_2S$ 231.0199, found 231.0214

**methyl 7-(bromomethyl)benzo[*c*][1,2,5]thiadiazole-4-carboxylate (15)**

To a solution of compound **14** (110 mg, 0.53 mmol) in $CCl_4$ (3.5 mL) was added NBS (94 mg, 0.52 mmol), BPO (10 mg, 0.05 mmol) and 33 µL of 33% HBr in AcOH solution. The resulting mixture was stirred at 75 °C for overnight. After being cooled to room temperature, $CCl_4$ was removed under reduced pressure. The residue was added DCM (30 mL) and Water (30 mL). The organic layer was separated, washed with brine (50 mL); it was dried ($Na_2SO_4$), and concentrated under reduced pressure. The residue was purified by silica gel column chromatography (eluent: 10% EtOAc in hexane) to afford compound **15** as a crude.

**7-(6-(dimethylamino)benzofuran-2-yl)benzo[*c*][1,2,5]thiadiazole-4-carboxylic acid (16)**

Compound **15** (250 mg, 0.87 mmol), compound 1 (173 g, 1.0 mmol), and $K_2CO_3$ (0.70 g, 5.2 mmol) were placed in DMF (10 mL) at 125 °C stirred for 16 hours. The dark reaction mixture was diluted with water and extracted with ethyl acetate. The organic solvent was evaporated and the residue was used for the next step without further purification. The residue was dissolved in THF/$H_2O$ (1.5 mL/1.5 mL) and LiOH•$H_2O$ (20 mg, 0.37 mol) was added to the solution. The resulting solution was stirred 95 °C for 3 h. Then, cool the reaction mixture and add 2 M HCl solution dropwise. The organic layer was washed with brine, dried over anhydrous $Na_2SO_4$ and concentrated. The residue was purified by reversed-phase HPLC using ODS-3 column and eluted with $H_2O$/acetonitrile containing 0.1% formic acid to yield **16** as a purple powder (70 mg; yield, 24%).

**$^1$H NMR (500 MHz, $CDCl_3$)** δ 8.42 (d, $J$ = 7.0 Hz, 1H), 8.16 (s, 1H), 8.14 (d, $J$ = 7.0 Hz, 1H), 7.61 (d, $J$ = 7.5 Hz, 1H), 6.94 (d, $J$ = 2.0 Hz, 1H), 6.85 (dd, $J$ = 7.0 Hz, $J$ = 2.0 Hz, 1H), 3.03 (s, 6H) ppm.

**$^{13}$C NMR (125 MHz, $CDCl_3$)** δ 164.7, 156.3, 151.8, 150.2, 149.8, 147.4, 130.2, 125.7, 121.8, 121.2, 120.7, 117.9, 110.9, 110.2, 92.9, 39.9 ppm.
**HRMS (FAB+)** (*m*/*z*): calcd $[M]^+$ for $C_{17}H_{13}N_3O_3S$ 339.0678, found 339.0661

**(7-(6-(dimethylamino)benzofuran-2-yl)benzo[*c*][1,2,5]thiadiazole-4-carbonyl)glycine (DMABT-FA2)**
Compound **16** (50 mg, 0.15 mmol) was dissolved in dry DMF (5 mL) and then HATU (70 mg, 0.18 mmol) and dry DIEA (51 µL, 0.29 mmol) were added to it. The reaction mixture stirred for 1-1.5 h at 0 °C under $N_2$ atmosphere. The compound glycine *tert*-butyl ester hydrochloride (37 mg, 0.22 mmol) dissolved in DMF (1 mL) was added to the reaction mixture and stirred 3 hours at room temperature under $N_2$ atmosphere. After removing the solvent, the residue was stirred for 1 hour in DCM/TFA (1:1) solution. After removing the solvent, the residue was purified by reversed-phase HPLC using ODS-3 column and eluted with $H_2O$/acetonitrile containing 0.1% formic acid to yield **DMABT-FA2** as a purple powder (6.4 mg, 41%).
**$^1$H NMR (500 MHz, $CDCl_3$)** δ 9.38 (t, *J* = 5.5 Hz, 1H), 8.49 (d, *J* = 7.0 Hz, 1H), 8.20 (d, *J* = 7.0 Hz, 1H), 8.16 (s, 1H), 7.61 (d, *J* = 7.5 Hz, 1H), 6.95 (d, *J*= 2.0 Hz, 1H), 6.85 (dd, *J* = 7.0 Hz, *J* = 2.0 Hz, 1H), 4.21 (d, *J* = 5.5 Hz, 2H) 3.03 (s, 6H) ppm.
**$^{13}$C NMR (125 MHz, $CDCl_3$)** δ 171.6, 171.3, 163.3, 157.2, 152.4, 148.6, 133.5, 131.3, 131.3, 126.2, 122.9, 119.2, 111.6, 94.2, 80.2, 42.3, 41.0 ppm.
**HRMS (FAB+)** (*m*/*z*): calcd $[M]^+$ for $C_{19}H_{16}N_4O_4S$ 396.0892, found 396.0898

**6-(7-(6-(dimethylamino)benzofuran-2-yl)benzo[*c*][1,2,5]thiadiazole-4-carboxamido)hexanoic acid (DMABT-FA6)**
Compound **16** (21 mg, 0.061 mmol) was dissolved in dry DMF (3 mL) and then HATU (23 mg, 0.066 mmol) and dry DIEA (51 µL, 0.29 mmol) were added to it. The reaction mixture stirred for 1-1.5 h at 0 °C under $N_2$ atmosphere. The compound 6 amino hexanoic acid (8 mg, 0.06 mmol) dissolved in DMF (1 mL) was added to the reaction mixture and stirred 3 hours at room temperature under $N_2$ atmosphere. After removing the solvent, the residue was purified by reversed-phase HPLC using ODS-3 column and eluted with $H_2O$/acetonitrile containing 0.1% formic acid to yield **DMABT-FA6** as a purple powder (10 mg, 43%).
**$^1$H NMR (500 MHz, $CDCl_3$)** δ 9.38 (t, *J* = 5.5 Hz, 1H), 8.35 (d, *J* = 7.0 Hz, 1H), 8.11 (d, *J* = 7.0 Hz, 1H), 8.05 (s, 1H), 7.53 (d, *J* = 7.5 Hz, 1H), 6.87 (d, *J* = 2.0 Hz, 1H), 6.78 (dd, *J* = 7.0 Hz, *J* = 2.0 Hz, 1H), 3.38 (m, *J* = 5.5 Hz, 2H), 2.95 (s, 6H) , 2.17 (t, *J* = 7.5 Hz, 2H) , 1.57-1.48 (m, 4H) , 1.33 (m, 2H) ppm.
**$^{13}$C NMR (125 MHz, $CDCl_3$)** δ 174.9, 163.2, 157.2, 152.5, 150.9, 150.8, 148.5, 132.7, 125.6, 123.9, 122.8, 122.7, 119.0, 111.3, 111.2, 94.0, 41.0, 40.9, 34.1, 29.4, 26.5, 24.7 ppm.
**HRMS (ESI+)** (*m*/*z*): calcd $[M+Na]^+$ for $C_{23}H_{24}N_4NaO_4S$ 475.1410, found 475.1395.

#### 1.4. Synthesis of benzocoumarin-and benzofuran–benzothiadiazole-based trialkylamine probes

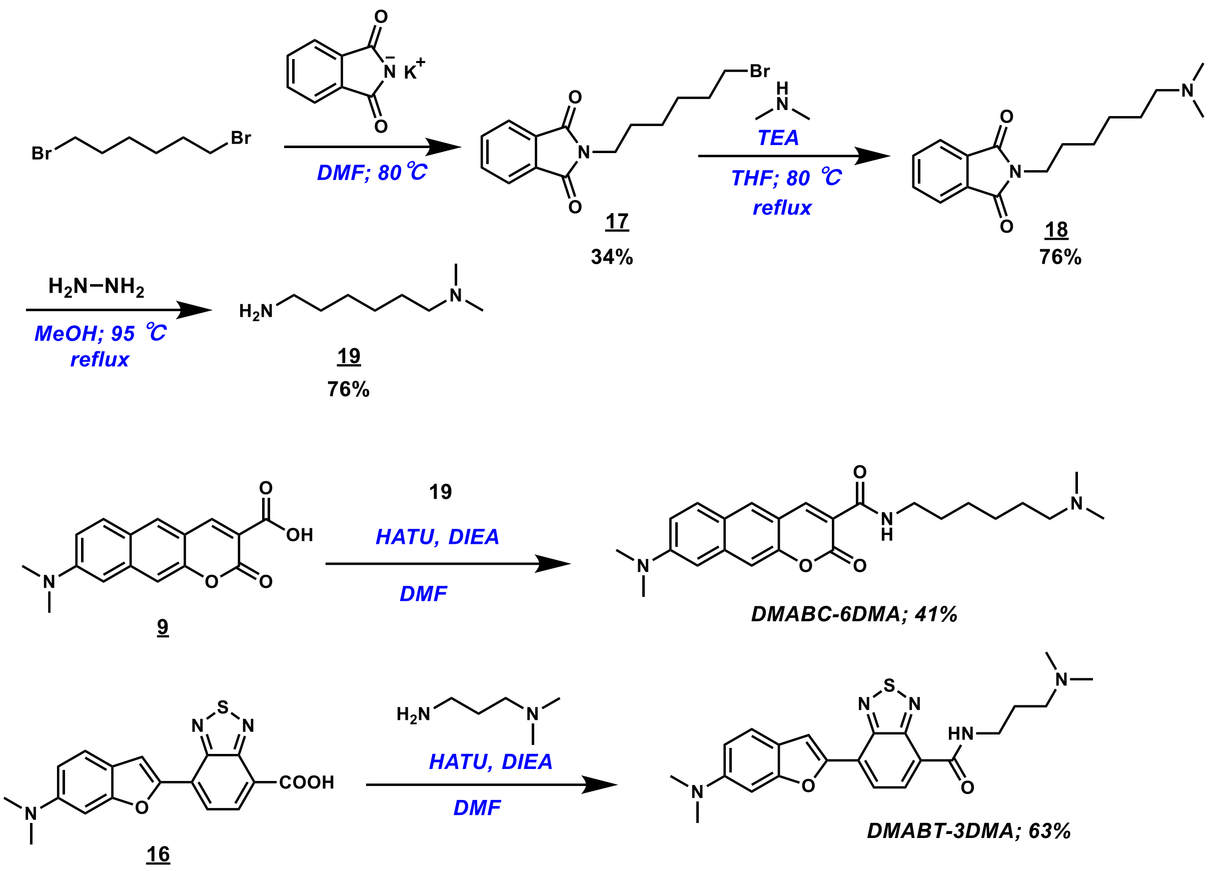


**2-(6-bromohexyl)isoindoline-1,3-dione (17)**

To a suspension of phthalimide potassium salt (780 mg, 4.1 mmol) was added in the mixture of dry DMF (5 mL) and 1,6-dibromohexane (2.0 g, 8.2 mmol). The resulting mixture was stirred at 80 °C for overnight. After being cooled to room temperature, water (50 mL) and EtOAc (50 mL) were added to the mixture. The two layers were separated, and the aqueous layer was extracted with EtOAc (100 mL). The combined organic extracts were washed with brine, dried over anhydrous $Na_2SO_4$ and concentrated. The residue was purified by silica gel column chromatography (eluent: 15% EtOAc in hexane) to afford compound **17** as a pale yellow oil (0.88 g, 34%).

**$^1$H NMR (500 MHz, CDCl$_3$)** δ 7.84 (dd, *J* = 6.5 Hz, *J* = 3.0 Hz, 2H), 7.71 (dd, *J* = 6.5 Hz, *J* = 3.0 Hz, 2H), 3.69 (t, *J* = 7.0 Hz, 2H), 3.40 (t, *J* = 7.0 Hz, 2H), 1.86 (m, 2H), 1.70 (m, 2H), 1.49 (m, 2H), 1.38 (m, 2H) ppm.

**$^{13}$C NMR (125MHz, CDCl$_3$)** δ 168.4, 133.8, 132.1, 123.1, 37.8, 33.7, 32.6, 28.4, 27.7, 26.0 ppm.

**HRMS (ESI+)** (*m*/*z*): calcd $[M+Na]^+$ for $C_{14}H_{16}BrNNaO_2$ 332.0257, found 332.0252.

**2-(6-(dimethylamino)hexyl)isoindoline-1,3-dione (18)**

To a solution of compound **17** (400 mg, 1.3 mmol) was added in the mixture of 2 M dimethylamine in THF solution (1.3 mL) and TEA (0.4 mL). The resulting mixture was stirred at 80 °C for overnight. After being cooled to room temperature,

THF was removed under reduced pressure, and then EtOAc (50 mL) was added to the residue. The organic layer was washed with sat. $NaHCO_3$, brine, dried over anhydrous $Na_2SO_4$ and concentrated to afford compound **18** as a white solid (0.27 g, 76%).

**$^1$H NMR (500 MHz, CDCl$_3$)** δ 7.84 (dd, *J* = 6.0 Hz, *J* = 3.0 Hz, 2H), 7.71 (dd, *J* = 6.0 Hz, *J* = 3.0 Hz, 2H), 3.68 (t, *J* = 7.0 Hz, 2H), 2.24 (t, *J* = 7.0 Hz, 2H), 2.20 (s, 6H), 1.68 (m, 2H), 1.46 (m, 2H), 1.35 (m, 4H) ppm.

**$^{13}$C NMR (125 MHz, CDCl$_3$)** δ 168.5, 133.9, 132.2, 123.2, 59.7, 45.5, 38.0, 28.6, 27.6, 27.1, 26.8 ppm.

**HRMS (ESI+)** (*m/z*): calcd $[M+H]^+$ for $C_{16}H_{23}N_2O_2$ 275.1755, found 275.1764.

**$N^1$,$N^1$-dimethylhexane-1,6-diamine (19)**

To a solution of compound **18** (270 mg, 1.0 mmol) was added in the mixture of MeOH (6 mL) and $NH_2NH_2$ (1 mL). The resulting mixture was stirred in refluxing for 4 hours. After being cooled to room temperature, the mixture was filtered. The filtrate was concentrated and then added DCM (10 mL). The solution was filtered and the filtrate was concentrated. Repeat this procedure three times to afford compound **19** as a pale yellow oil (0.11 g, 76%).

**$^1$H NMR (500 MHz, CDCl$_3$)** δ 2.69 (t, *J* = 7.0 Hz, 2H), 2.26 (t, *J* = 7.0 Hz, 2H), 2.22 (s, 6H), 1.89 (bs, 2H), 1.46 (m, 4H), 1.35 (m, 4H) ppm.

**$^{13}$C NMR (125 MHz, CDCl$_3$)** δ 59.8, 45.4 42.1, 33.7, 27.7, 27.3, 26.8 ppm.

**HRMS (ESI+)** (*m/z*): calcd $[M+H]^+$ for $C_8H_{21}N_2$ 145.1700, found 145.1709.

**8-(dimethylamino)-*N*-(6-(dimethylamino)hexyl)-2-oxo-2H-benzo[*g*]chromene-3-carboxamide (DMABC-6DMA)**

The compound **9** (10 mg, 0.035 mmol) was dissolved in dry DMF (2 mL) and then HATU (16 mg, 0.042 mmol) and dry DIEA (13 μL, 0.071 mmol) were added to it. The reaction mixture stirred for 1-1.5 h at 0 °C under $N_2$ atmosphere. The compound **19** (8.0 mg, 0.05 mmol) dissolved in DMF (1 mL) was added to the reaction mixture and stirred 3 hours at room temperature under $N_2$ atmosphere. After removing the solvent, the residue was purified by reversed-phase HPLC using ODS-3 column and eluted with $H_2O$/acetonitrile containing 0.1% formic acid to yield **DMABC-6DMA** as an orange powder (6.4 mg, 41%).

**$^1$H NMR (500 MHz, CDCl$_3$)** δ 8.84 (s, 1H), 8.83 (s, 1H), 7.98 (s, 1H), 7.78 (d, *J* = 8.0 Hz, 1H), 7.43 (s, 1H), 7.15 (dd, *J* = 8.0 Hz, *J* = 2.5 Hz, 1H), 6.79 (d, *J* = 2.5 Hz, 1H), 3.46 (q, *J* = 7.0 Hz, 2H), 3.15 (s, 6H), 2.29 (t, *J* = 7.0 Hz, 2H), 2.24 (s, 6H), 1.64 (m, 2H), 1.52-1.33 (m, 6H) ppm.

**$^{13}$C NMR (125 MHz, CDCl$_3$)** δ 162.4, 162.2, 151.3, 150.6, 148.7, 138.3, 130.9, 130.4, 123.8, 116.2, 115.4, 114.8, 109.5, 103.8, 59.7, 45.3, 40.3, 39.8, 29.5, 27.4, 27.2, 27.0 ppm.

**HRMS (ESI+)** (*m/z*): calcd $[M+H]^+$ for $C_{24}H_{32}N_3O_3$ 410.2439, found 410.2449.

**7-(6-(dimethylamino)benzofuran-2-yl)-*N*-(3-(dimethylamino)propyl)-*N*-methylbenzo[*c*][1,2,5]thiadiazole-4-carboxamide (DMABT-3DMA)**

Compound **16** (20 mg, 0.059 mmol) was dissolved in dry DMF (5 mL) and then HATU (26 mg, 0.070 mmol) and dry DIEA (20 μL, 0.12 mmol) were added to it. The reaction mixture stirred for 1-1.5 h at 0 °C under $N_2$ atmosphere. The compound *N,N*-dimethyl-1,3-diaminopropane (9.0 mg, 0.09 mmol) dissolved in DMF (1 mL) was added to the reaction mixture and stirred 3 hours at room temperature under $N_2$ atmosphere. After removing the solvent, the residue was purified by reversed-phase HPLC using ODS-3 column and eluted with $H_2O$/acetonitrile containing 0.1% formic acid to yield **DMABT-3DMA** as a purple powder (15 mg, 63%).

**$^1$H NMR (500 MHz, DMSO-*d*$_6$)** δ 9.13 (t, *J* = 6.0 Hz, 1H), 8.43 (d, *J* = 7.0 Hz, 1H), 8.19 (d, *J* = 7.0 Hz, 1H), 8.13 (s, 1H), 7.61 (d, *J* = 7.5 Hz, 1H), 6.93 (d, *J* = 2.0 Hz, 1H), 6.85 (dd, *J* = 7.0 Hz, *J* = 2.0 Hz, 1H), 3.54 (m, *J* = 5.5 Hz, 2H), 3.16 (t, *J* = 8.0 Hz, 2H), 3.03 (s, 6H), 2.81 (s, 6H), 1.99 (m, 2H) ppm.

**$^{13}$C NMR (125 MHz, DMSO-*d*$_6$)** δ 164.1, 157.4, 152.7, 151.3, 150.5, 148.7, 133.7, 126.9, 122.6, 122.2, 121.9, 119.5, 111.6, 110.6, 94.0, 56.8, 44.7, 41.0, 37.9, 26.8 ppm.

**HRMS (FAB+)** (*m*/*z*): calcd $[M+H]^+$ for $C_{22}H_{26}N_5O_2S$ 424.1802, found 424.1789

#### 2. NMR spectra

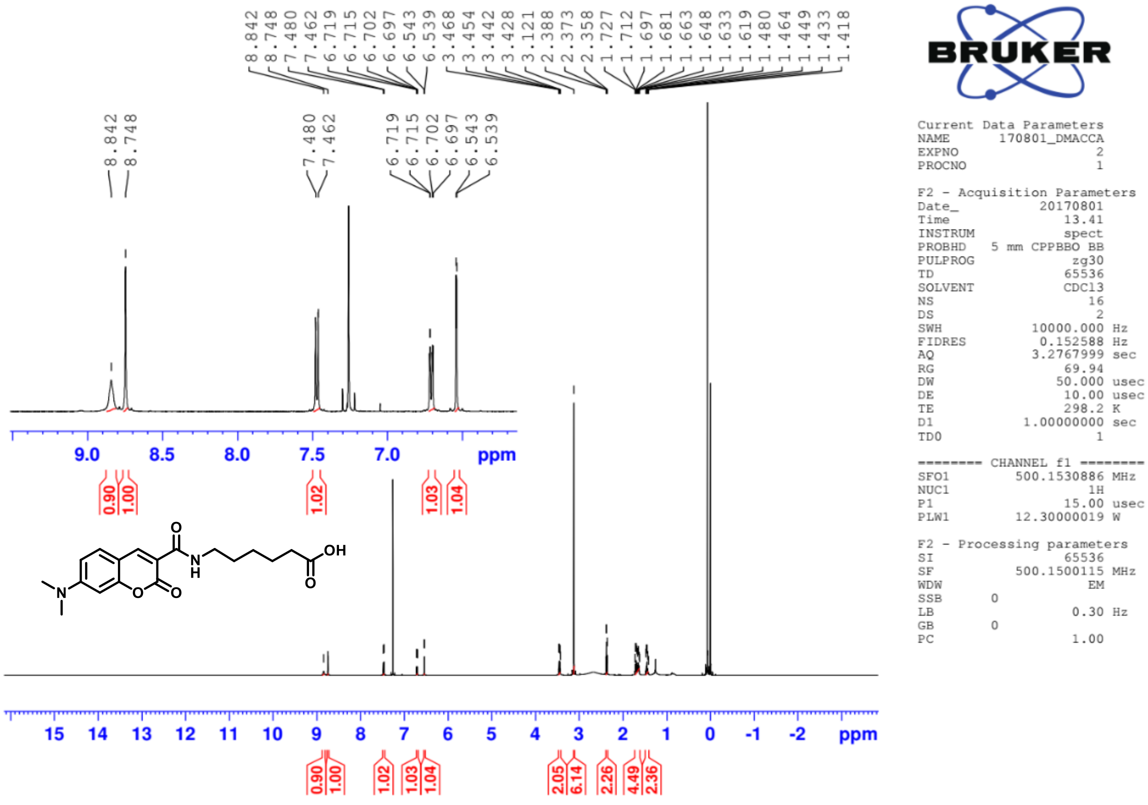


[1]H NMR spectrum of compound **DMAC-FA6** in 500 MHz ($CDCl_3$)

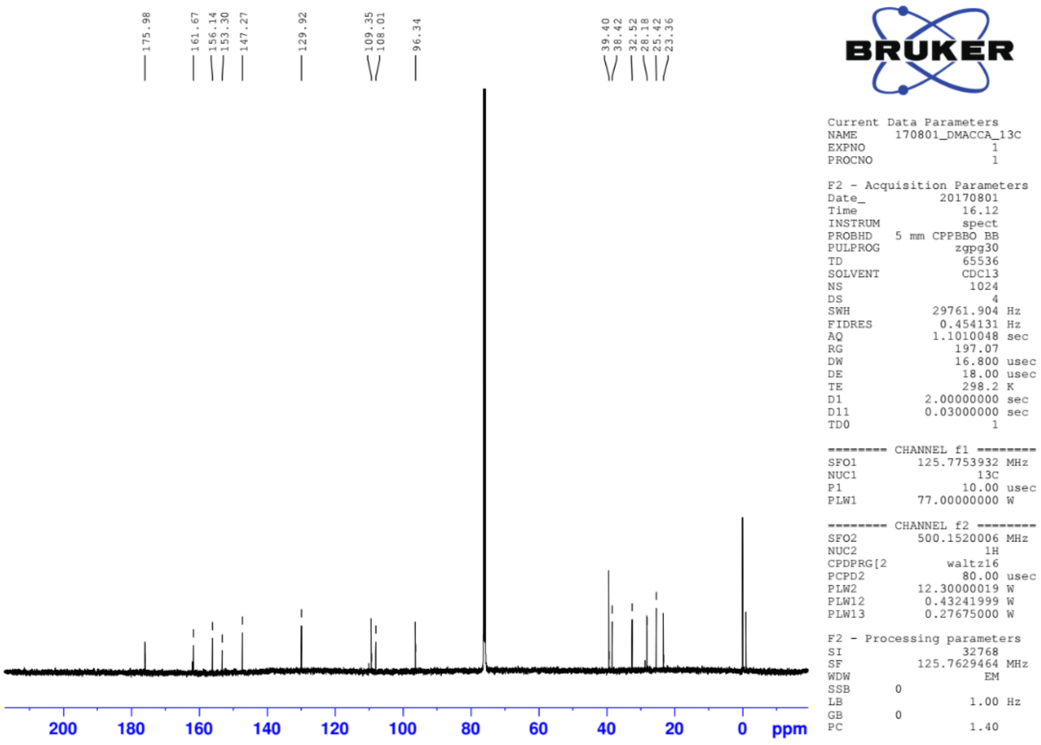


[13]C NMR spectrum of compound **DMAC-FA6** in 125 MHz ($CDCl_3$)

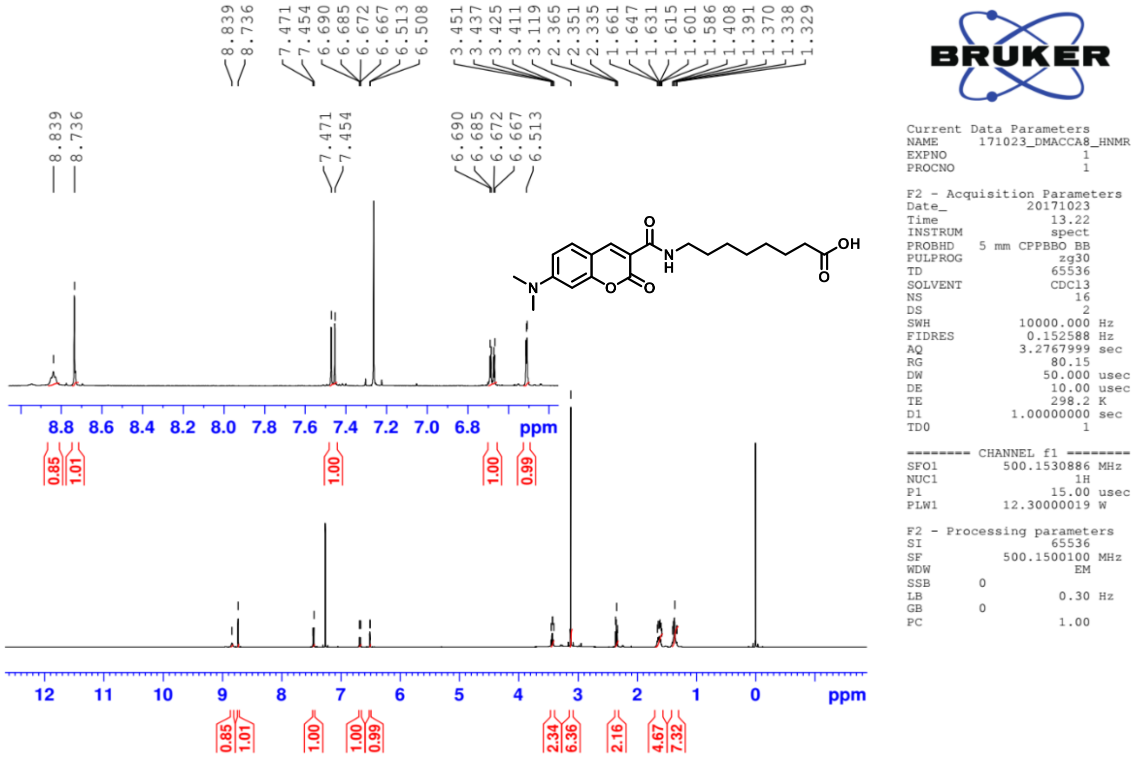


$^{1}H$ NMR spectrum of compound **DMAC-FA8** in 500 MHz ($CDCl_3$)

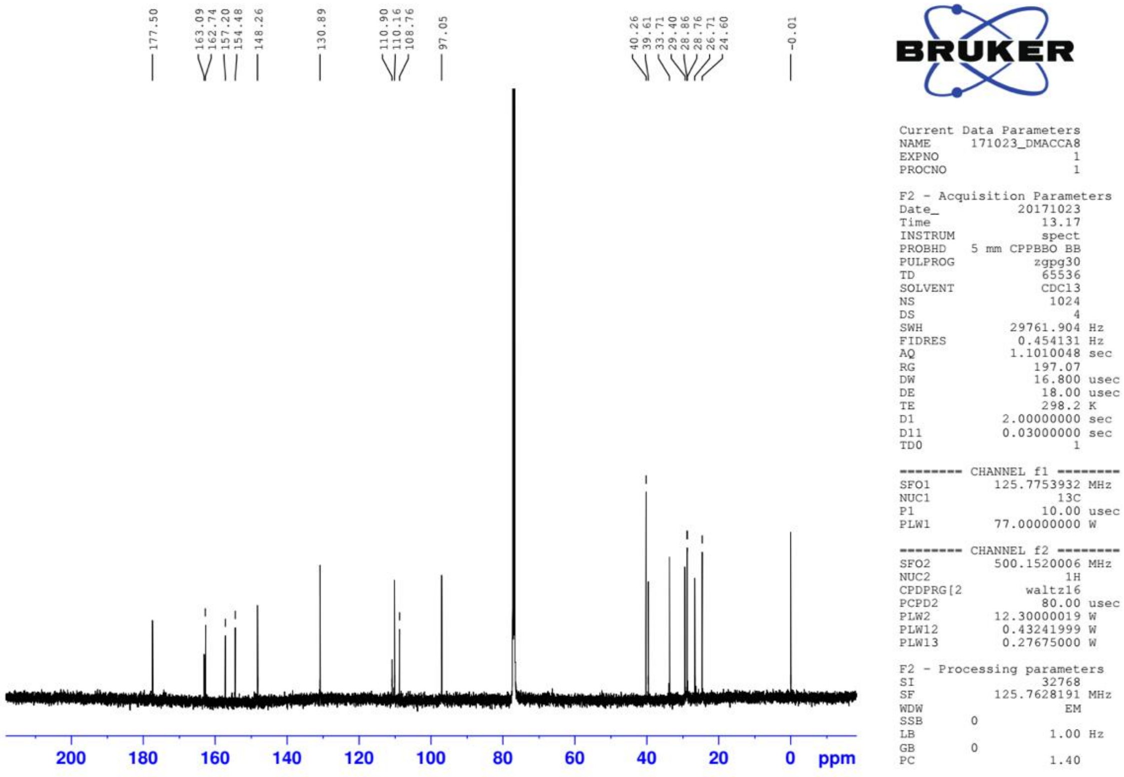


$^{13}C$ NMR spectrum of compound **DMAC-FA8** in 125 MHz ($CDCl_3$)

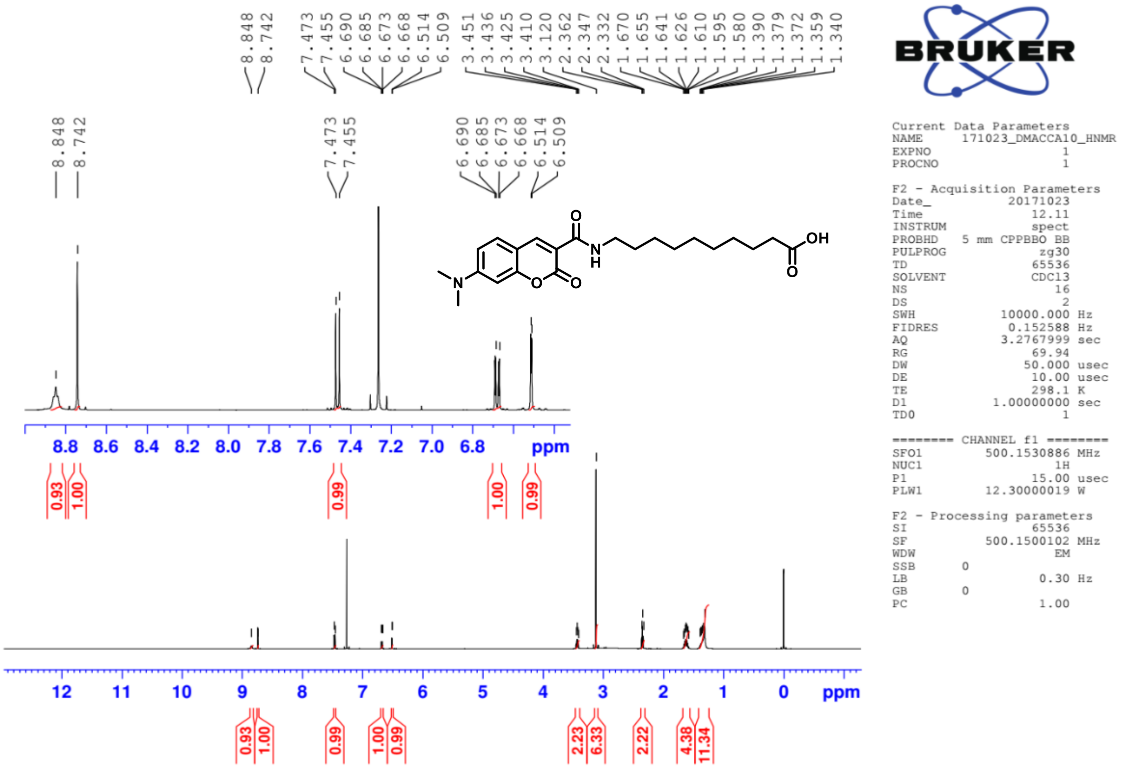


[1]H NMR spectrum of compound **DMAC-FA10** in 500 MHz ($CDCl_3$)

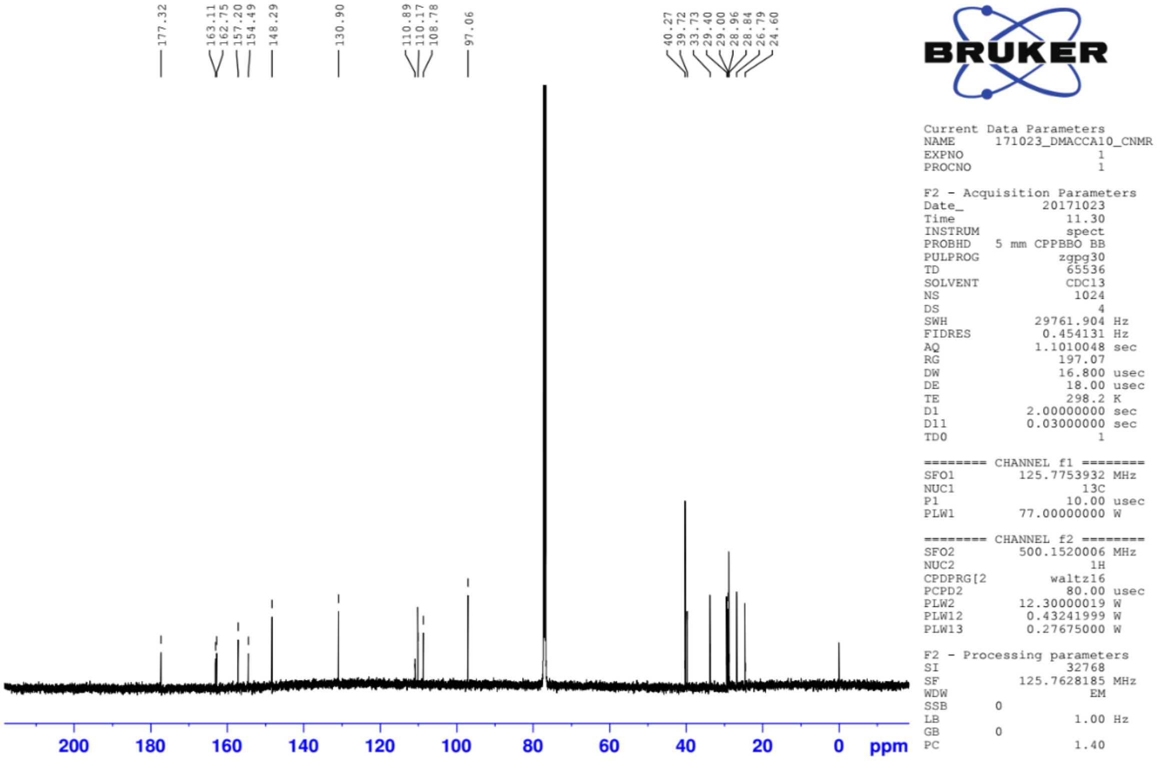


[13]C NMR spectrum of compound **DMAC-FA10** in 125 MHz ($CDCl_3$)

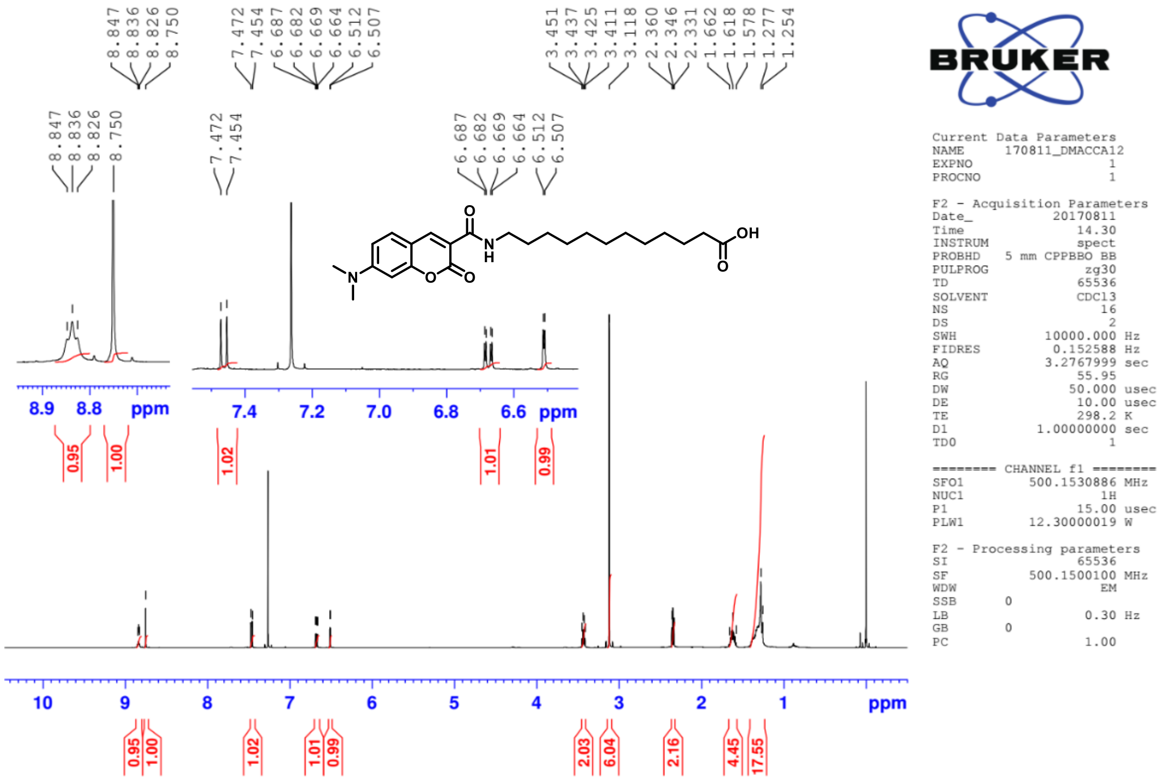


$^{1}$H NMR spectrum of compound **DMAC-FA12** in 500 MHz ($CDCl_3$)

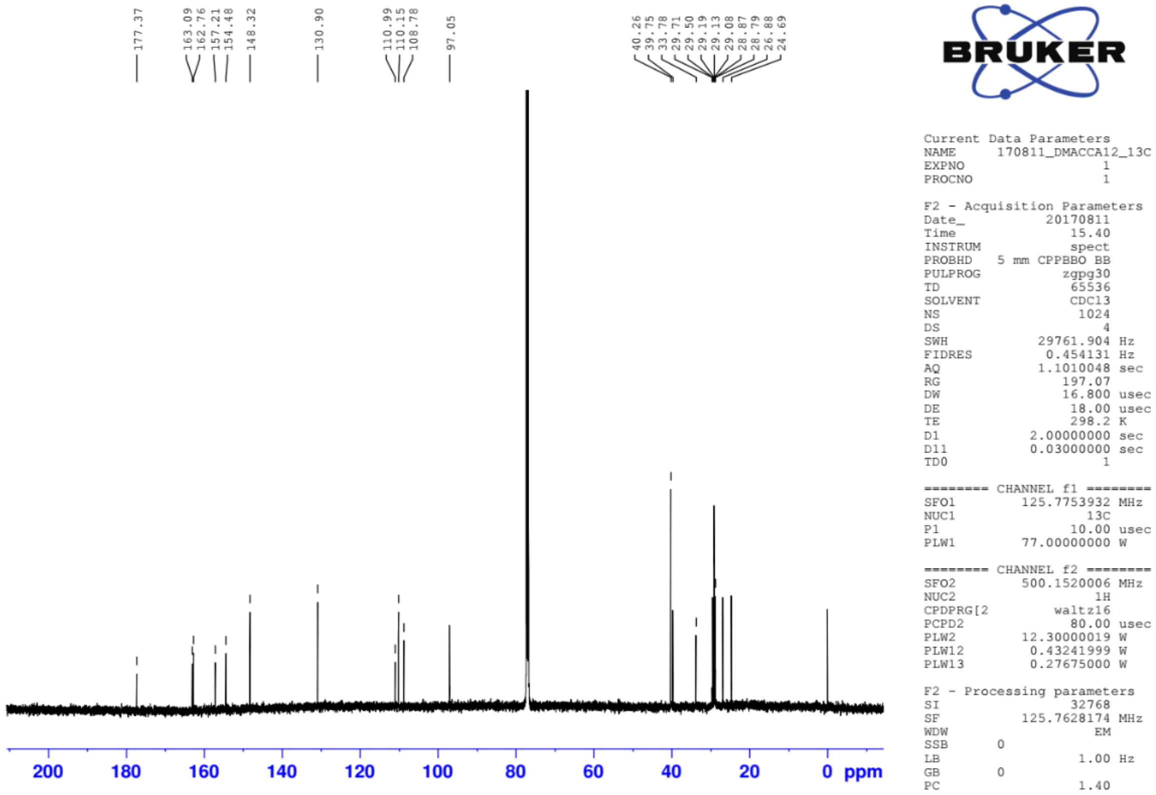


$^{13}$C NMR spectrum of compound **DMAC-FA12** in 125 MHz ($CDCl_3$)

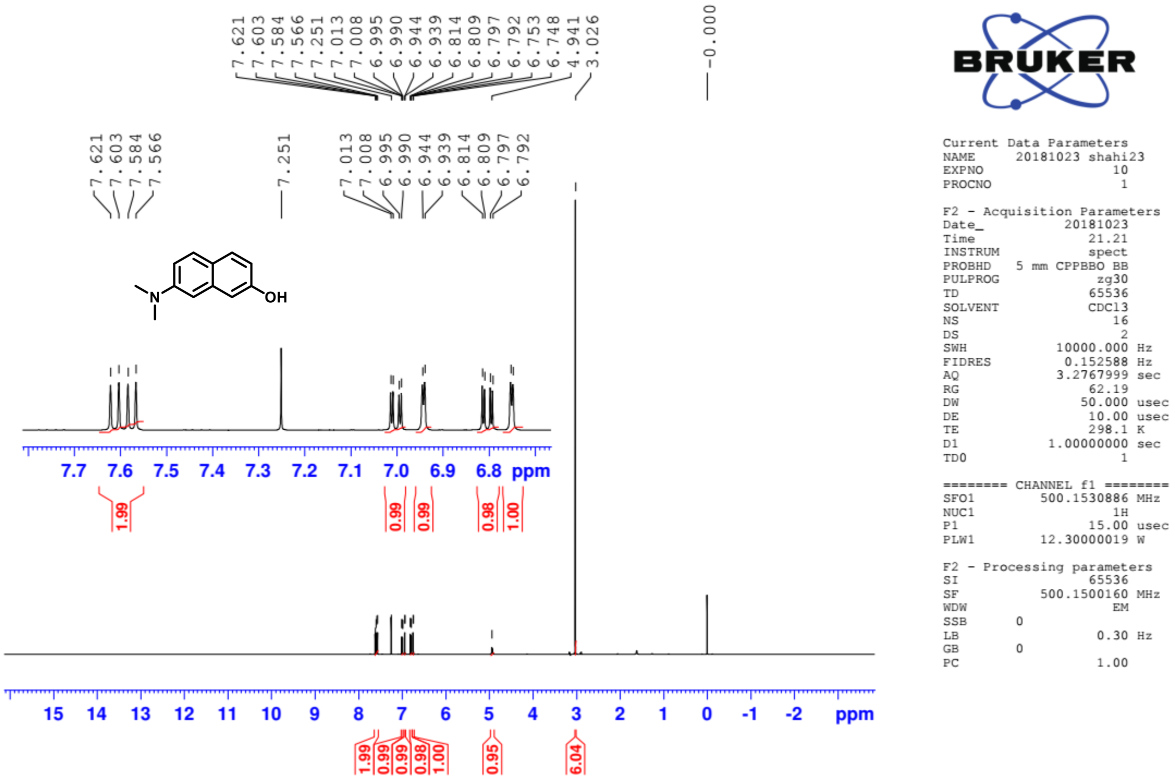


[1]H NMR spectrum of compound **4** in 500 MHz ($CDCl_3$)

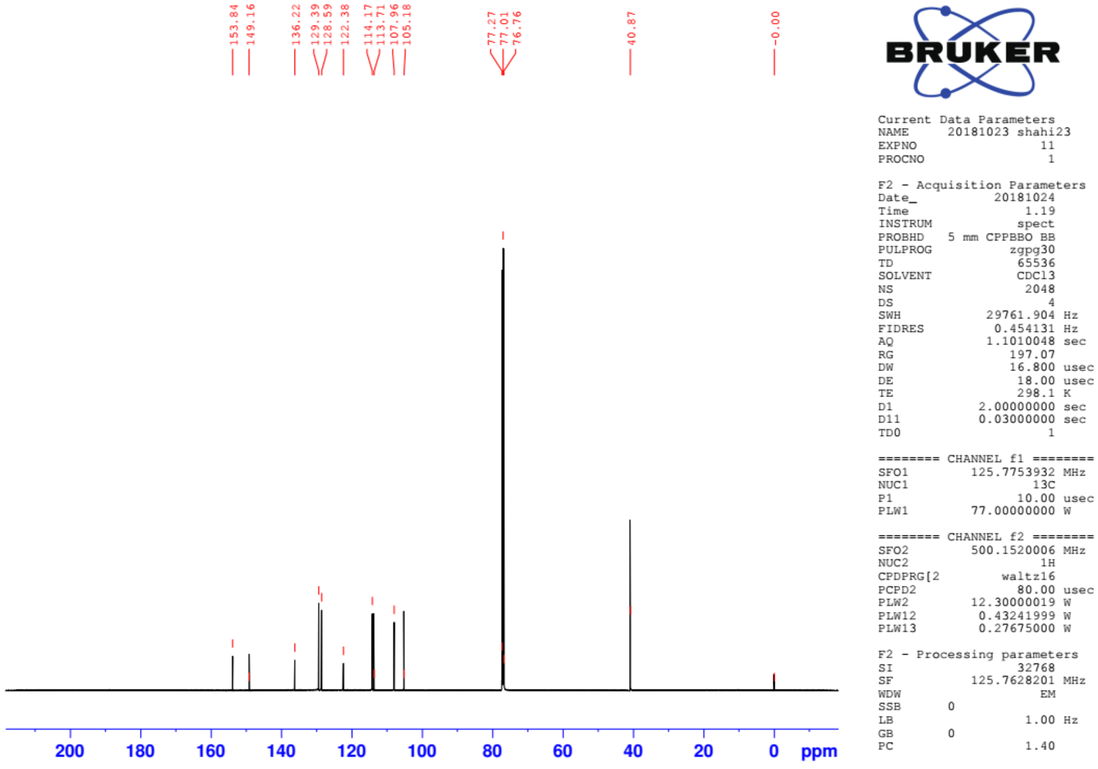


[13]C NMR spectrum of compound **4** in 125 MHz ($CDCl_3$)

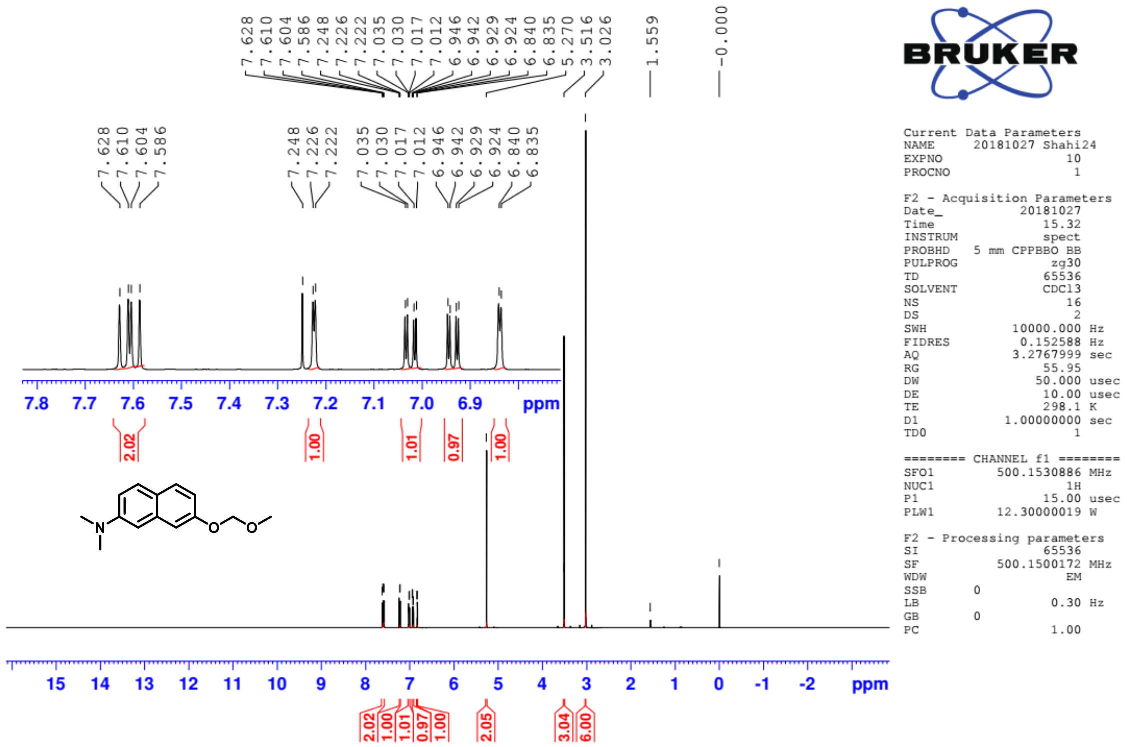

[1]H NMR spectrum of compound **5** in 500 MHz ($CDCl_3$)

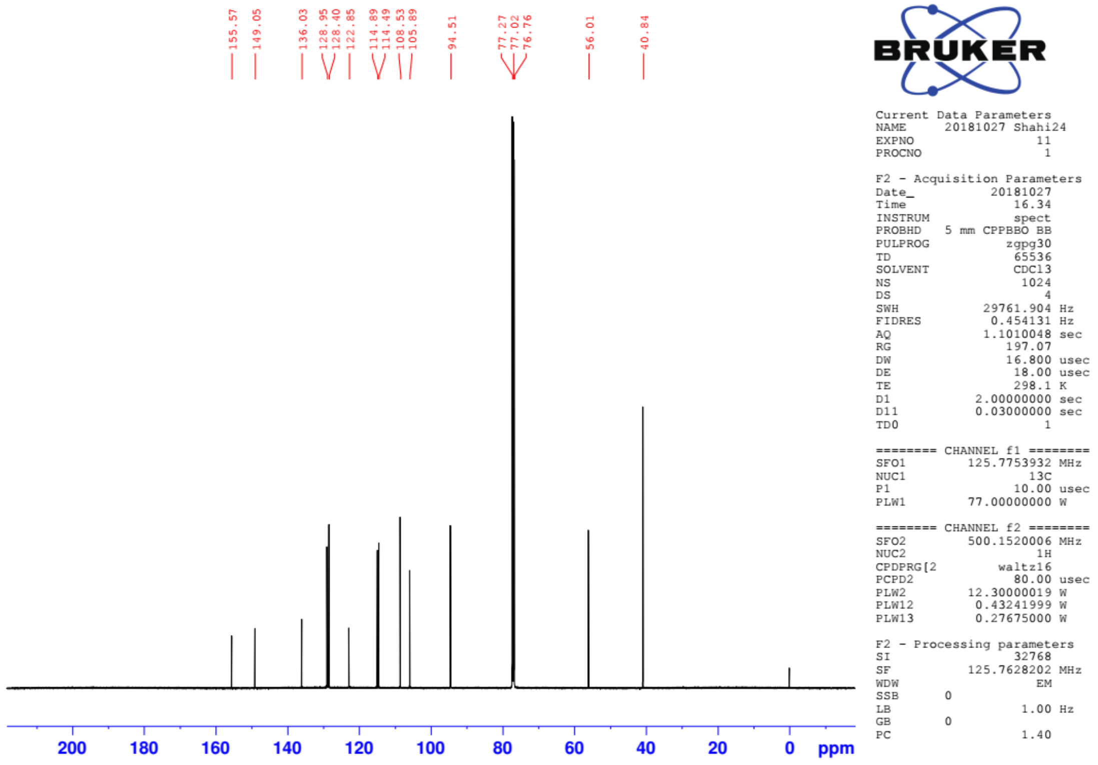

[13]C NMR spectrum of compound **5** in 125 MHz ($CDCl_3$)

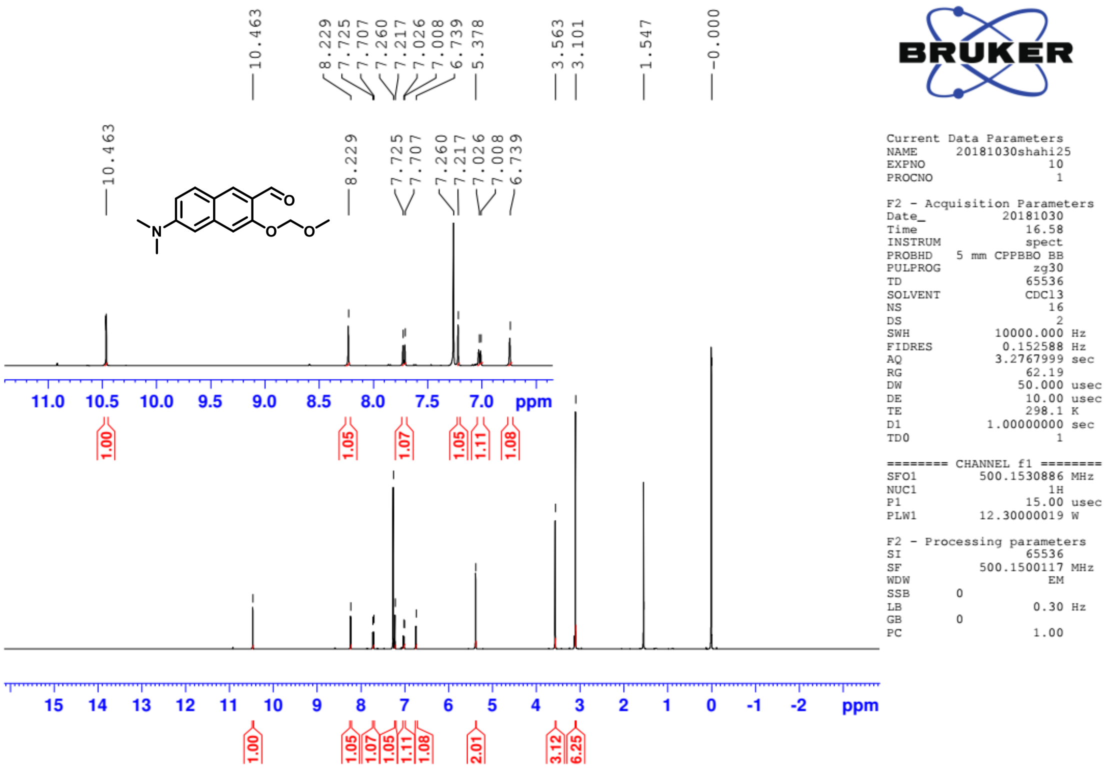

[1]H NMR spectrum of compound **6** in 500 MHz ($CDCl_3$)

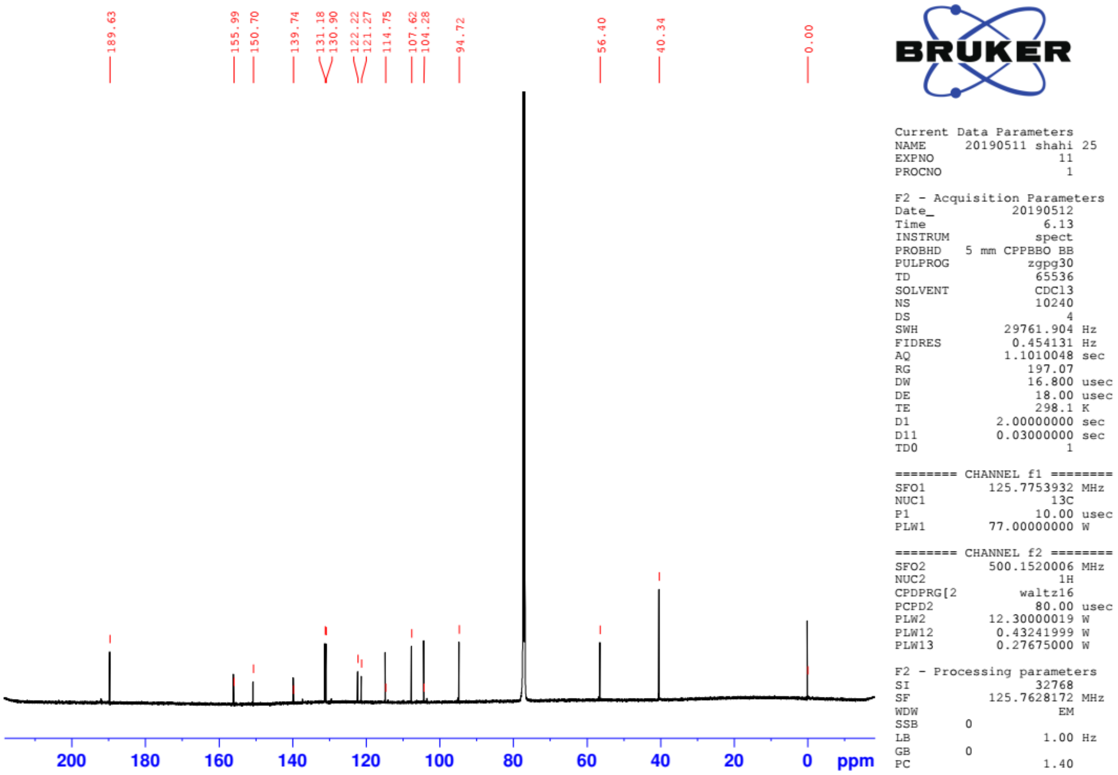

[13]C NMR spectrum of compound **6** in 125 MHz ($CDCl_3$)

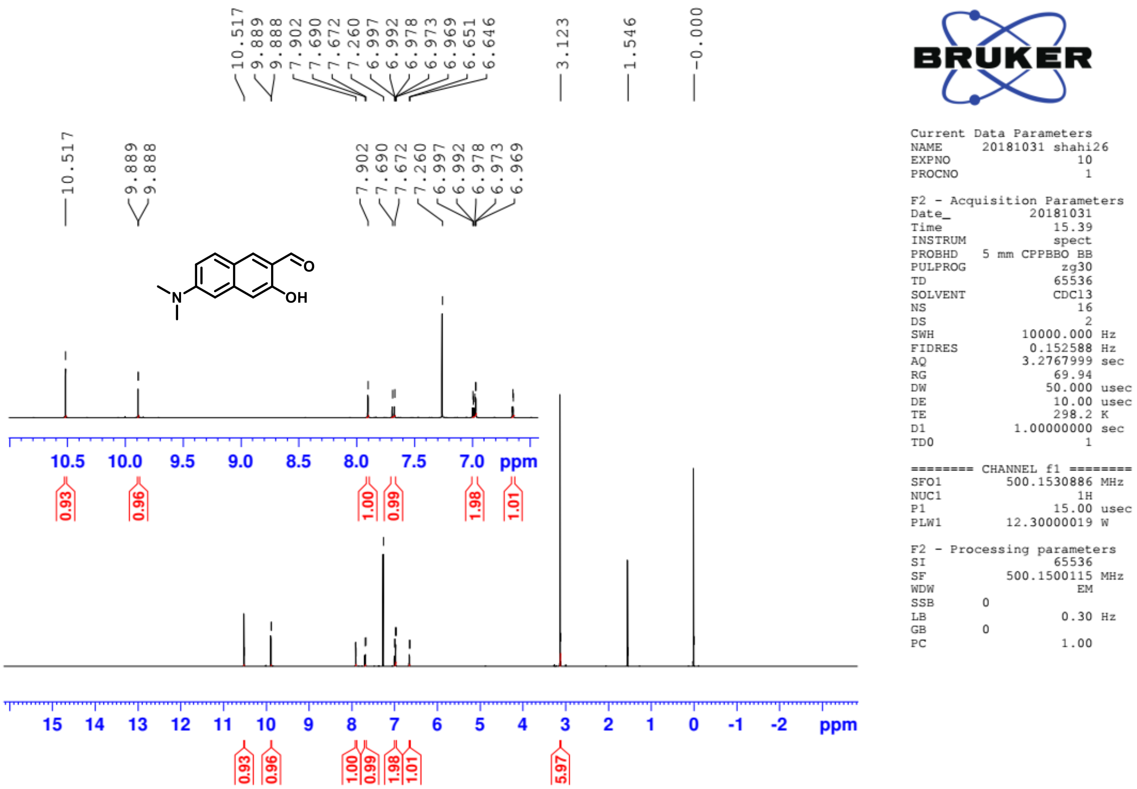


$^{1}H$ NMR spectrum of compound **7** in 500 MHz ($CDCl_3$)

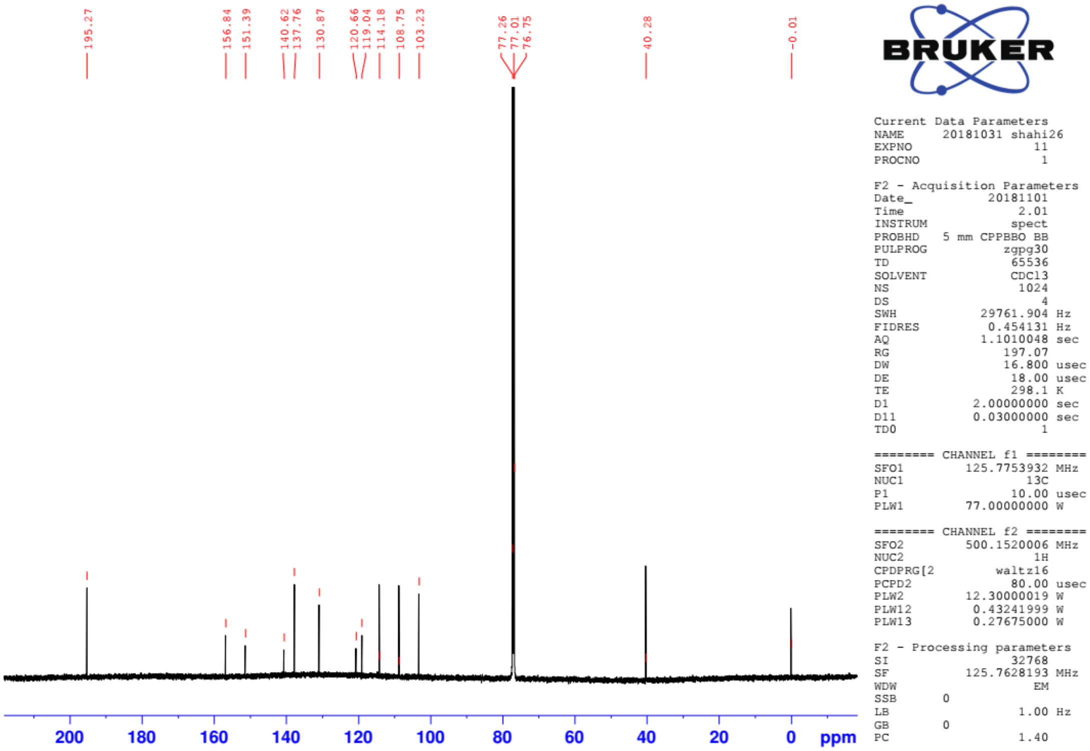


$^{13}C$ NMR spectrum of compound **7** in 125 MHz ($CDCl_3$)

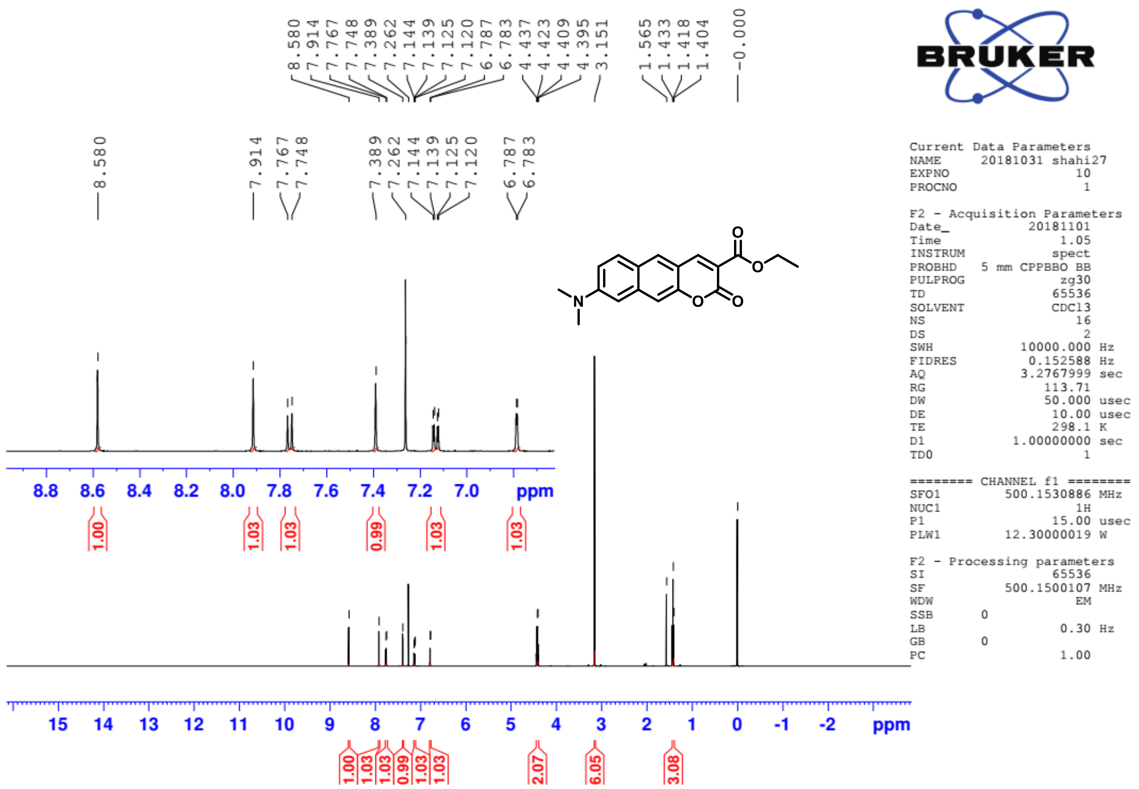


$^1$H NMR spectrum of compound **8** in 500 MHz ($CDCl_3$)

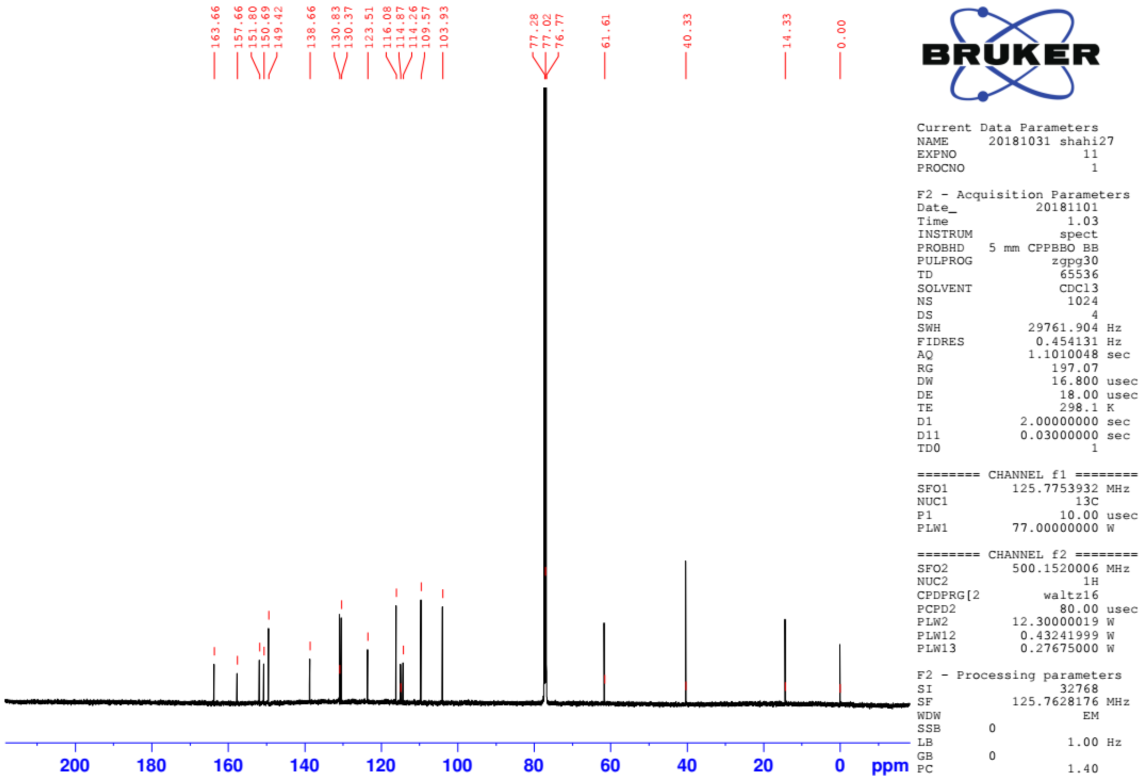


$^{13}$C NMR spectrum of compound **8** in 125 MHz ($CDCl_3$)

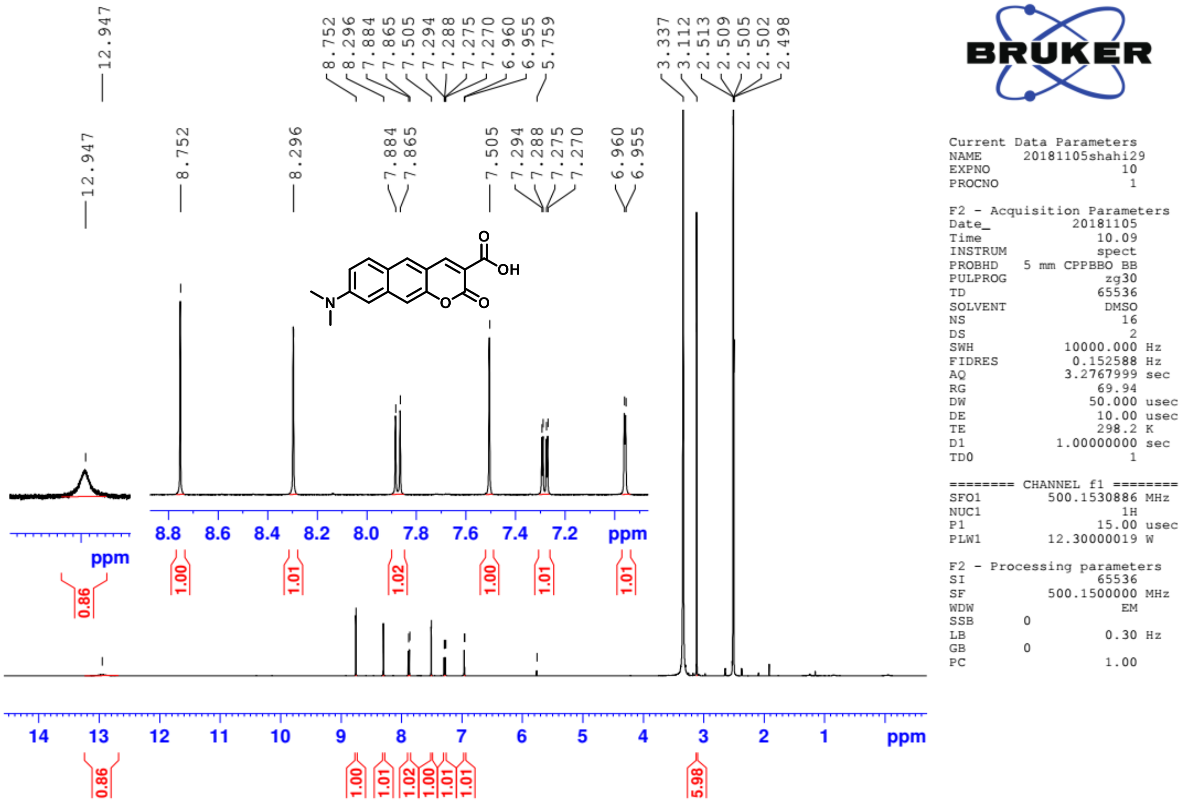

$^{1}$H NMR spectrum of compound **9** in 500 MHz (DMSO-$d_6$)

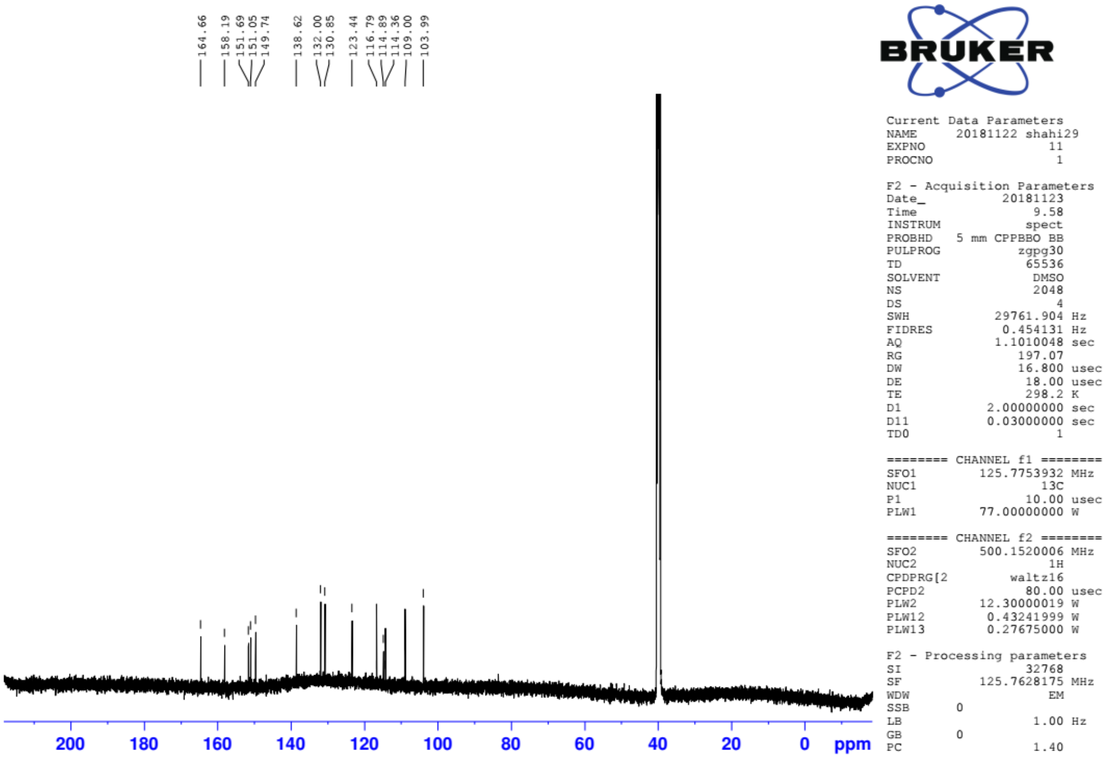

$^{13}$C NMR spectrum of compound **9** in 125 MHz (DMSO-$d_6$)

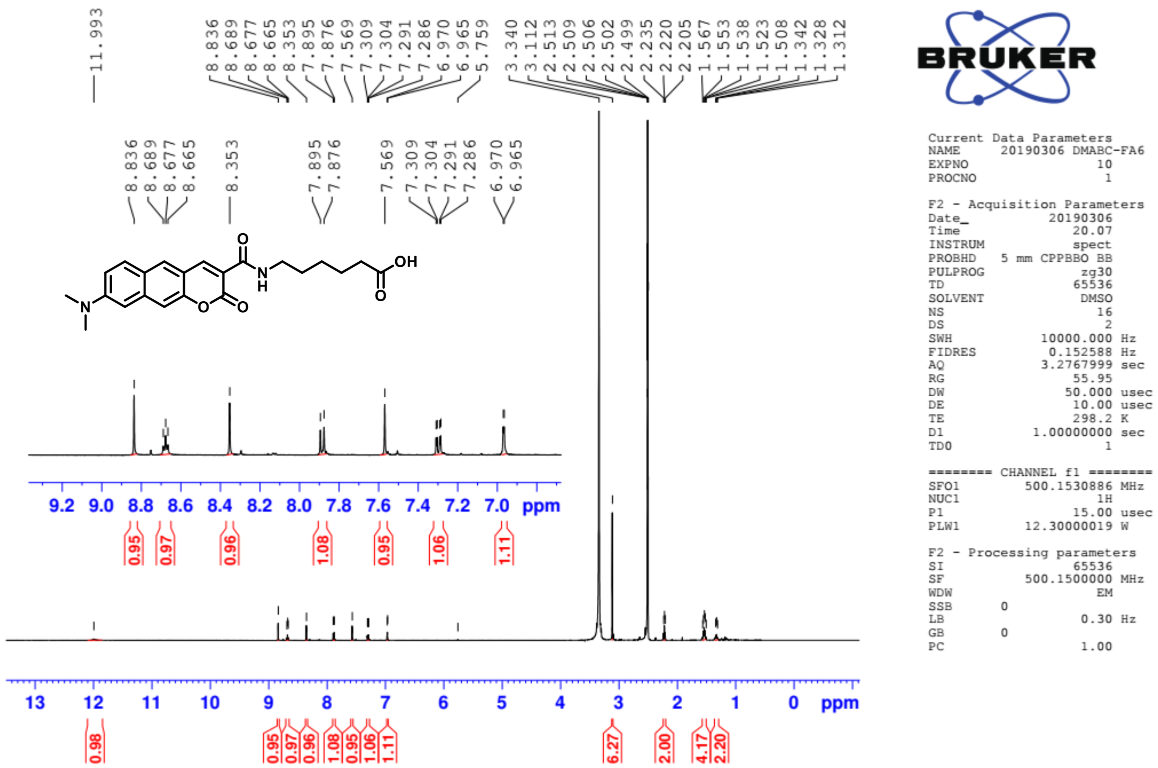


[1]H NMR spectrum of compound **DMABC-FA6** in 500 MHz (DMSO-$d_6$)

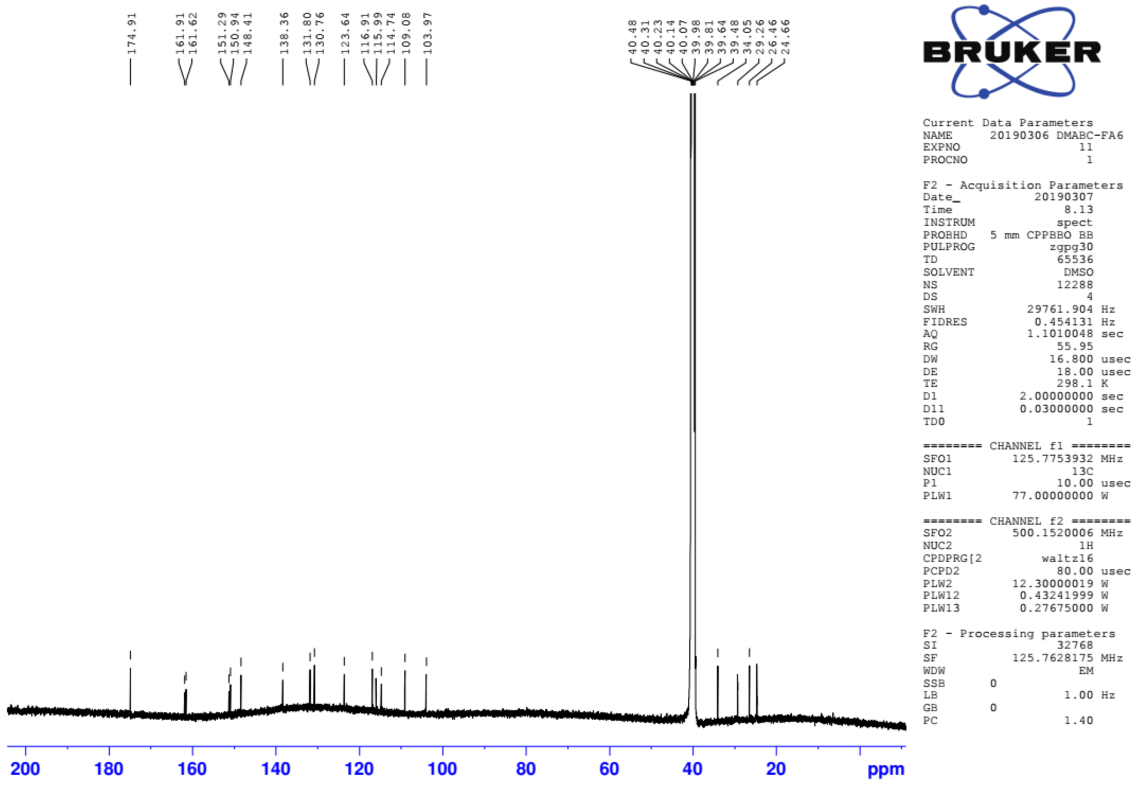


[13]C NMR spectrum of compound **DMABC-FA6** in 125 MHz (DMSO-$d_6$)

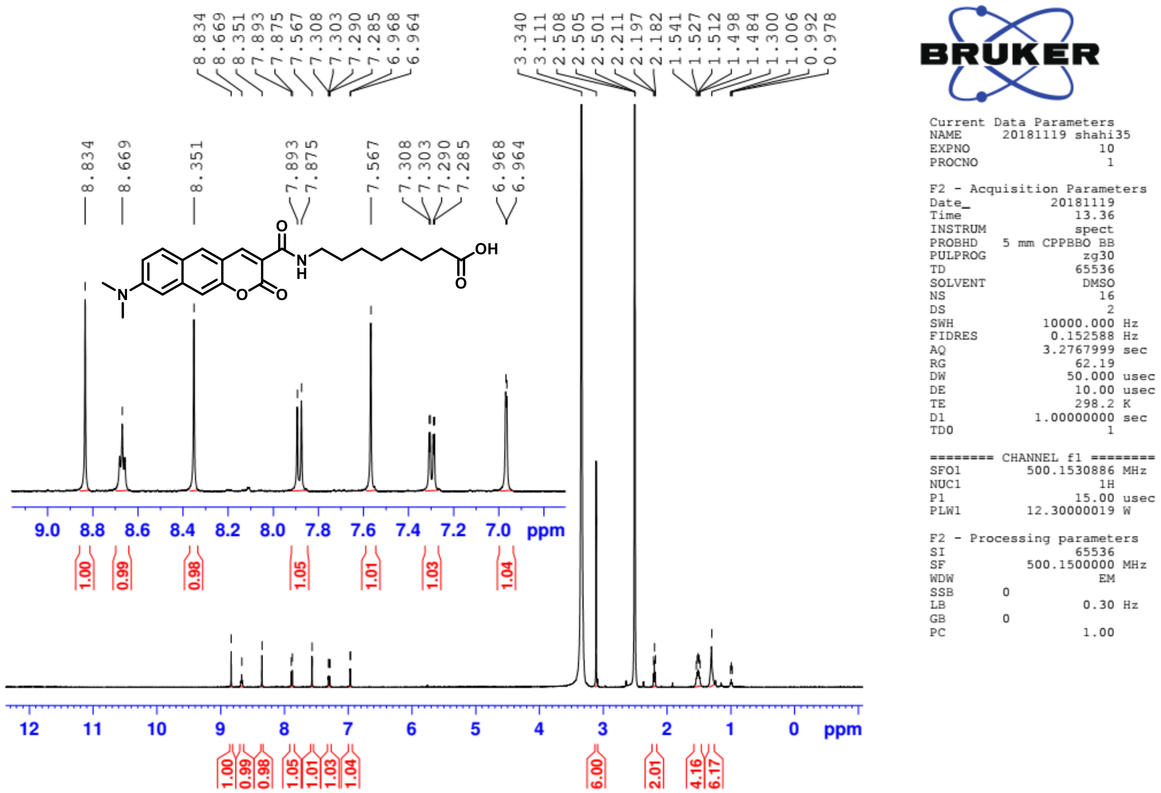

[1]H NMR spectrum of **DMABC-FA8** in 500 MHz (DMSO-$d_6$)

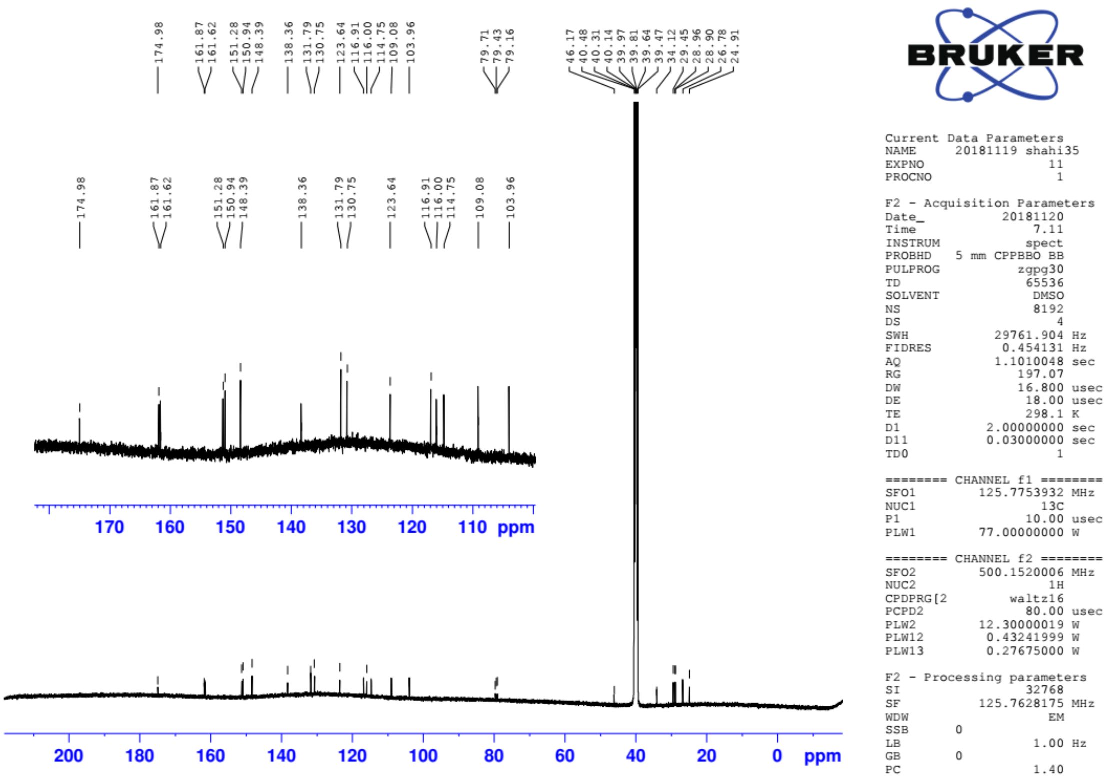

[13]C NMR spectrum of **DMABC-FA8** in 125 MHz (DMSO-$d_6$)

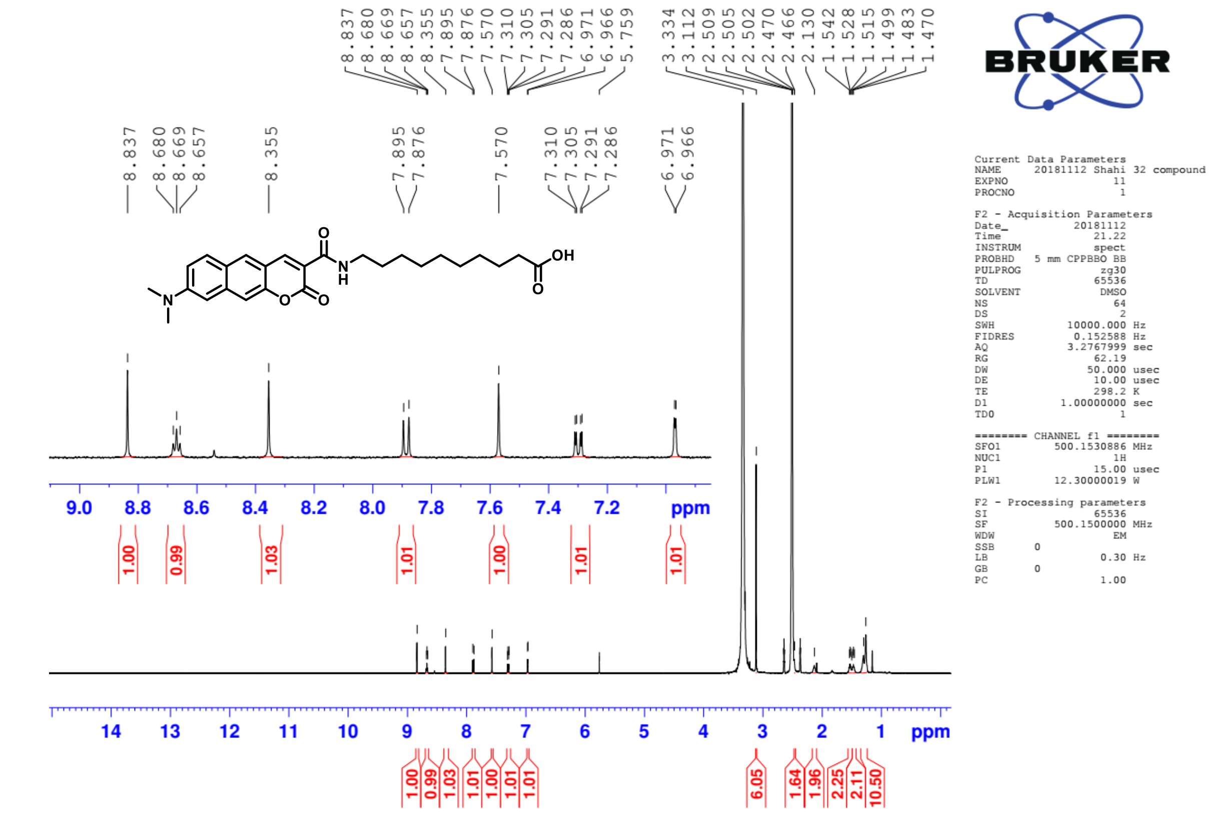


$^1$H NMR spectrum of **DMABC-FA10** in 500 MHz (DMSO-$d_6$)

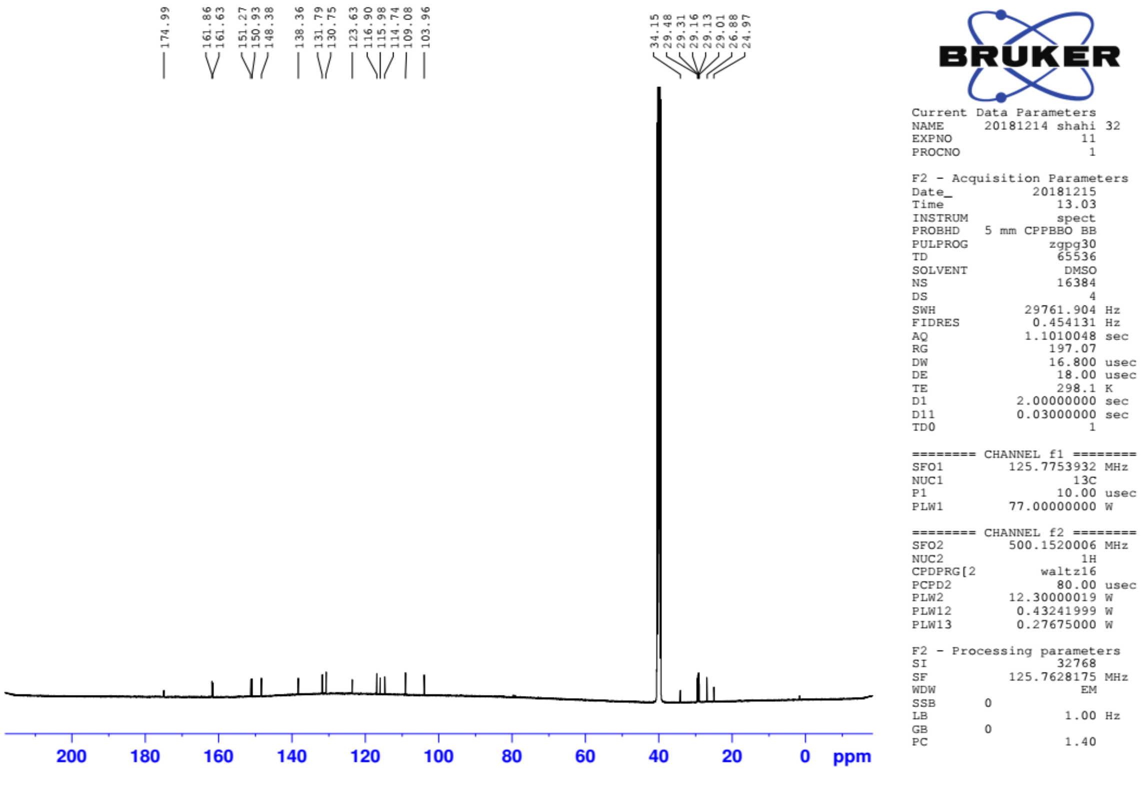


$^1$H NMR spectrum of **DMABC-FA10** in 125 MHz (DMSO-$d_6$)

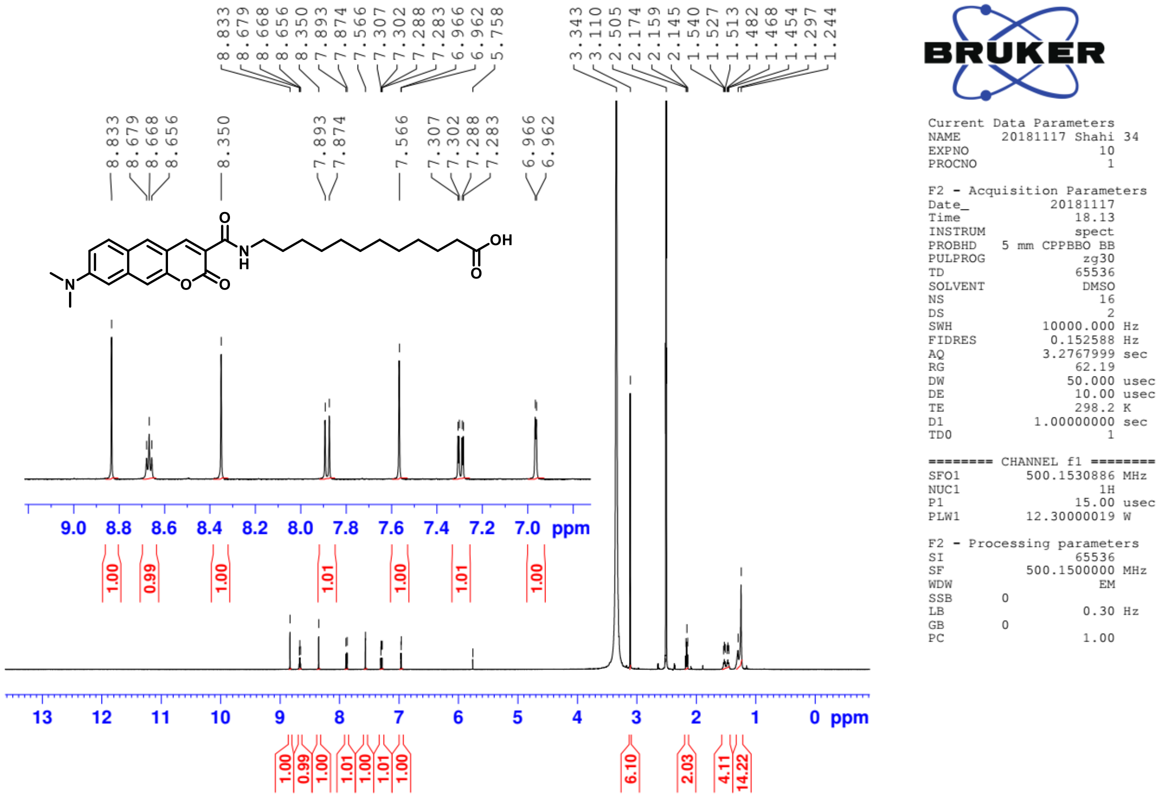

$^{1}H$ NMR spectrum of **DMABC-FA12** in 500 MHz (DMSO-$d_6$)

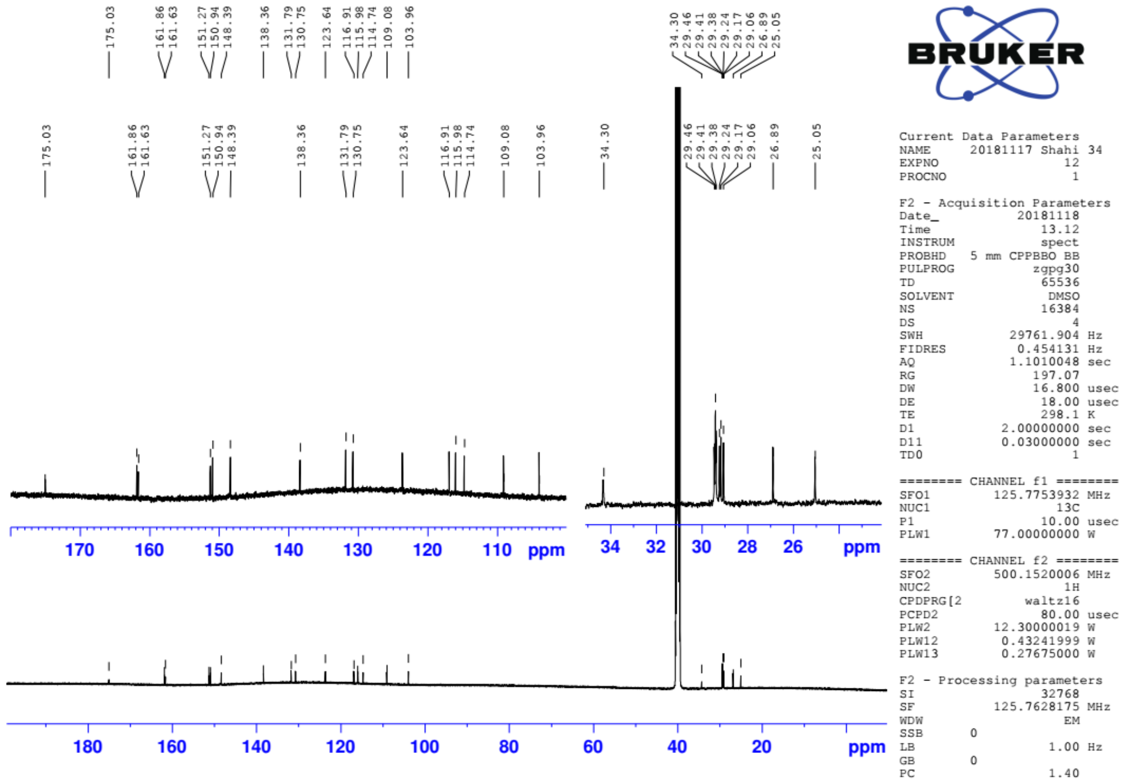

$^{13}C$ NMR spectrum of **DMABC-FA12** in 125 MHz (DMSO-$d_6$)

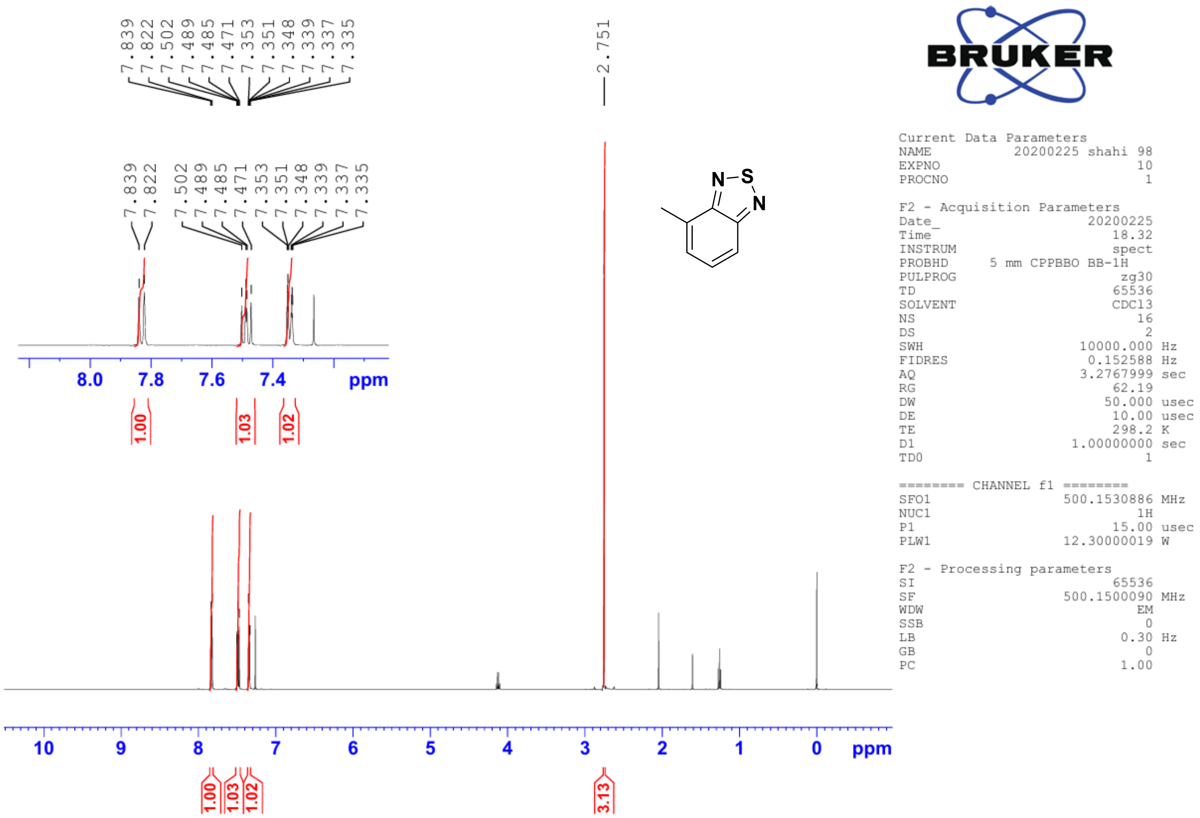


$^{1}$H NMR spectrum of compound **10** in 500 MHz ($CDCl_3$)

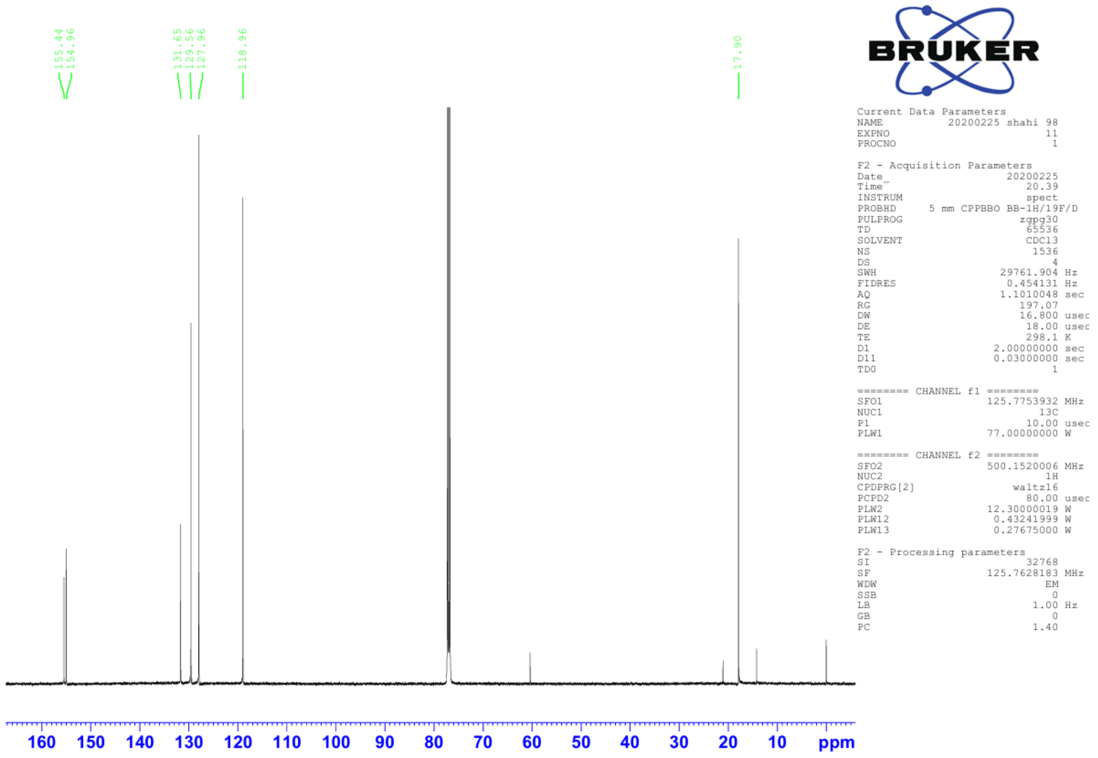


$^{13}$C NMR spectrum of compound **10** in 125 MHz ($CDCl_3$)

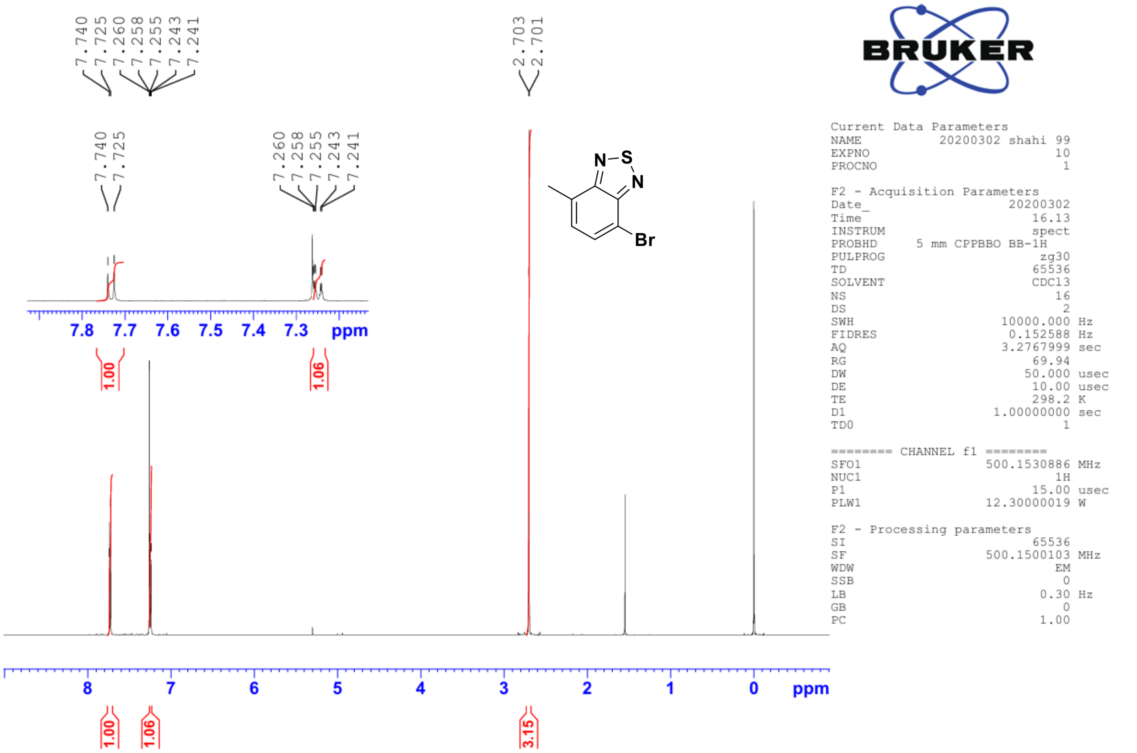


$^{1}$H NMR spectrum of compound **11** in 500 MHz ($CDCl_3$)

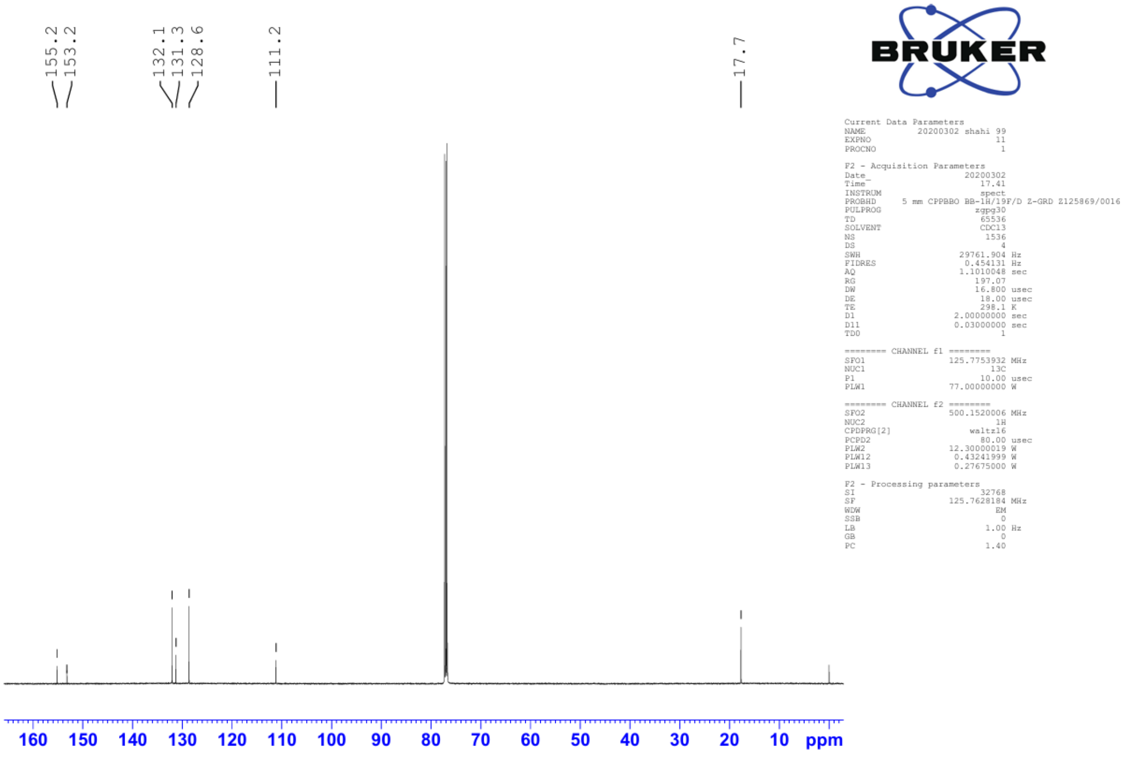


$^{13}$C NMR spectrum of compound **11** in 125 MHz ($CDCl_3$)

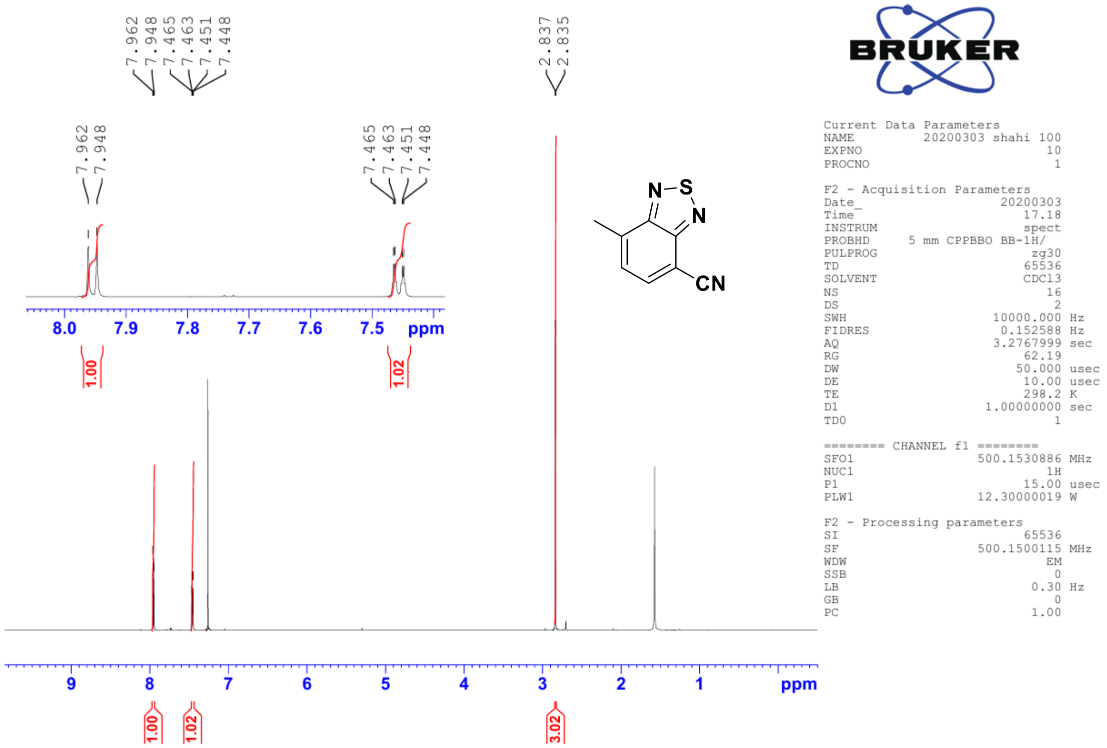


[1]H NMR spectrum of compound **12** in 500 MHz ($CDCl_3$)

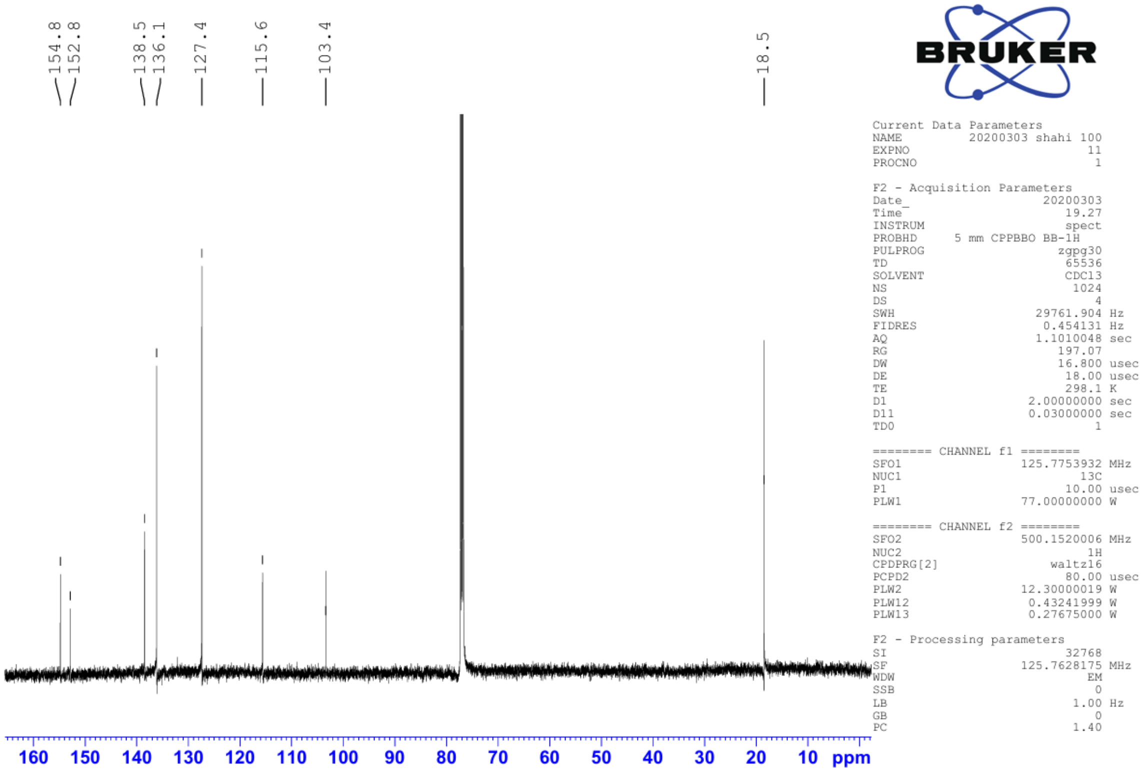


[13]C NMR spectrum of compound **12** in 125 MHz ($CDCl_3$)

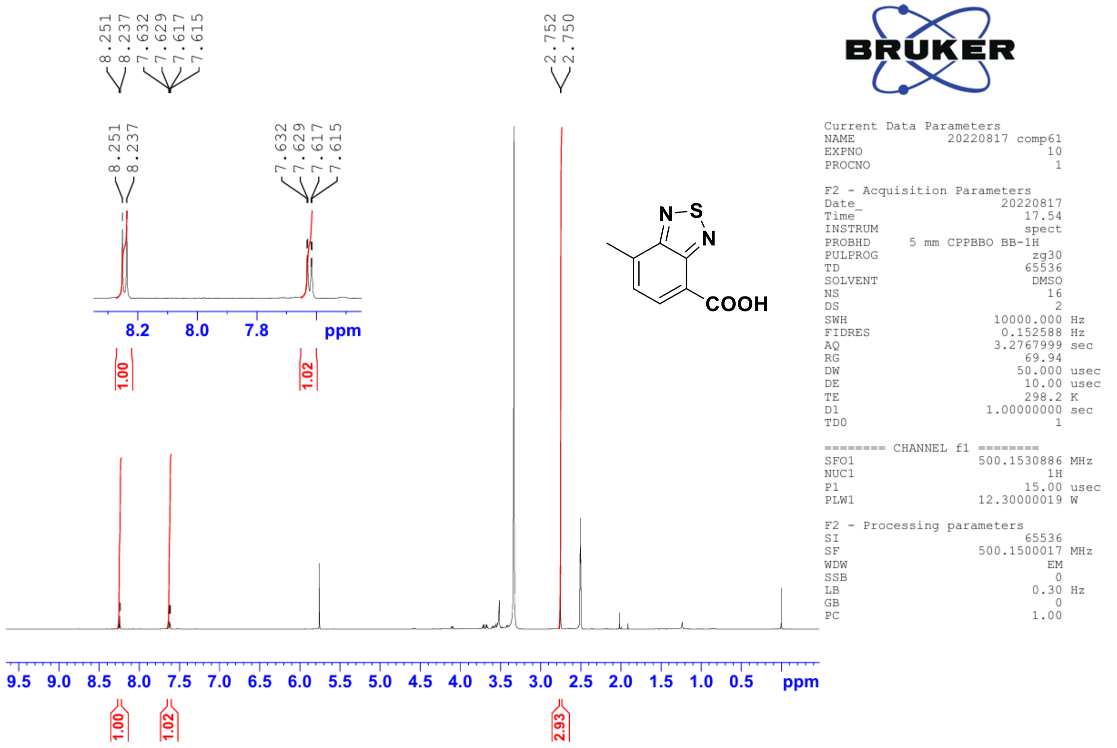


$^{1}H$ NMR spectrum of compound **13** in 500 MHz (DMSO-$d_6$)

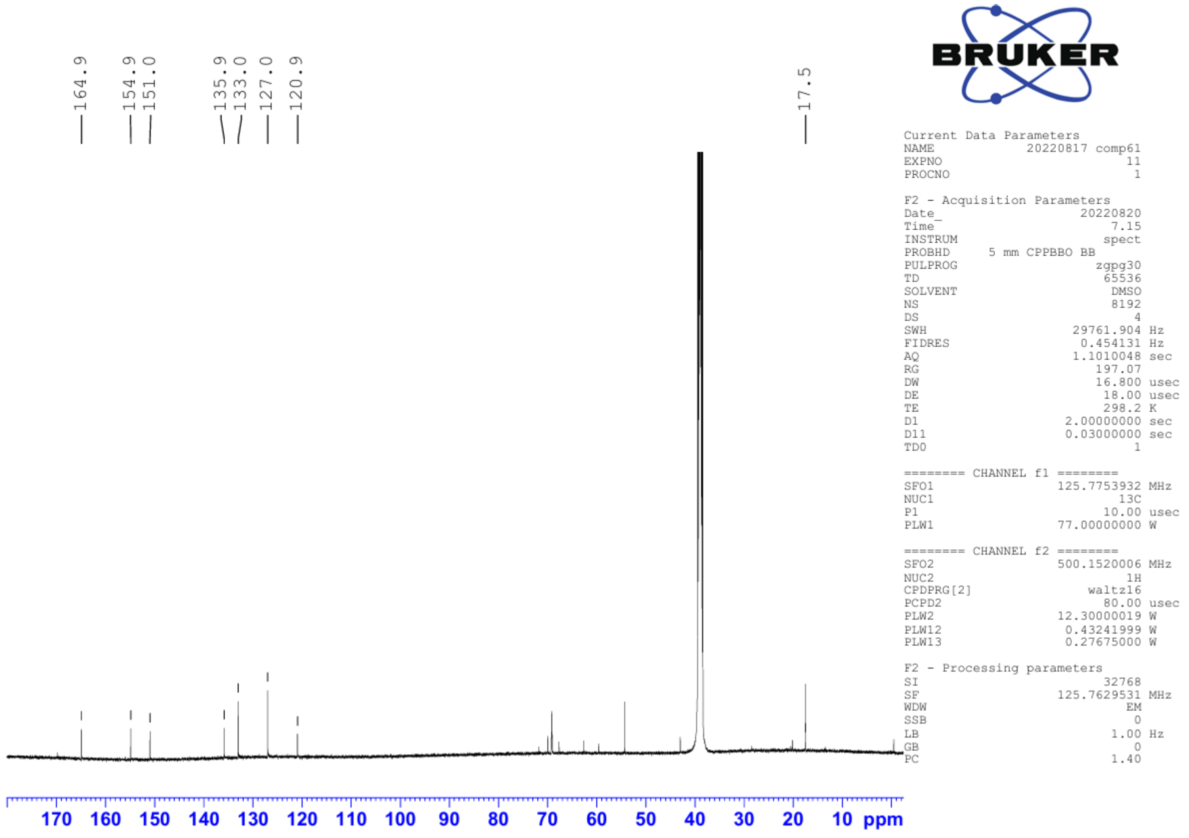


$^{13}C$ NMR spectrum of compound **13** in 125 MHz (DMSO-$d_6$)

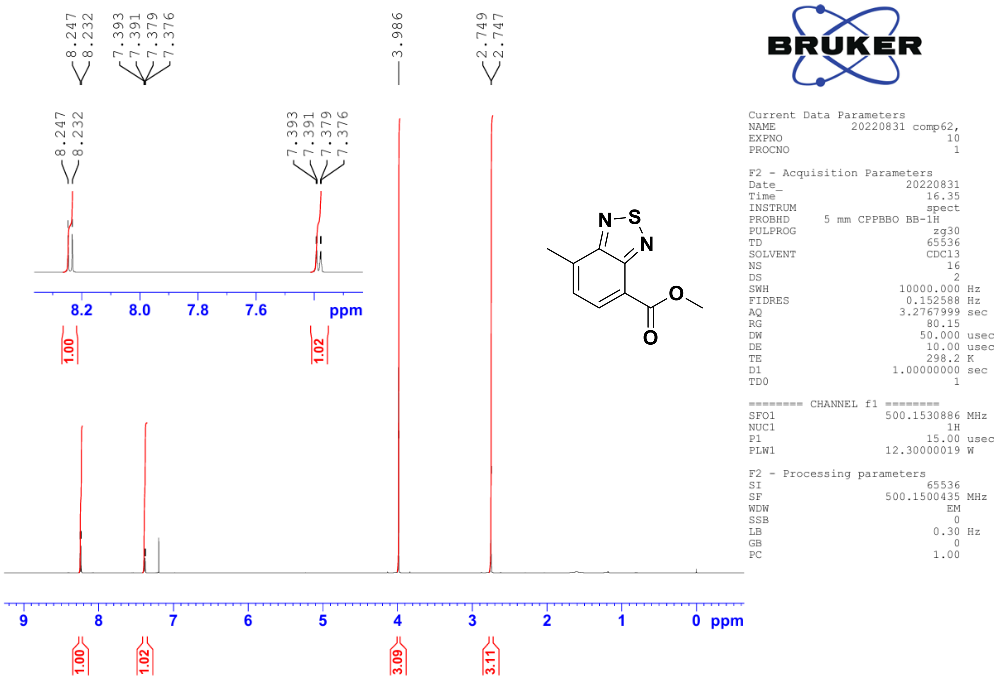

[1]H NMR spectrum of compound **14** in 500 MHz ($CDCl_3$)

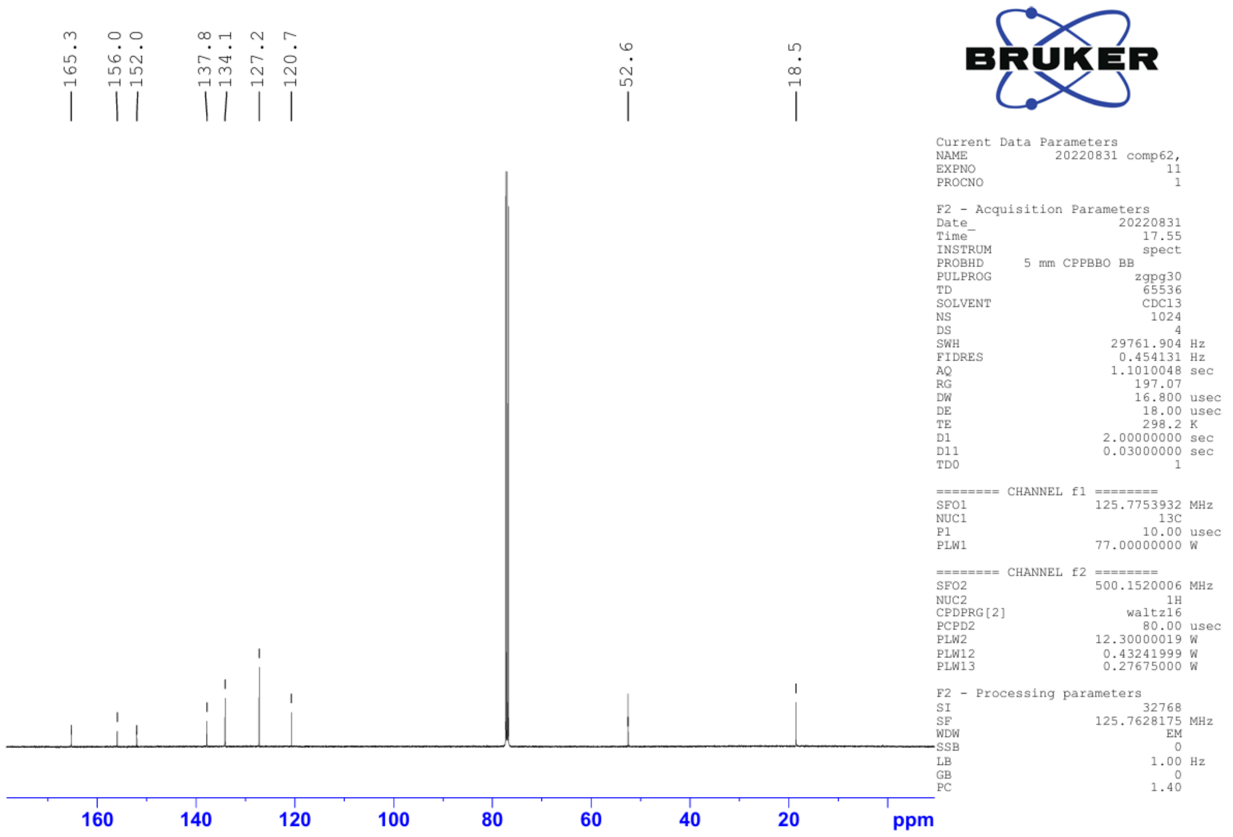

[13]C NMR spectrum of compound **14** in 125 MHz ($CDCl_3$)

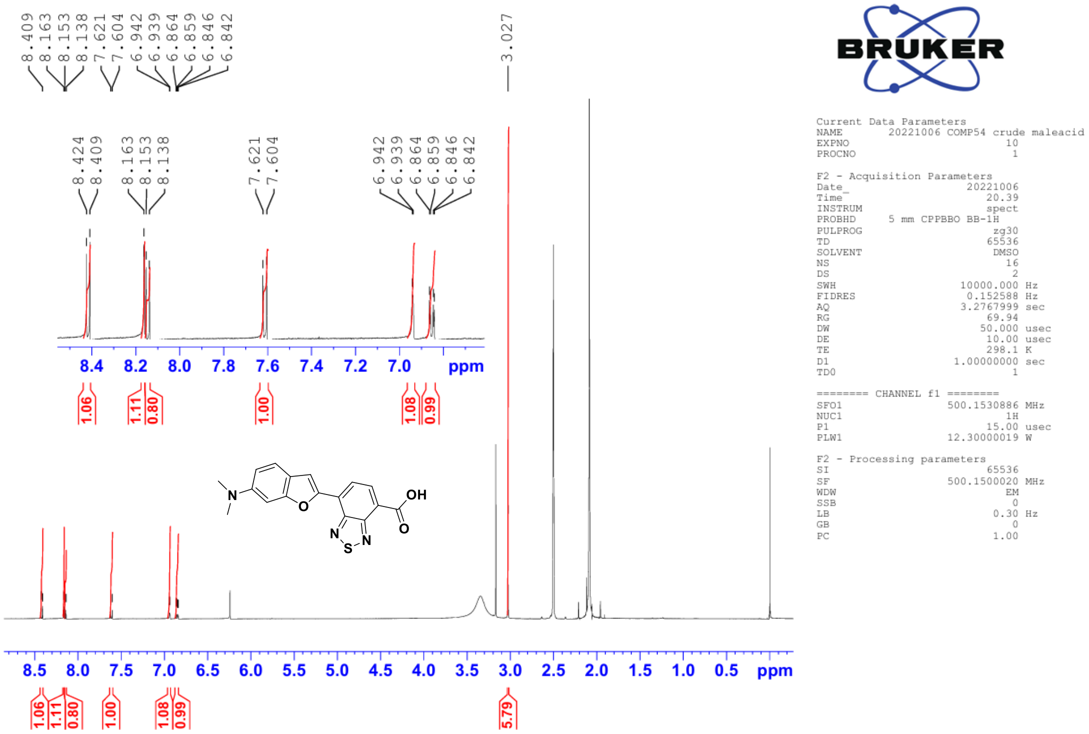


[1]H NMR spectrum of compound **16** in 500 MHz (DMSO-$d_6$)

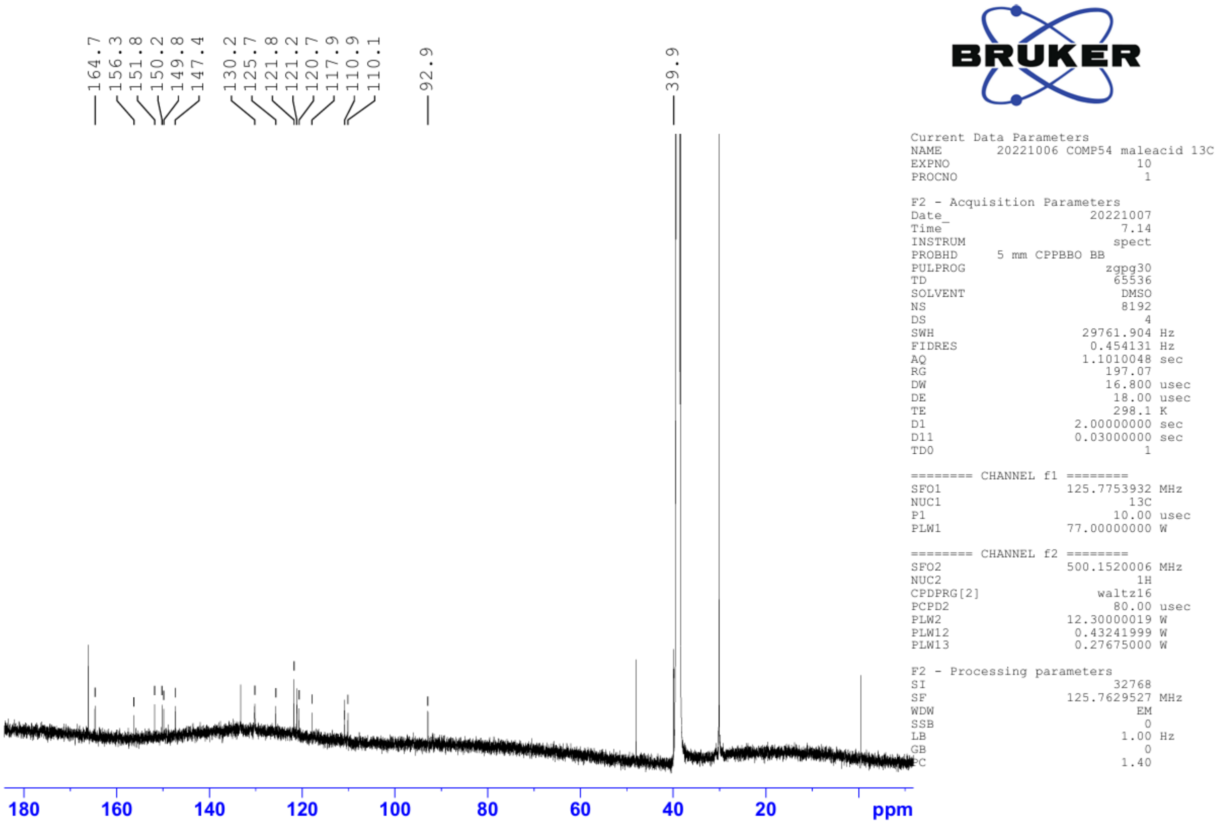


[13]C NMR spectrum of compound **16** in 125 MHz (DMSO-$d_6$)

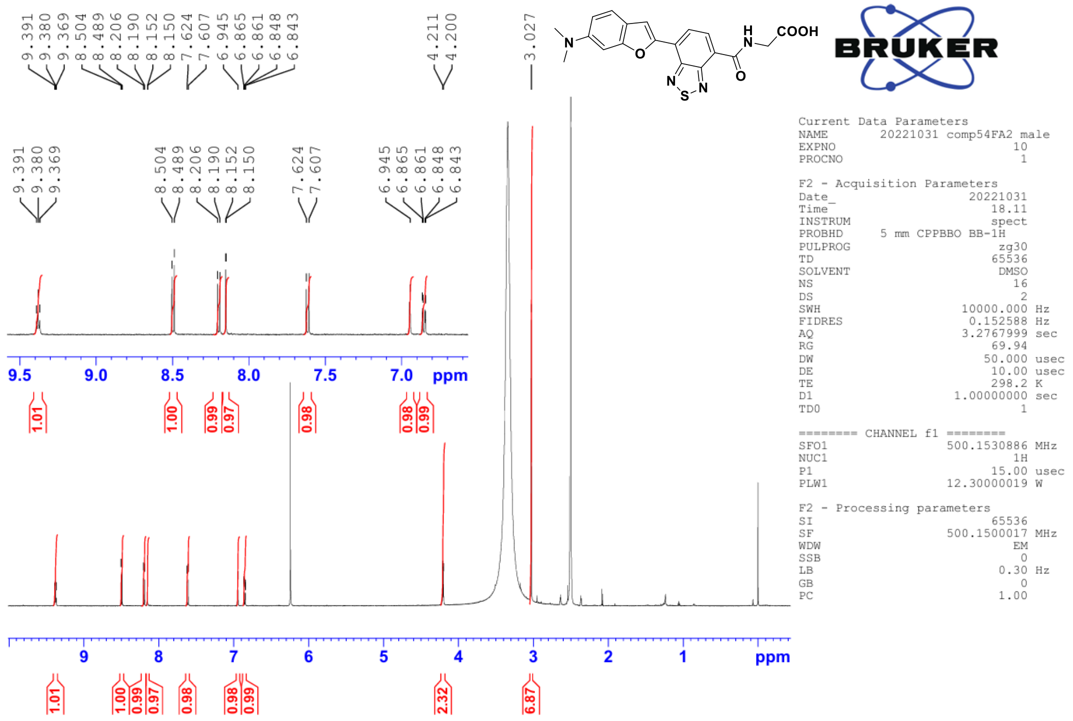

[1]H NMR spectrum of compound **DMABT-FA2** in 500 MHz (DMSO-$d_6$)

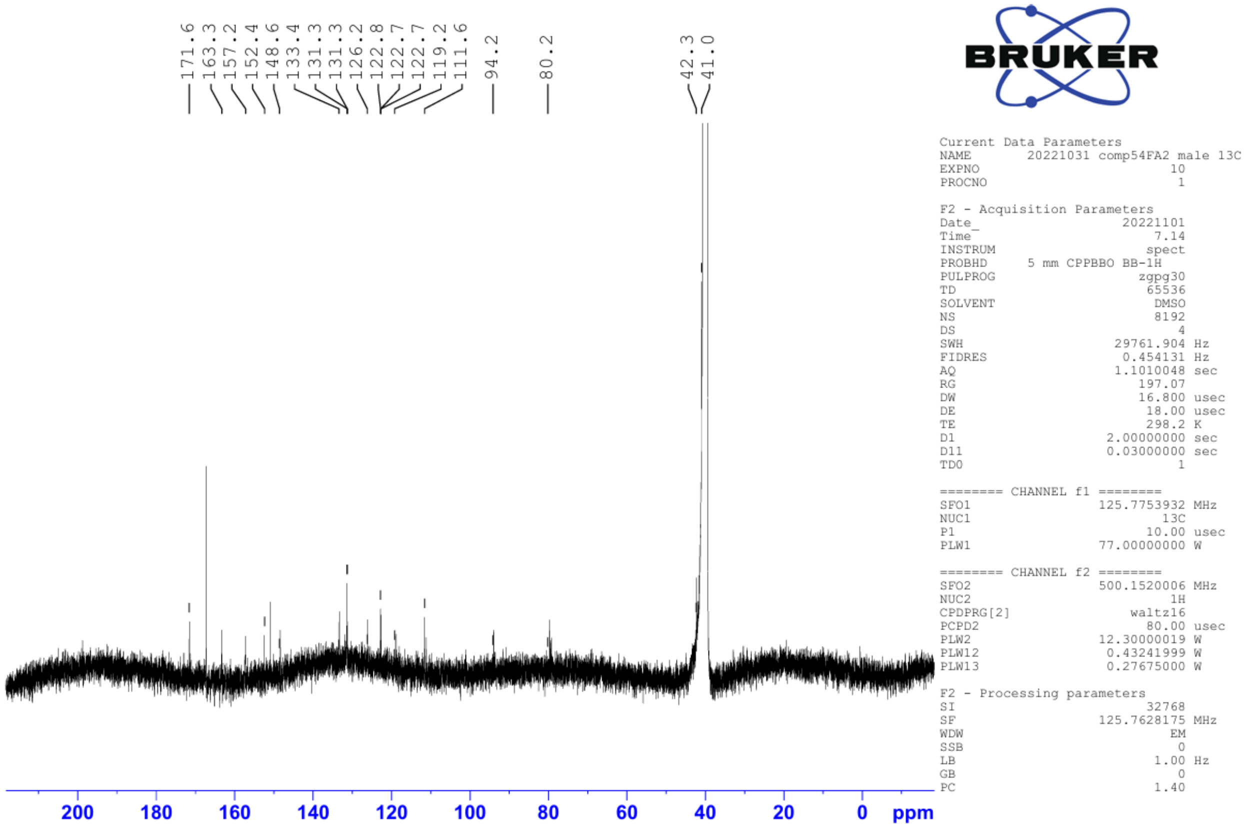

[13]C NMR spectrum of compound **DMABT-FA2** in 125 MHz (DMSO-$d_6$)

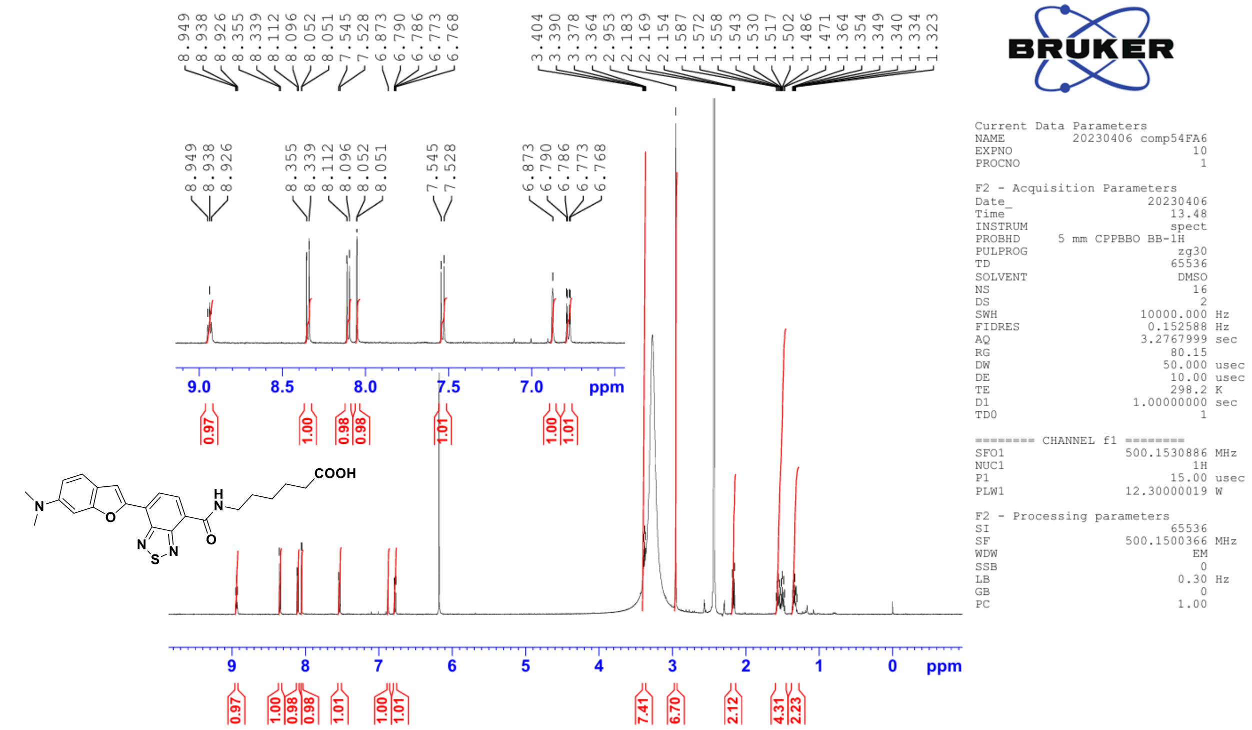


[1]H NMR spectrum of compound **DMABT-FA6** in 500 MHz (DMSO-$d_6$)

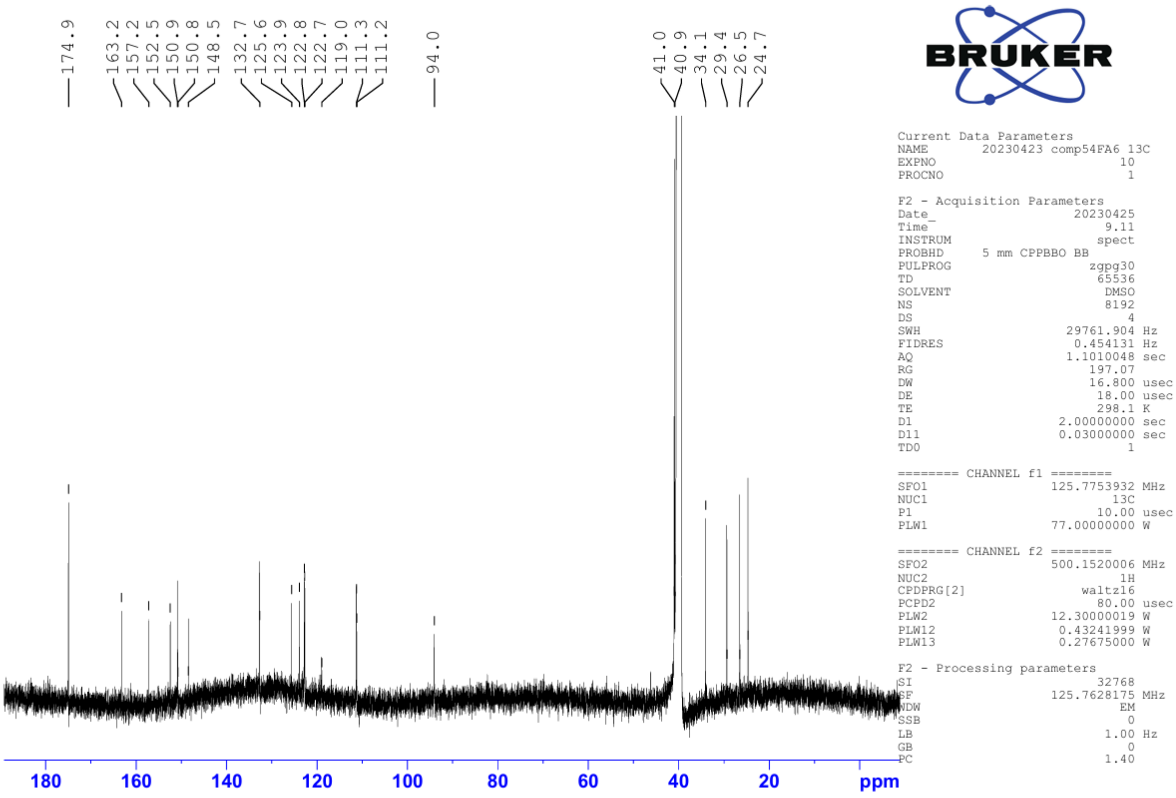


[13]C NMR spectrum of compound **DMABT-FA6** in 125 MHz (DMSO-$d_6$)

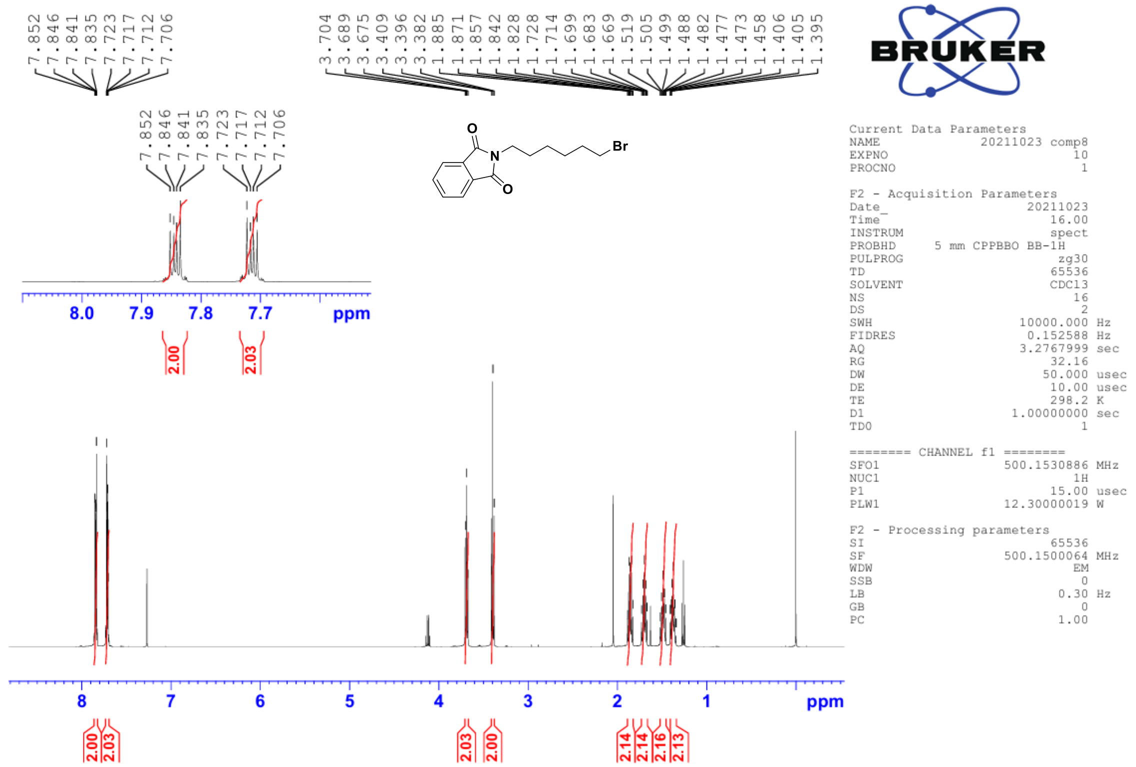


$^{1}H$ NMR spectrum of compound **17** in 500 MHz ($CDCl_3$)

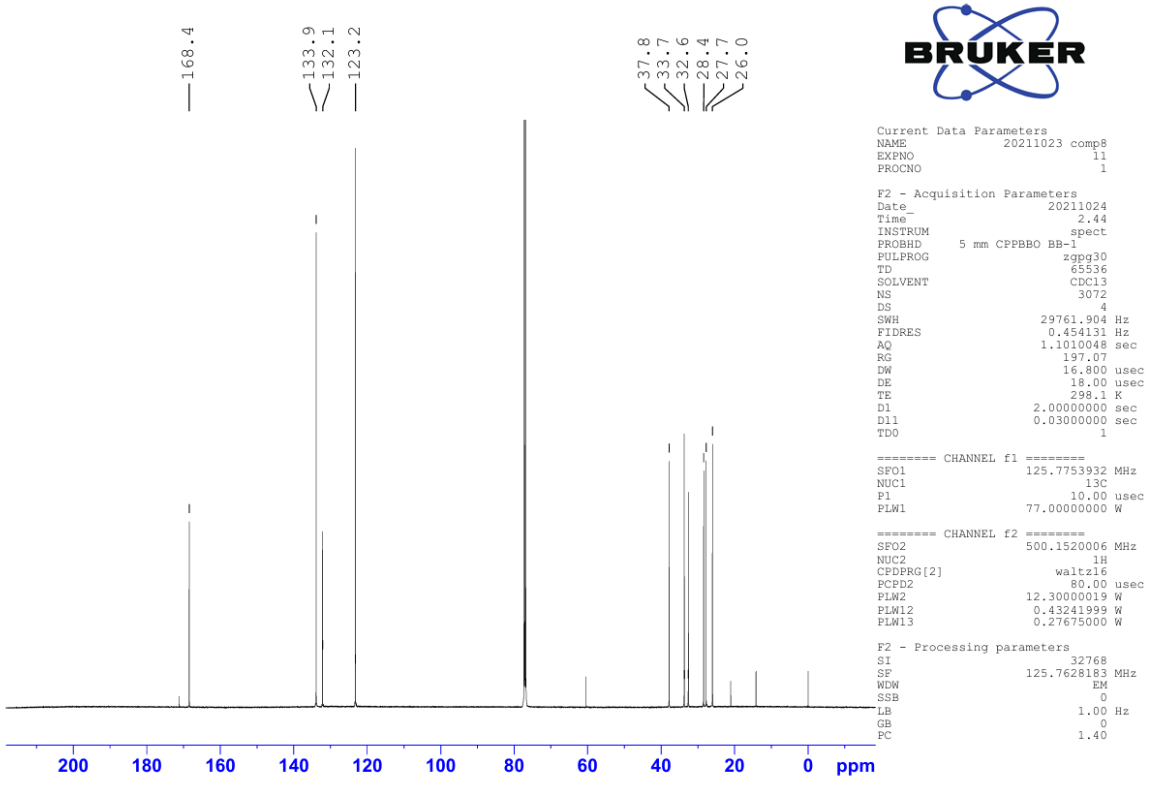


$^{13}C$ NMR spectrum of compound **17** in 125 MHz ($CDCl_3$)

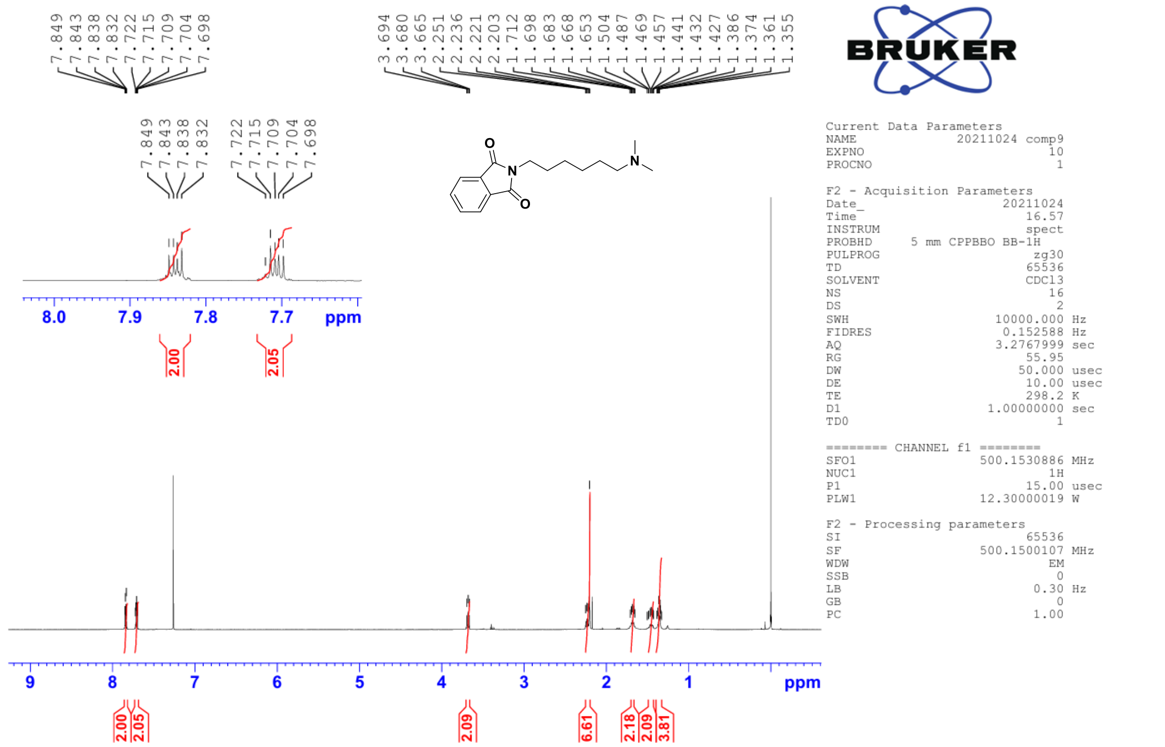

$^{1}$H NMR spectrum of compound **17** in 500 MHz ($CDCl_3$)

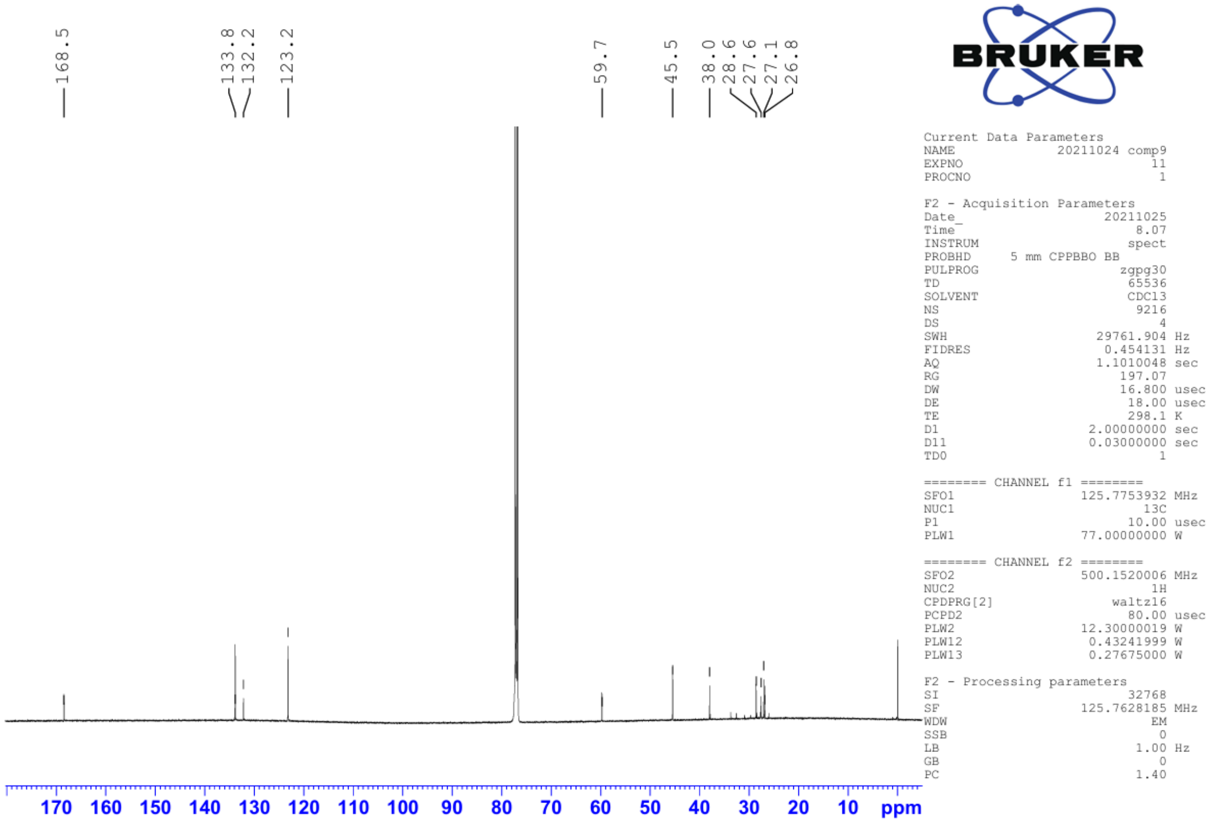

$^{13}$C NMR spectrum of compound **17** in 125 MHz ($CDCl_3$)

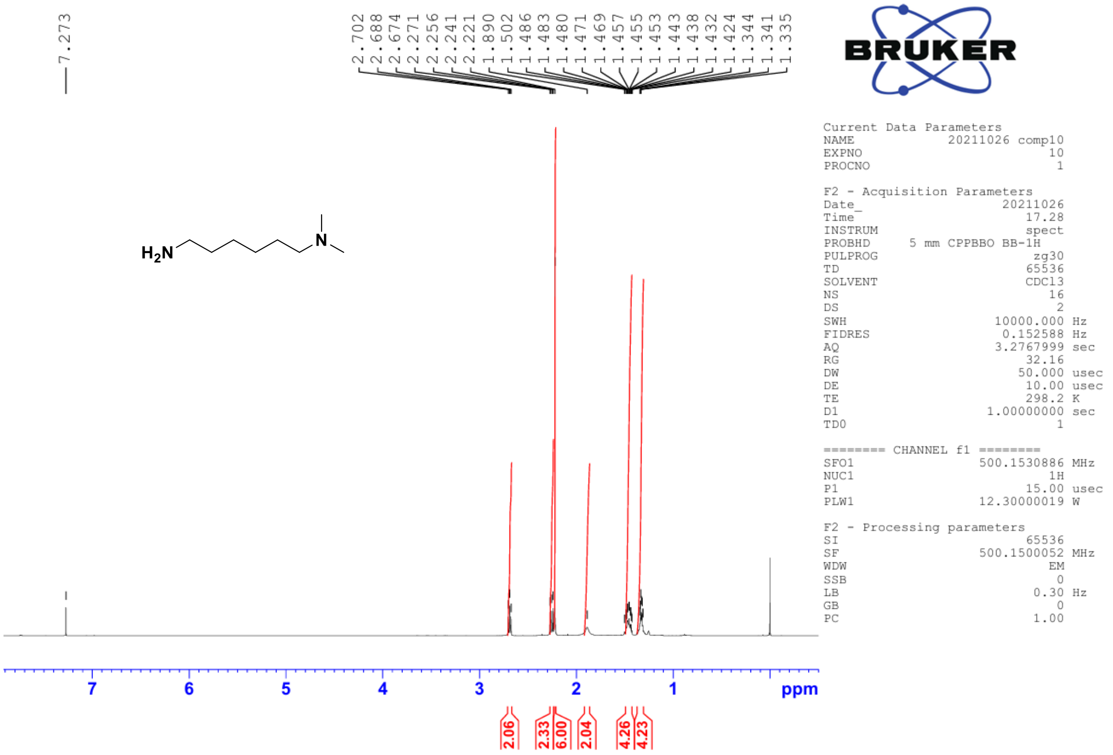

$^{1}H$ NMR spectrum of compound **18** in 500 MHz ($CDCl_3$)

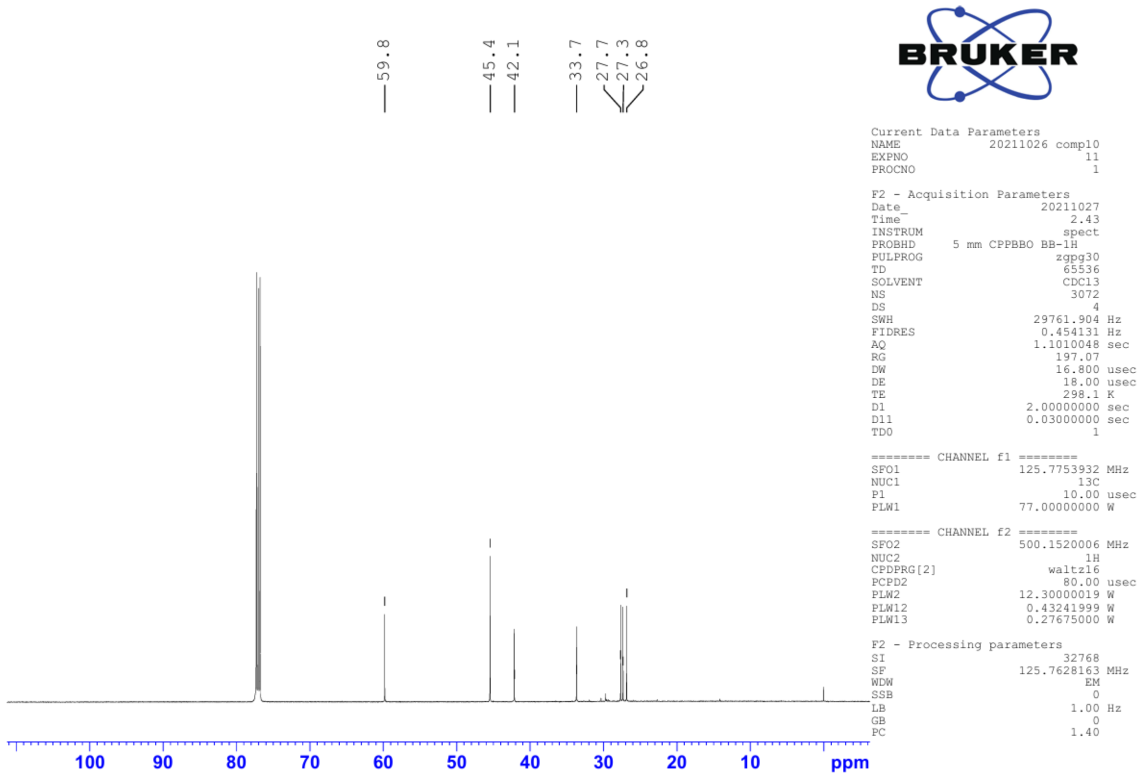

$^{13}C$ NMR spectrum of compound **18** in 125 MHz ($CDCl_3$)

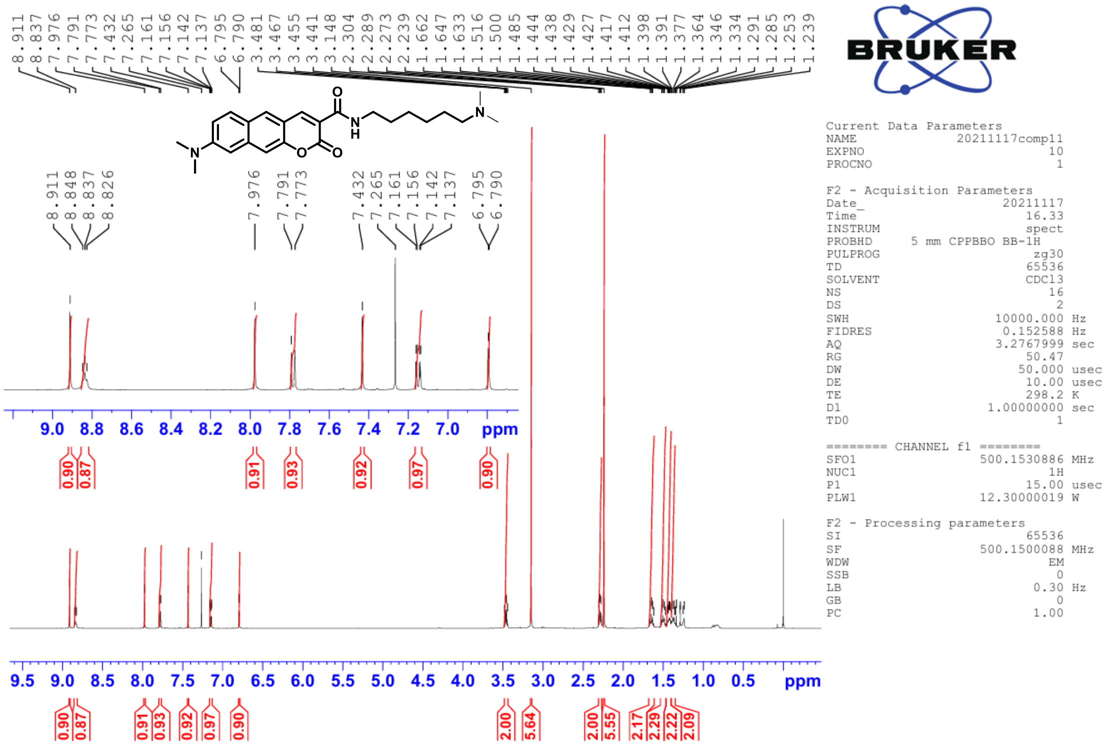


$^{1}H$ NMR spectrum of compound **DMABC-6DMA** in 500 MHz ($CDCl_3$)

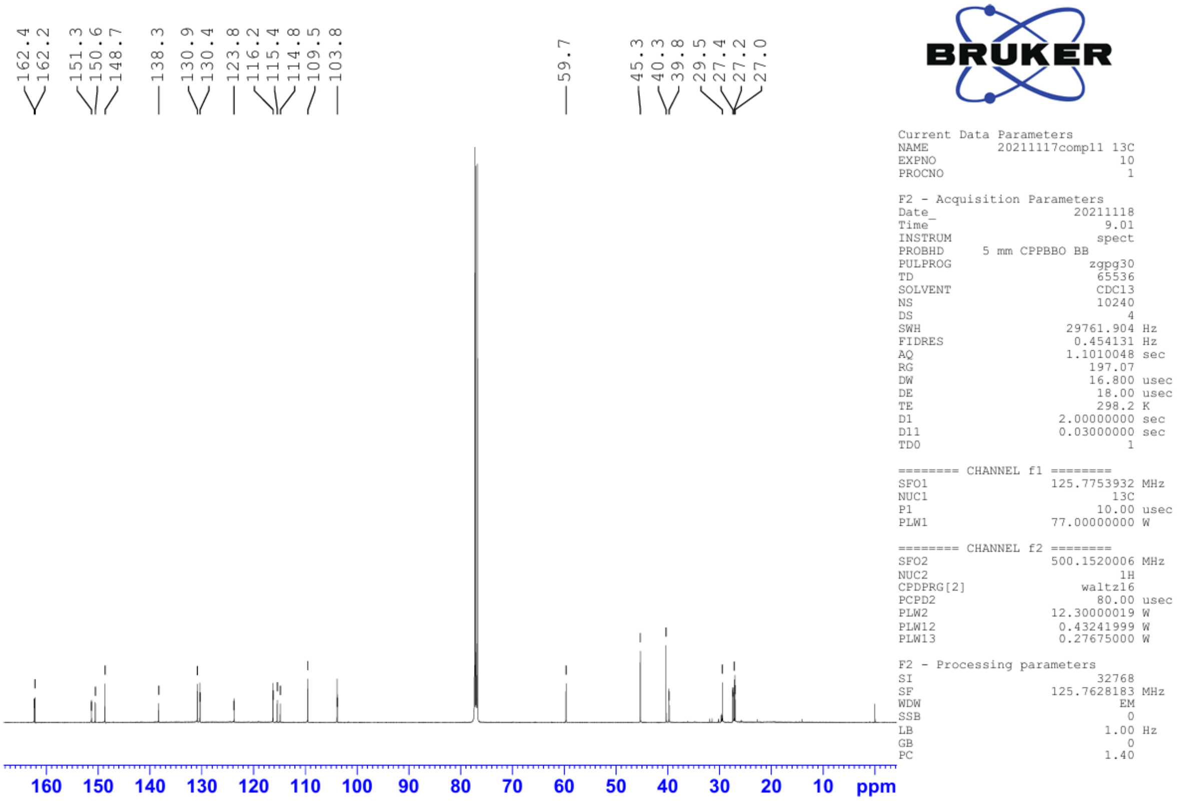


$^{13}C$ NMR spectrum of compound **DMABC-6DMA** in 125 MHz ($CDCl_3$)

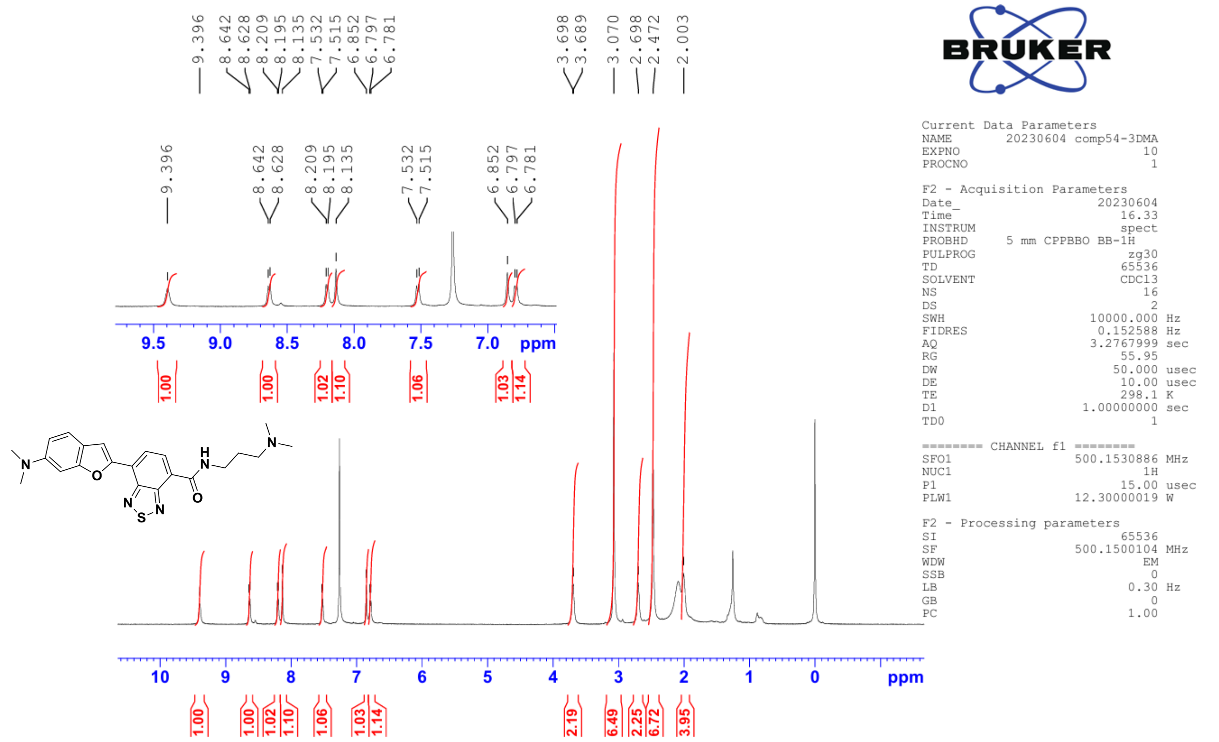


$^{1}H$ NMR spectrum of compound **DMABT-3DMA** in 500 MHz ($CDCl_3$)

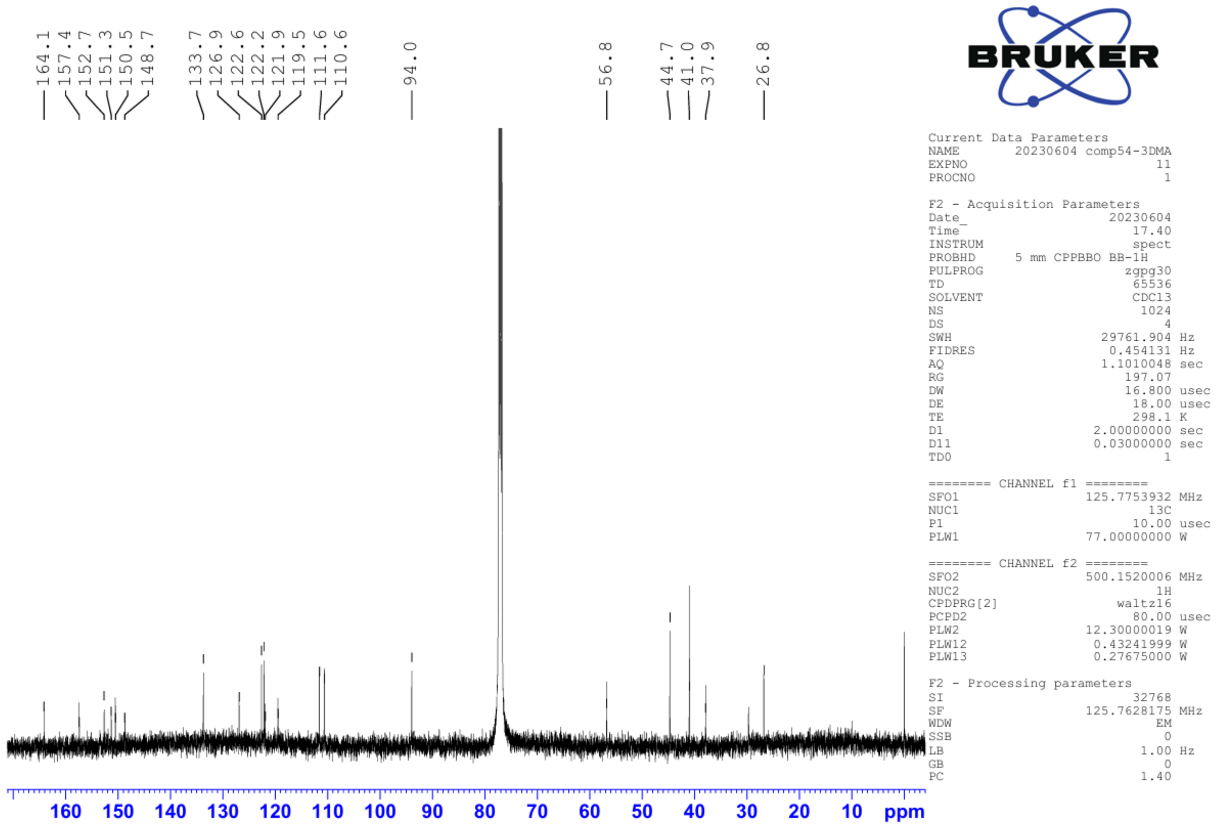


$^{13}C$ NMR spectrum of compound **DMABT-3DMA** in 125 MHz ($CDCl_3$)
